# Inhibition of Bacteriophage Infection by Natural and Synthetic DNA-binding Compounds

**DOI:** 10.64898/2026.11.23.768644

**Authors:** Vladimir Panteleev, Nika F. Petrova, Ivan Moldon, Alena I. Sidorova, Artem Isaev, Aleksandr Andriianov, Albert Arutyunyan, Andrey Moiseenko, Pavel N. Solyev, Andrey Kulbachinskiy, Daria Gelfenbein

## Abstract

The arms race between bacteria and phages involves a wide range of immune and anti-immune systems on both sides. One of the long-known mechanisms of antiphage defense involves DNA intercalators such as daunorubicin and its analogues. Daunorubicin was also shown to inhibit bacterial growth as well as eukaryotic cell division, underlying its use in chemotherapy. Other DNA intercalators and minor groove binding dyes are widely used for nucleic acids staining, but have not been studied for their effects on viral infection. Here, we systematically investigate the effects of known intercalator and DNA-binding agents, including daunorubicin and Hoechst derivatives, on phage infection in *Escherichia coli*. Using a diverse panel of phages with various replication mechanisms we identify phylogenetic groups sensitive to DNA-binding dyes and determined which stage of the viral life cycle was inhibited. Our data suggest that the active antiviral compounds block early steps of infection without cells killing, arguing against an abortive infection mechanism and lethal membrane permeabilization suggested recently. Together, our results expand the list of small molecules that affect phage infection and reveal DNA binders that inhibit viral DNA entry and/or replication without affecting cell physiology.

## Introduction

An arms race between bacteria and bacteriophages represents one of the most ancient biological conflicts on Earth. Among the diverse bacterial anti-viral defense strategies, production of small molecules has been recently re-established (1,2). These compounds can act as universal antiphage agents, particularly when secreted into the surrounding media (3). However, the first evidence of small molecules produced by Actinomycetes and *Aspergillus* sp., which were shown to inhibit coliphages, streptophages and staphylophages, was obtained in the first half of the 20th century (4,5). The subsequent decade witnessed extensive studies of Actinomycete-derived compounds (predominantly from *Streptomyces* species), including phagolessin A58, phagomycin, phagocidin, phagostin, aklavin, and rutilantin (6-10).

Igor Asheshov pioneered this work and postulated that studying antiphage activity of natural compounds in a relatively simple system could be extrapolated to help identify chemicals active against dangerous human viruses (11). This hypothesis proved to be at least partially correct, as demonstrated for daunorubicin, which was later shown to inhibit both hepatitis B virus and HIV (12,13). The chemical structure of rutilantin, described in 1961, is close to that of daunorubicin. Daunorubicin, an anthracycline antibiotic with antitumor activity, is able to intercalate DNA and has become one of the most widely studied examples of antiphage agents produced by *Streptomyces*. It exhibits antiphage activity against both double-stranded and single-stranded DNA viruses and has been well known as an antiphage compound for over 60 years (14,15). In addition to acting against virulent phages, it can also induce prophages and eliminate prophages, likely due to its DNA-damaging action (16,17).

Daunorubicin-like compounds are commonly produced by *Streptomyces* and may represent a basal antiphage defense system. Extracts of 14 out of 48 investigated *Streptomyces* strains contained daunorubicin-like compounds, suppressing phage infection in *Streptomyces coelicolor, Escherichia coli* and *Pseudomonas aeruginosa* systems, without affecting bacterial growth. Although the mechanism of anti-phage activity was not determined, these compounds acted at early steps of the infection cycle (2). It was proposed that the linear dsDNA of the phage is more sensitive to intercalators compared to the bacterial supercoiled genome. However, this mechanism cannot be universal, as daunorubicin also inhibits the ssDNA phage φX174 (14). Alternative mechanisms have been proposed, including enhanced susceptibility of intercalated DNA to restriction enzymes, defects in DNA circularization, or direct damage via hydroxyl radicals (18). Nevertheless, these hypotheses fail to account for the observed phage specificity.

More recently, other small molecules, such as synthetic benzimidazolylpyrazoles and aminoglycosides were shown to inhibit both ssDNA and dsDNA phages, acting at early post-adsorption steps (19). These compounds do not intercalate DNA, raising the possibility that intercalating agents may also inhibit phage infection through mechanisms other than direct binding to viral DNA.

In support of this idea, it was recently demonstrated that both intercalating (daunorubicin) and non-intercalating (kanamycin) compounds block viral DNA injection after the adsorption step, an effect attributed to membrane permeabilization leading to growth arrest (20). In particular, it was shown that daunorubicin accumulates in the cell during the early steps of phage entry, reaching concentrations that lead to abortive infection (20). Transcriptomic and proteomic analyses of daunorubicin-mediated infection inhibition further suggested an abortive infection mechanism of protection (1). Additionally, a recent study demonstrated synergy between small antiphage molecules and restriction-modification systems, suggesting that these small molecules may potentiate other types of immunity (1).

While daunorubicin is a well-studied compound in the context of phage suppression, only sporadic data exist on the antiphage effects of other DNA-binding chemicals, such as DAPI and propidium iodide (PI). A previous screen of nearly 5,000 natural and synthetic molecules in *Escherichia coli* identified 11 compounds with antiphage activities, 9 of which belong to intercalators (2). However, these reports remain scarce, as the primary role of most intercalating agents and DNA-binders – such as Hoechst dyes, acridine orange, and various cyanine and monomeric cyanine dyes – has been limited to nucleic acid detection and cell staining.

In this study, we performed comparative analysis of the antiphage activity of DNA-binding dyes widely used for live-cell staining (e.g., various Hoechst derivatives). These dyes bind to the minor groove of DNA, preferentially at A/T-rich sequences, and are generally less toxic than DAPI (21). Despite their well-characterized DNA-binding properties, the potential of these dyes to modulate phage infection has never been investigated. Here, we identified widely used synthetic DNA dye Hoechst 33258 as the strong phage infection inhibitor. The results demonstrate a novel role for DNA-binding agents in modulating phage infection and provide a foundation for future studies exploring the antiphage properties of other small chemicals, both synthetic and natural. Our data also highlight that DNA staining compounds commonly used for visualization of phage infection and bacterial anti-phage immunity phenotypes, may directly affect experimental outcomes by interfering with phage infection.

## Materials and methods

### Synthetic routes to obtain new bisbenzimidazoles

Synthesis of new dimeric bisbenzimidazoles with the 2-aminoimidazole fragment RDB_2_(3) and RDB_2_ (7) included 7 steps (Scheme 1, Supplementary materials, Section 2). First, we reduced the nitro group of I in the excess of hydrazine. The diamine II was cyclized into the carbamate III by the condensation with carbonyldiimidazole, and later converted into the chloro derivative by boiling with phosphorus oxychloride to produce IV. Then we substituted the chloride with N-methylpiperazine, saturated V with dry hydrochloric acid and used VI for the condensation with diaminobenzoic acid. Compounds RDB_2_ (3) and RDB_2_ (7) were obtained by the reaction with the corresponding diamines through the active ester formation. For the synthesis of DB_2_Py we coupled X with pyridinedicarboxylic acid using HCTU as a condensing agent. For the synthesis of MB_2_Cl we saturated IV with dry hydrochloric acid to obtain the imino ester XII and added XIV immediately after reducing the nitro group of XIII in a flow of hydrogen. Detailed analysis of obtained compounds including MALDI MS, NMR, HRMS spectra is provided in Supplementary materials, Section 2.

### Strains and vectors

*Escherichia coli* K-12 MG1655 was used as standard strain in all experiments except microscopy. A plasmid for phage DNA visualization (pIM11) expressed seqA_G4S_EGFP under arabinose promoter in pBAD vector was constructed using Gibson Assembly. *E. coli* K-12 MG1655 dam– strain (EC *dam*) was obtained via P1 transduction (22) from donor dam::Tn9 (CmR) GM2929 (*E. coli* Genetic Stock Center) and recipient *E. coli* K-12 MG1655 strains. Phages were obtained from the laboratory collection, Basel collection (Bas), Hong Kong collection (HK), Moscow collection (MSK), provided by Artem Isaev.

### *E. coli* growth in the presence of DNA dyes

To measure possible toxicity of DNA dyes, *Escherichia coli* K-12 MG1655 was grown overnight, diluted 1:50 in fresh LB medium, and added to 96-well plates containing twofold serial dilutions of the indicated chemicals, with a final volume of 100 µL per well. Growth was monitored using either a CLARIOstar Plus plate reader (BMG LABTECH) or an Infinite 200 PRO plate reader (Tecan). In both instruments, cultures were incubated at 30 °C with shaking at 200 rpm. Optical density at 600 nm (OD600) was recorded every 10 min in the CLARIOstar Plus and every 15 min in the Infinite 200 PRO (Tecan). MICs were calculated from growth curves and used for selection of optimal working concentration for each compound (see Figure S2). If not indicated in the assay description, working concentration was used in the experiment.

### Phage lysis assay in liquid culture

To test the effect of the chemicals on phage infectivity in liquid culture, *E. coli* K-12 MG1655 was grown overnight, diluted 1:50 in fresh LB medium supplemented with 5 mM CaCl_2_, 10 mM MgCl_2_ (LBphage medium), studied compounds, and aliquoted into 96-well plates at 100 µL per well. Phage suspensions were added in 20 µL volumes at the multiplicity of infection (MOI) indicated for each experiment, using 10-fold serial dilutions. Growth was monitored using either a CLARIOstar Plus plate reader (BMG LABTECH), an Infinite 200 PRO (Tecan) or FlexA-200 (Allsheng) plate reader under the conditions described above. Of note, for Qbeta RNA phage, *E. coli* NEB Turbo (F’ proA+B+ lacIq ΔlacZM15 / fhuA2 Δ(lac-proAB) glnV galK16 galE15 R(zgb-210::Tn10)TetS endA1 thi-1 Δ(hsdS-mcrB)5) strain was used.

### Efficiency Of Plating assay

To test the effect of the compounds on plaque formation efficiency, an overnight culture of *E. coli* K-12 MG1655 was diluted 1:100 in LB medium and incubated at 37 °C with shaking at 200 rpm until the culture reached an OD600 of 1.0. Then 1.5 ml of culture was mixed with 30 mL of molten 0.5% LB top agar maintained at 45 °C and supplemented with 10 mM MgCl_2_, 5 mM CaCl_2_ (LBphage agar), and studied compounds in working concentration. The mixture was poured onto 12x12-cm square plates containing 15 mL of solid 1.5% LB agar and allowed to solidify. Next, 3 µL of 10-fold serial dilutions of phage suspensions were spotted onto the surface of the solidified top agar and allowed to dry. Plates were incubated for 14 h at 30 °C and imaged using an Ingenius LHR2 Gel Imaging System (Syngene).

### Influence of the timing of Hoechst 33258 addition on phage infection

To determine whether the timing of Hoechst 33258 addition affected phage infection, phages T5, P1, and Lambda at indicated MOI were preincubated with 500 µL of *E. coli* K-12 MG1655 culture adjusted to OD600 = 0.5 for 30 min. The following conditions were compared: (i) no Hoechst 33258; (ii) Hoechst 33258 present during preincubation and maintained throughout the experiment without washing; (iii) Hoechst 33258 added before phage inoculation and removed by washing after 30-min adsorption time; and (iv) Hoechst 33258 added after adsorption. After 30 min of incubation with phage, 100 µL aliquots were transferred to 96-well plates for growth curve measurements in the Infinite 200 PRO reader (Tecan).

### Effect of chemical pretreatment of phages on infectivity

To test if phages might be inactivated by DNA dyes, T5, P1, and Lambda phage suspensions were pretreated with Hoechst 33258 at the indicated working concentration for 1 h at 37 °C, followed by overnight incubation at 4 °C in the dark. Phage infectivity was subsequently assessed using the growth curve assay described above. Briefly, *E. coli* K-12 MG1655 was grown overnight, diluted 1:50 in fresh LBphage medium, and aliquoted into 96-well plates at 100 µL per well. Hoechst 33258-treated and non-treated control phages were added in 20 µL volumes at the indicated multiplicity of infection (MOI). Growth was monitored using Infinite 200 PRO plate reader (Tecan) under the conditions described above. Controls included LB supplemented with working concentration of Hoechst 33258.

In addition to growth curve analysis, phage infectivity was assessed using the efficiency of plating (EOP) assay as described above using phages that were pre-soaked in solution containing or lacking DNA binders. Control plates contained Hoechst 33258 at the working concentration and no DNA binder was added to the experimental plates.

### Measurement of phage elimination and adsorption efficiencies

To measure phage elimination and adsorption efficiencies, an overnight culture of *E. coli* K-12 MG1655 was diluted 1:100 in LBphage medium, and incubated at 37 °C with shaking at 200 rpm until the culture reached an OD600 of 1.0. Then, 500 µL of the resulting culture with or without compounds was infected with 20 µL of phage dilution (∼700 plaque-forming units [PFU] in total), and adsorption was allowed to proceed for 20 min at 30 °C. Cells and adsorbed phages were pelleted at 4602 × g for 3 min in an Eppendorf 5424 centrifuge. Next, 400 µL of the supernatant was collected for plaque assay (the “Supernatant” sample), while the remaining pellet was washed with 1 mL of fresh LB medium (the “Pellet” sample). In parallel, the same procedure was conducted without centrifugation, and the whole sample was taken for plaque assay (the “Total” sample). For the plaque assay, the entire volumes of the mixtures were mixed with 10 mL of LBphage agar. For the “Supernatant” sample, an additional 500 µL of fresh bacteria at OD600 = 1.0 was added. The mixtures were poured onto 9-cm Petri dishes containing 5 mL of solid 1.5% LB agar and allowed to solidify. Plates were incubated for 14 h at 30 °C and imaged using an Ingenius LHR2 Gel Imaging System (Syngene).

### Efficiency of Center Of Infection assay

To measure the efficiency of phage replication in initially infected cells, an overnight culture of *E. coli* K-12 MG1655 was diluted 1:100 in LBphage medium, and incubated at 37 °C with shaking at 200 rpm until the culture reached an OD600 of 1.0. Then, 100 µL of the resulting culture with or without chemicals was infected with 20 µL of phage dilution (∼700 plaque-forming units [PFU] in total), and adsorption was allowed to proceed for 20 min at 30 °C. Cells were washed twice with 1 ml of fresh LB medium and using pelleting at 4602 × g for 3 min in an Eppendorf 5424 centrifuge. The resulting pellet was supplemented with 400 µl of fresh bacteria of OD600 = 1 and mixed with 10 mL of LBphage agar. The mixtures were poured onto 9-cm Petri dishes containing 5 mL of solid 1.5% LB agar and allowed to solidify. Plates were incubated for 14 h at 30 °C and imaged using an Ingenius LHR2 Gel Imaging System (Syngene).

### Dye fluorescence measurement

To measure intrinsic chemical fluorescence changes during phage infection, an overnight culture of *Escherichia coli* K-12 MG1655 was diluted 1:100 in LB medium and incubated at 37 °C with shaking at 200 rpm until the culture reached an OD600 of 1.0. The culture was then resuspended in LBphage medium with chemicals in working concentration, and SYTO 9 to a final OD600 of 0.5. 100 µL aliquots were transferred to black 96-well plates with transparent bottoms (Corning). Phage suspensions were added in 20 µL volumes at MOI indicated for each experiment, using 10-fold serial dilutions. Optical density at 600 nm (OD_600_) and fluorescence were measured using a CLARIOstar Plus plate reader (BMG LABTECH). Fluorescence was recorded using the following settings: Hoechst 33258, excitation at 355 ± 20 nm, a 402.5-nm dichroic filter, and emission detection at 455 ± 30 nm; SYTO9, excitation at 470 ± 10 nm, a 490-nm dichroic filter, and emission detection at 510 ± 10 nm; and daunorubicin, excitation at 470 ± 10 nm, a 532.5-nm dichroic filter, and emission detection at 600 ± 20 nm.

### Quantitative PCR experiment for measuring phage replication and transcription

To quantify the effect of Hoechst 33258 presence on relative amounts of phage and bacterial DNA in cells and transcription of early and late phage genes, an overnight culture of *E. coli* K-12 MG1655 was diluted 1:100 in LB medium and grown at 37 °C with shaking at 200 rpm until the culture reached an OD600 of 1. Cells were harvested by centrifugation in an Eppendorf 5424 centrifuge at 7,000 rpm for 3 min and resuspended in LBphage medium to a final OD600 of 2. Lambda phage was added at MOIs of 0.3, in the presence of 25 µM Hoechst 33258. Following adsorption for 15 min at room temperature, the cells were pelleted by centrifugation and diluted to an OD600 of 1 with a twofold volume of fresh LB medium containing 25 µM Hoechst 33258. The cell suspension was aliquoted into a 24-well plate at 1 ml per well and incubated at 30 °C with shaking at 200 rpm. At 0, 15, 45, and 75 min after the adsorption, two 450 µl samples were collected into 1.5 ml tubes and placed on ice. To remove free phage particles, cells were pelleted by centrifugation in an Eppendorf 5424 centrifuge at 7,000 rpm for 3 min, the supernatant was removed, and the pellets were resuspended in 1 ml of cold LB medium. The cells were centrifuged again under the same conditions, the supernatant was discarded, and the resulting pellets were snap-frozen in liquid nitrogen and stored at −80°C.

Primers for qPCR assays targeting bacterial and phage genomes and certain genes were designed using Primer-BLAST (23). The absence of primer dimers was verified using the Thermo Fisher Scientific Multiple Primer Analyzer. The efficiency of each primer pair was determined by standard-curve analysis, and amplicon specificity was assessed by melting-curve analysis. qPCR was performed using qPCRmix-HS SYBR Green I premix (Evrogen, Russia) on a Bio-Rad CFX96 Touch Real-Time PCR Detection System. Reactions were carried out in a final volume of 15 µl under the following cycling conditions: initial denaturation at 95°C for 3 min; followed by 40 cycles of 95°C for 10 s, 57°C for 15 s, and 72°C for 15 s. Ct values were automatically determined using the threshold method in the CFX96 software.

To measure the expression levels of phage genes, RNA was purified from cell pellets using the GeneJET RNA Purification Kit (Thermo Scientific) according to the manufacturer’s instructions. Residual DNA was removed from the purified samples by treatment with RNase-free DNase I (New England Biolabs) according to the manufacturer’s instructions. cDNA was synthesized using random hexamer primers (Thermo Scientific) and MMLV reverse transcriptase (Evrogen, Russia) according to the manufacturer’s instructions. qPCR was performed using primers targeting the bacterial gapA gene (gapA_F and gapA_R; amplicon length, 301 nt), as well as primers specific for early and late phage transcripts. For phage Lambda, primer pairs targeting early and late genes were used: Lambda_early_F/R with an amplicon length of 213 nt and Lambda_late_F/R with an amplicon length of 209 nt, respectively.

To quantify bacterial and phage DNA, the samples from the protocol above before DNase I treatment were taken for qPCR. qPCR was performed using primer pairs targeting the bacterial chromosome, which were the same as those used for bacterial gene expression quantification (Lambda_F and Lambda_R; amplicon length, 196 nt). The phage-to-cell DNA ratio at each time point was calculated as 2^ΔCt, where ΔCt was defined as the difference between the chromosomal and phage Ct values. Primers are listed in Supplementary table 1.

### Time-lapse microscopy of infected cells in the presence of intercalators

The overnight culture of *E. coli dam-* was grown in LB medium with appropriate antibiotics and 0.03% arabinose at 30 °C and 220 rpm for 17 h. Next day, the culture was diluted 1:100 into 5 mL of fresh LB medium with 10 mM MgCl_2_, 5 mM CaCl_2_ and 0.03% arabinose and grown to an OD600 of ∼0.4 for about 2 hours. Then 50 µL of cells were added to daunorubicin or Hoechst 33258 or DMSO were added and incubated for 5 min at 30 °C and 220 rpm. 50 µL of cells (8,9×10^7^ CFU/mL) were mixed with 20 µL of a phage (2,3×10^9^ PFU/mL) to achieve MOI 10. The mixture was incubated at 30 °C for 10 minutes without shaking. After that the timer for photos was started. To prepare the sample for the microscopy, the phage-cell mixture was 2-fold diluted and 1 µL of the mixture was dropped on the 1 mm LB+1.2% agarose pad supplied with 10 mM MgCl_2_, 5mM CaCl_2_, 0.03% arabinose and intercalates or 1% DMSO in equivalent concentration. After the drop drying, the pad was slipped over to the confocal dish and transferred to the microscope.

The imaging was performed on Nikon Eclipse TE2000-U inverted epifluorescent microscope using a 60× objective (Plan Apo; numerical aperture [NA], 1.40; oil immersion) with camera (Dhyana 400BSI V3), within the cage of an incubator (Solent Scientific) at 30 °C and mercury lamp USH-102D (USHIO) equipped with neutral density filters (ND4, ND8) to reduce photobleaching. Cells were imaged under phase contrast and the following fluorescent filter cubes B-2E/C, UV-2A, G-2A (Nikon). During the time-lapse movie, the sample was imaged in phase-contrast (20 ms exposure for cell recognition), EGFP (in B-2E/C, 20 ms exposure), daunorubicin (G-2A, ND4 and ND8 switched on, 20 ms exposure), Hoechst 33258 (UV-2A, ND8 switched on, 100 ms exposure) in channels at intervals of 7 min for the first 30 min and then 13 min until cell fate was visible (3 h 30 min in total). Image preprocessing was performed semi-automatically in ImageJ software.

## Results

### Design of DNA-binding fluorescent Hoechst derivatives

We hypothesized that the class of aromatic heterocyclic compounds known as bisbenzimidazoles might represent a promising source of novel phage inhibitors. Bisbenzimidazole-based dyes such as Hoechst 33258, Hoechst 33342, and their derivatives are known to bind site-specifically and non-covalently to three consecutive A/T base pairs in the minor groove of B-form DNA (24). Their affinity to nucleic acids and fluorescence properties can be enhanced by creating bisbenzimidazole dimers, typically via a “head-to-head” linkage. The typical design for such dimers is a “head-to-head” linkage via a linker, whereas the reverse “tail-to-tail” configuration has never been explored (Fig. 1A, S1). In this study, we designed and synthesized several new bisbenzimidazoles, including two reverse “tail-to-tail” dimers (RDB_2_(3) and RDB_2_(7)), a “head-to-head” dimer (DB_2_Py), and a halogen-containing monomer (MB_2_Cl) to test their effect on viral infection. These novel compounds were compared with the known dimers DB_2_(3), DB_2_(7), DB_2_P(1), and Hoechst 33258 (Fig. S1A,B) (25,26). All eight compounds differ in their overall geometry and level of freedom, likely affecting their DNA-binding capabilities.

**Figure 1.**
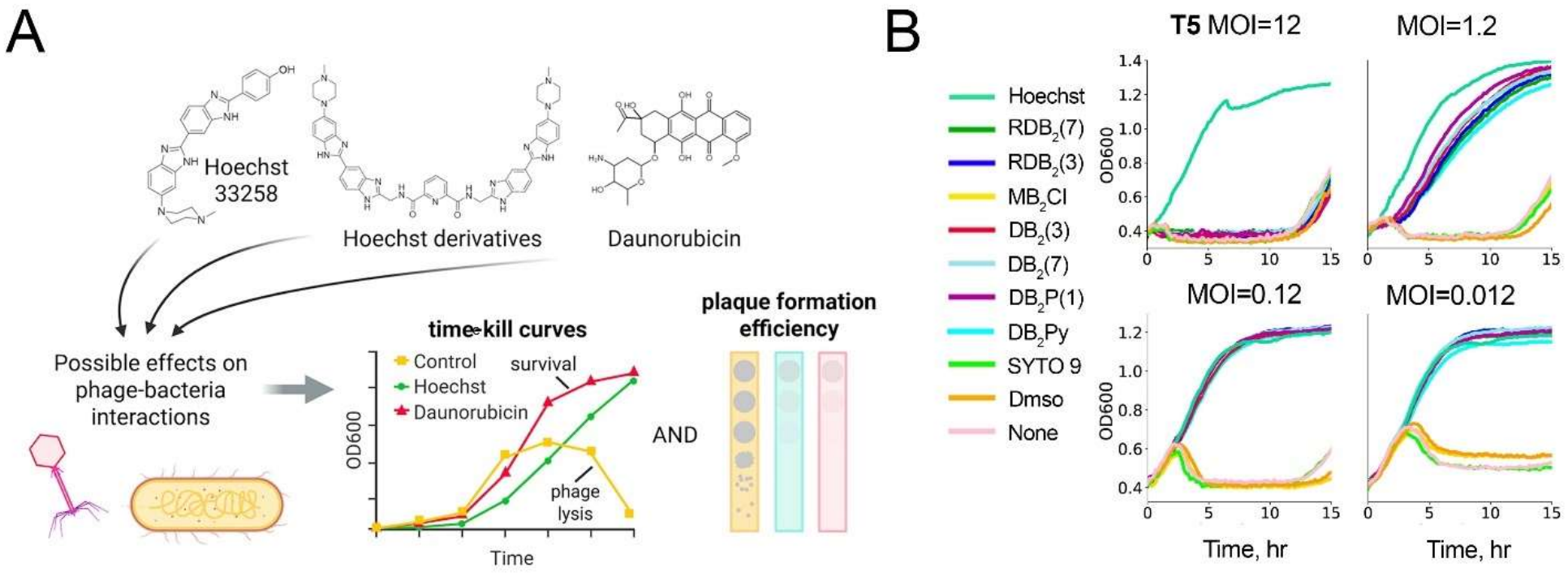
The scheme of the study and T5 infection in the presence of DNA dyes. **A)** Schematic overview of phage susceptibility assays in liquid and solid culture used to assess the effects of DNA-binding compounds on the phage infection cycle. **B)** Growth curves of T5-infected bacteria at four MOIs in the presence of Hoechst 33258, Hoechst-derived compounds, SYTO 9, or DMSO.

To confirm DNA binding by these compounds, we measured changes in fluorescence upon addition of dsDNA (Fig. S1C). The “tail-to-tail” dimers RDB_2_(3) and RDB_2_(7) showed no fluorescence in the absence of DNA, but fluorescence increased upon addition of dsDNA, confirming the DNA-binding activity of these molecules. Classical “head-to-head” bisbenzimidazole DB_2_Py demonstrated a hypsochromic shift upon increasing nucleic acid concentration. MB_2_Cl was the only compound that exhibited decreased fluorescence upon DNA interaction, potentially due to aggregation of the formed complexes. Notably, the amplitude of fluorescence change was the highest in the case of RDB_2_(3) suggesting that it formed tight complexes with DNA (Fig. S1C).

### Natural and synthetic DNA binding compounds inhibit phage infection in *E. coli*

We tested nine bisbenzimidazoles (Hoechst 33342, Hoechst 33258 and 7 new variants) alongside the common nucleic acid stain SYTO 9 and daunorubicin for their ability to inhibit phage infection in *E. coli* (Fig. 1B). First, we identified boundary concentrations for all compounds, at which cell growth was not inhibited (Fig. S2). Next, we measured growth curves for cell cultures infected with classical phages T5, P1 and λ in the presence of each of the compounds. T5-mediated lysis was suppressed without affecting cell growth in the presence of 25 μM Hoechst 33258 and 115 μM daunorubicin (Fig. 1B). In comparison, Hoechst 33342, which is known to penetrate cell membranes efficiently, affected viral infection at lower concentrations (3-6 μM), but was toxic at these concentrations and was therefore excluded from further screening (Fig. S2, S3). Most compounds (Hoechst 33258, RDB_2_(7), RDB_2_(3), MB_2_Cl, DB_2_(3), DB_2_(7), DB_2_P(1)) prevented T5-mediated lysis at Multiplicities of Infection (MOI) ranging from 0.012 to 1.2 (Fig. 1B, Fig. S3-S4). Remarkably, Hoechst 33258 remained effective even at an MOI of 12, where each bacterial cell should be infected, arguing against abortive infection mechanism. In contrast, DB_2_Py and SYTO 9 did not prevent T5-mediated lysis. Hoechst 33258 also inhibited P1 and λ infection, whereas none of the other compounds affected these phages (Fig. S4). Given their broad activity, we selected Hoechst 33258 as a new synthetic antiphage molecule and daunorubicin as a well-known natural antiphage agent for further investigation.

### Hoechst 33258 and daunorubicin show distinct phage specificity profiles

To compare the spectrum of activity of Hoechst 33258 and daunorubicin, we tested them against a panel of phylogenetically diverse phages by measuring the efficiency of plating (EOP) on solid media (Fig. 2A). Hoechst 33258 completely abolished plaque formation by Bas27, Bas33, Bas34 (*Demerecviridae*); HK446 (*Hendrixvirinae*); MSK8 (*Vequintavirinae*); P2 (*Peduovirus*); 9g (*Queuovirinae*), mEp213 (*Andersonviridae*), and almost completely for phage T5 (*Markadamsvirinae*). Hoechst 33258 also reduced EOP by one to several orders of magnitude for viruses HK106, HK542, HK75 (*Hendrixvirinae*); HK225, HK578 (*Dhillonvirus*); P1 (*Punavirus*). In contrast, Hoechst 33258 had no effect on plaque formation by phages T6 (*Straboviridae*) and T7 (*Autotranscriptaviridae*) (Fig. 2A, Fig.S5). We also observed altered phage spot morphology in the presence of Hoechst 33258, characterized by smaller spot size and less defined borders, for MSK1, MSK2, MSK6, MSK7 (*Drexlerviridae*), HK106, HK542, HK578, HK75, and T5.

**Figure 2.**
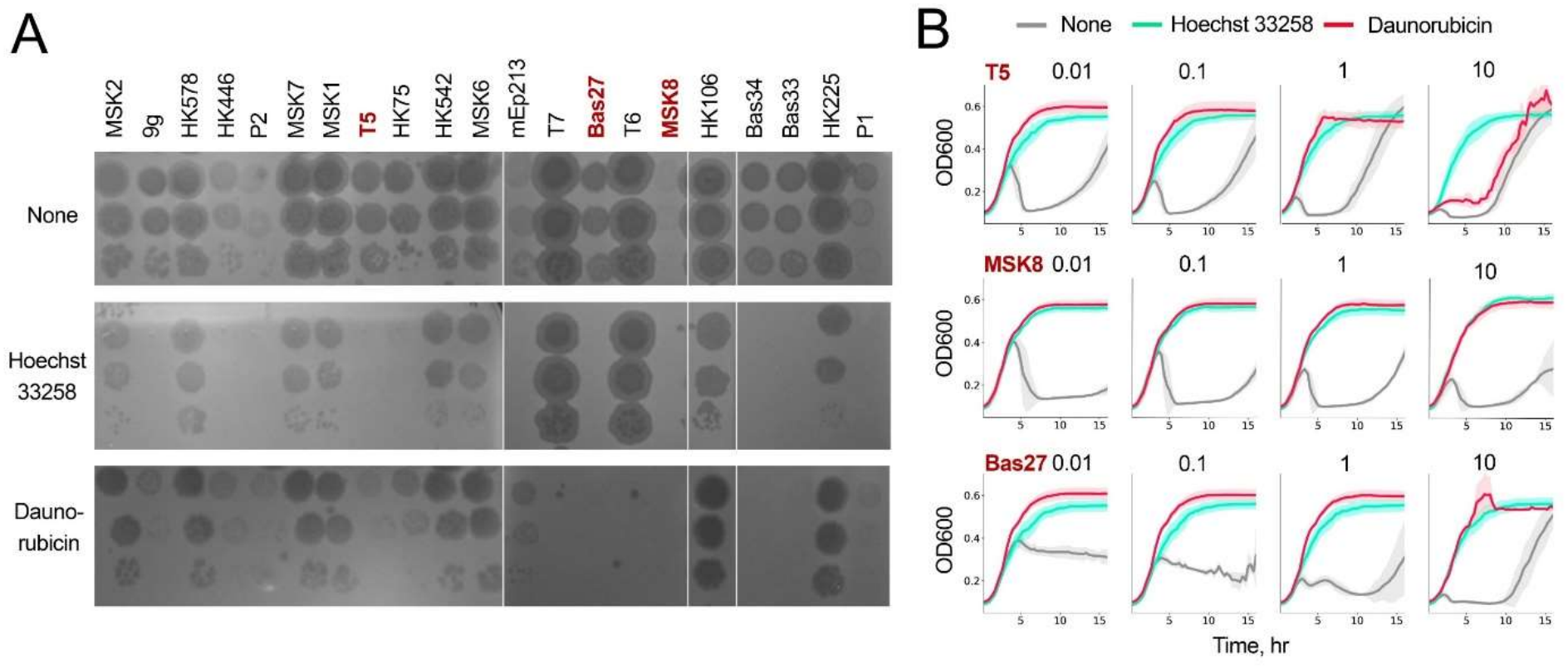
Antiphage activity of Hoechst 33258 and daunorubicin. **A)** Plaque assays for a panel of *E. coli* phages in the presence of Hoechst 33258 or daunorubicin. **B)** Growth curves of bacteria infected with T5, MSK8, or Bas27at four MOIs in the presence of Hoechst 33258 or daunorubicin (3 replicas).

Daunorubicin showed partially overlapping but distinct inhibition patterns. It strongly reduced plaque formation by Bas27, Bas33, Bas34, MSK8, T6, T7; had moderate effects on HK446, HK75, mEp213, 9g, P2, P1, and T5, and showed no effect on HK542, MSK2, phages. Changes in phage spot morphology in the presence of daunorubicin were also observed for several phages (HK106, HK225, HK578, HK75, P2, T5, MSK1, MSK7) (Fig. 2A).

We further compared the effects of Hoechst 33258 vs daunorubicin on viral infection at different MOIs in liquid cultures. In contrast to Hoechst 33258, daunorubicin did not protect *E. coli* from T5 infection at MOI=10 (Fig. 2B). Both daunorubicin and Hoechst 33258 provided complete protection against phages Bas27 and MSK8 at MOI=10 (Fig. 2B). Conversely, only daunorubicin provided protection against phage Qβ (the only RNA phage included in our analysis), even at high MOI (Fig. S5). Furthermore, Hoechst 33258 protected cells from HK75 and HK446 across the entire tested MOI range. In contrast, daunorubicin was effective only at low MOI, a finding consistent with the EOP data (Fig. S6). P2 and mEp213 infections were inhibited by both compounds (Fig. S6), while only Hoechst 33258 was efficient against mEp213 phage in the EOP assay (Fig. S5). T6 infection was prevented by daunorubicin in solid media, but not in liquid culture at all tested MOIs (Fig 2A, Fig. S5B).

These results demonstrate that both DNA binders can block phage infection, with good correlation between the bulk liquid and solid media assays, but the patterns of protection are different between these compounds and are phage-specific. The observed prevention of phage infection at high phage titers argues against the abortive infection mechanism previously proposed for daunorubicin (1). We therefore investigated the exact mechanism of antiphage activity of these compounds in further experiments.

### Tracking cell growth and phage infection by fluorescence measurements

To track the dynamics of DNA-binding agents during phage infection, we monitored the fluorescence of three compounds – daunorubicin (a DNA intercalator that affects phage infection), Hoechst 33258 (a minor groove DNA binder that affects phage infection), and SYTO 9 (a minor groove binder that does not affect phage infection, used as a control) during infection with T6, T7, Bas27, Bas34 (Fig. S6). Daunorubicin fluorescence signal (measured at 600 nm) slowly decreased during incubation in conditions of inhibited infection (low MOI for each phage, Fig. S6B,E) but remained almost constant when cell growth was constrained by phage infection (high MOI, Fig. S6C,F); the observed changes anti-correlated with the cell density and could possibly be explained by light absorption. In contrast, Hoechst 33258 showed a peak in fluorescence signal (measured at 455 nm) during successful infection with phages T6 and T7 at both MOI, coinciding with cell lysis detected by OD600 measurements. No increase in Hoechst 33258 fluorescence was observed for protected cells with phages Bas27 and Bas34 (except for late stages of cell growth at high MOI) (Fig. S6B,C,E,F). Similarly, an increase in fluorescence signal from SYTO 9 (measured at 510 nm) coincided with cell lysis during infection with T6, T7, Bas27, and Bas34 at both MOI. The observed changes could be likely explained by Hoechst 33258 and SYTO 9 binding to extracellular DNA after cell lysis and can thus be used as an additional marker of culture lysis and infection dynamics (27). In contrast, daunorubicin fluorescence is not increased after cell lysis, making it difficult to track culture growth by following its fluorescence.

### Hoechst 33258 inhibits phage replication and transcription

Our initial experiments identified Hoechst 33258 as the most potent inhibitor of phage infection among the newly tested compounds. To further understand the effects of Hoechst 33258 on phage development, we measured viral RNA and DNA levels during phage λ infection, which was chosen because Hoechst 33258 had moderate protection effect with this phage and prevented cell lysis at low but not at high MOI (Fig. 3A). In the presence of Hoechst 33258, phage DNA amount normalized to bacterial genomic DNA was decreased by one-two orders of magnitude at all measured time points (15 to 90 minutes after phage addition at MOI of 0.3), correlating with significantly delayed lysis (Fig. 3B). Importantly, in the presence of Hoechst 33258 the ratio of phage to cell DNA remained <1, indicating inefficient infection. This reduction in the bulk phage replication rate could be explained by a decrease in the number of cells supporting phage infection, or decreased phage progeny rate across all infected cells. To distinguish between these possibilities, we measured the levels of early and late phage transcripts in the absence and in the presence of Hoechst 33258.

**Figure 3.**
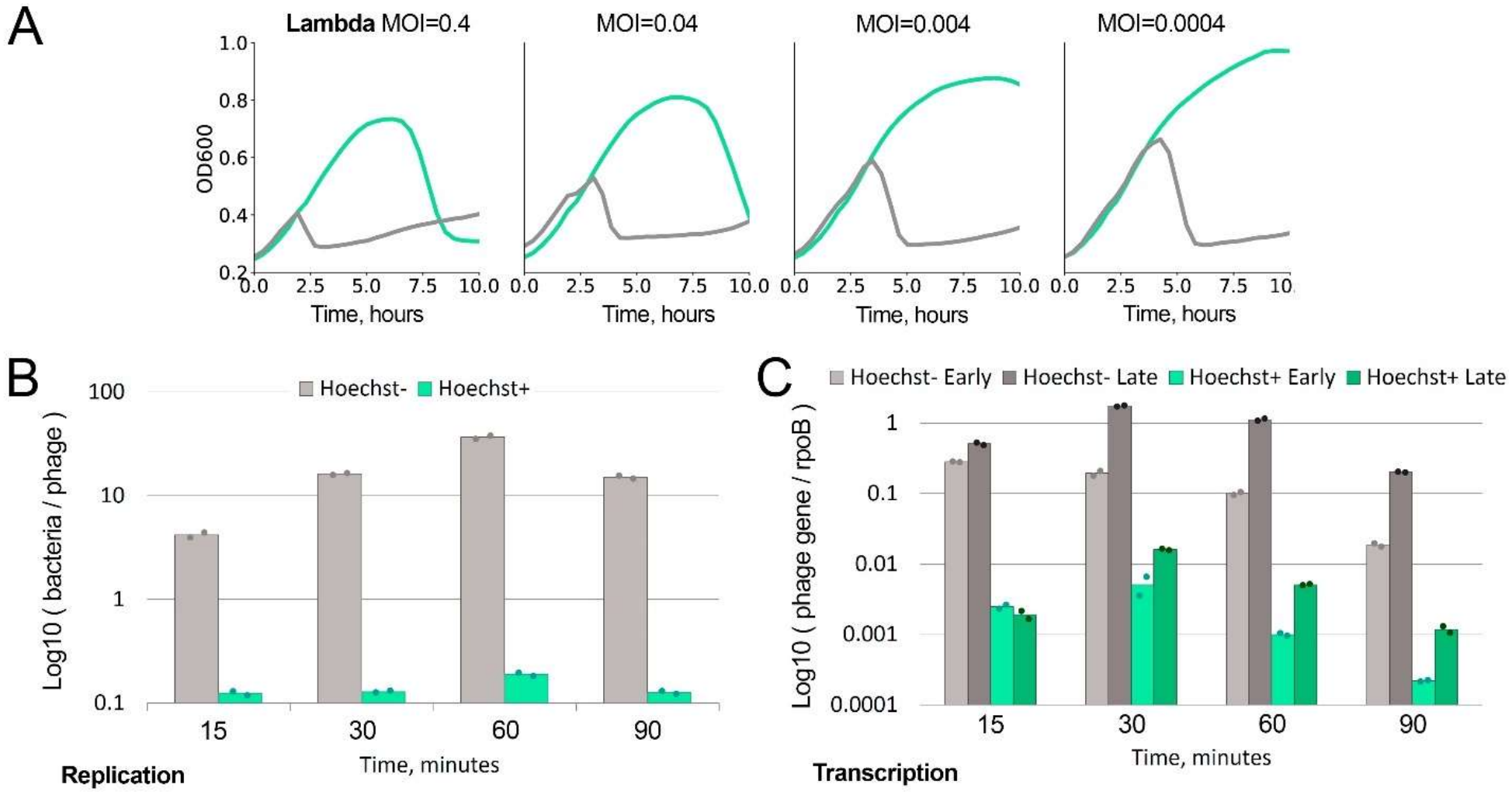
Phage DNA processing in infected bacteria during Hoechst 33258 treatment. **A)** Changes in phage λ DNA levels during infection in the presence of Hoechst 33258, normalized to bacterial DNA. **B)** Changes in transcription of early and late genes of phages λ during infection, normalized to the bacterial rpoB gene.

If a small number of cells produced the same number of viral particles per cell in the presence of Hoechst 33258, while the majority of population remained uninfected, we would not observe a shift in the early to late gene ratio. In the absence of Hoechst 33258, early gene expression was the highest at 15 minutes and gradually decreased by 90 minutes, while late gene expression peaked at 30-60 minutes after infection start, reflecting the end of the first round of infection. Hoechst 33258 addition decreased both early and late genes expression by two orders of magnitude. Importantly, the late to early gene ratio was lower at all time points in the presence of Hoechst 33258 (1.8/0.8, 9/3.2, 11.2/5.1, 10.9/5.4 minus/plus Hoechst 33258 at 15, 30, 60, 90 minutes) (Fig. 3C). This indicates that Hoechst 33258 impairs the transition from early to late stages of viral cycle, not simply reducing the number of infected cells. This is consistent with a model in which Hoechst 33258 acts at early steps of infection (as previously proposed for daunorubicin) (1,20).

### Hoechst 33258 acts during early steps of infection without directly inactivating virions

To identify the step(s) of the viral cycle affected by Hoechst 33258, we pre-mixed bacteria with phage T5, and added the compound at different time points post-adsorption. When the compound was present in the initial phage-bacteria mixture, it fully prevented lysis at MOI of 0.1 or 1 (Fig. 4A). When it was present only during early infection steps (the first 30 minutes of infection, followed by washing), it also prevented cell lysis (with normal cell growth observed during first 4 hours). In contrast, Hoechst 33258 did not protect bacterial population form lysis, when added 30 minutes after phage-bacteria mixing at MOI 1. However, at low MOI, the population survived, likely due to the low number of initially infected cells, with Hoechst 33258 preventing subsequent rounds of infection. These results indicate that the compound must be present during the adsorption or/and DNA injection steps to exert its full protective effect.

**Figure 4.**
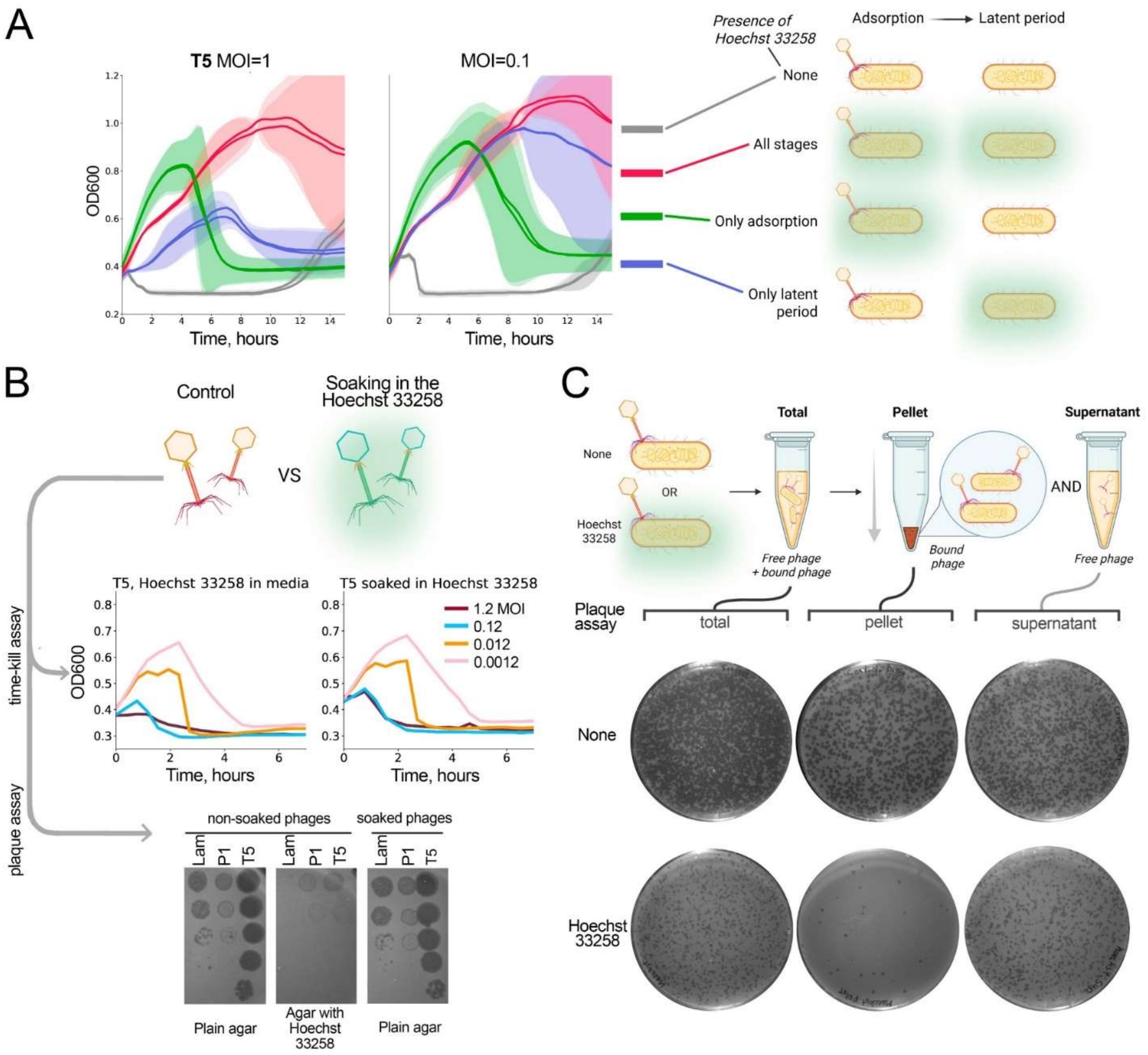
Inference of the phage infection stage affected by Hoechst 33258. **A)** Growth assay of T5-infected bacteria at two MOIs with Hoechst 33258 added at different stages of infection: (1) before infection and maintained throughout the assay, affecting all stages; (2) before infection and washed out after phage adsorption, affecting adsorption only; and (3) after phage adsorption, affecting the latent period only. **B)** Effect of pre-soaking T5 phages in Hoechst 33258 on phage infectivity in liquid culture and plaque formation efficiency. Pre-soaked phages were compared with untreated phages in a time-kill assay and plaque formation assay. **C)** Assay measuring phage adsorption efficiency and loss of plaque formation in the presence of Hoechst 33258. After T5 adsorption in the presence of Hoechst 33258, samples were collected either directly from the culture for plaque formation analysis (“total” sample) or after centrifugation from the pellet and supernatant fractions.

We next tested whether Hoechst 33258 can directly inactivate phage particles. Phage particles (T5, P1 or λ) were pre-soaked in 25 μM solution of Hoechst 33258 and then mixed with bacterial cultures, with 10-fold dilution reducing the compound concentration below the effective threshold. Under these conditions, no reduction in infectivity was observed compared to controls both in liquid and solid media (Fig. 4B, Fig. S7). Thus, Hoechst 33258 does not irreversibly damage virions but requires continuous presence during early steps of infection.

Infection in solid media further confirmed that Hoechst 33258 acts at early stages of the phage life cycle, likely during adsorption and/or injection. First, we performed a simple double-layer agar (DLA) assay with phage T5, which showed that Hoechst 33258 inhibited plaque formation (Fig. S8A), consistent with the EOP and growth curves data. This effect could be due to prevention of adsorption and subsequent infection, or due to severely impaired viral reproduction leading to undetectable plaques.

Next, to directly test whether Hoechst 33258 affected the virus adsorption step, we measured the titer of the free phage particles after infected culture centrifugation (Fig. 4C, S8C). After brief co-incubation of phage T5 (∼10^3^ particles) with bacteria in the presence or absence of Hoechst 33258, we separated cells and adsorbed phages (pellet) from free phages (supernatant) and quantified PFU values in each fraction in comparison with the unfractionated sample. In the absence of Hoechst 33258, the infectious phage particles were equally distributed between the pellet and supernatant (Fig. 4C, Fig. S8C top raw of plates). In the presence of Hoechst 33258, the total number of plaques was strongly reduced and they were primarily observed in the supernatant fraction, while the pellet fraction produced no centers of infection (Fig. 4C, second replicate in Fig. S8C). This result suggests that adsorption itself was not completely inhibited, yet phages adsorbed to bacterial cells in the presence of Hoechst 33258 did not produce viable progeny.

Finally, to check whether Hoechst 33258 could completely prevent phage infection after adsorption, we performed an ECOI (Efficiency of Center Of Infection) assay. This assay allows to calculate the number of successfully infected bacteria by plating them on a lawn formed by untreated cells; this allows visualization of phage plaques even if the burst size if strongly reduced in initially infected bacteria (Fig. S8B) (27). In this experiment, *E. coli* cells were pre-incubated with phage T5 in the absence or in the presence of Hoechst 33258 to allow phage adsorption, washed out of unbound phages and dye, and mixed with an excess of untreated *E. coli* cells for further plating on DLA (Fig. S8B). The ECOI results showed strong inhibition of plaque formation in the presence of Hoechst 33258, indicating that it fully inhibits phage production in the affected cells.

In summary, these experiments suggest that Hoechst 33258 irreversibly inactivates the virus during initial cell contact or short after it but is much less efficient if added during the latent period, after the injection of viral DNA.

### Analysis of infection in single cells in the presence of Hoechst 33258 and daunorubicin

While the experiments described above reflect population-level measurements, single-cell analysis may help to clarify the underlying mechanisms and cell-to-cell variability during infection. We monitored infection of phage Bas27, strongly affected by both Hoechst 33258 and daunorubicin, at MOI=10 using time-lapse microscopy of living cells on agarose pads. Control *E. coli dam*^-^ cells grown without virus demonstrated elongated morphology (Fig. 5, top). We explain it by SeqA-EGFP expression which was aimed to monitor viral DNA entry, but Bas27 was lack of DNA methylation discarding EGFP foci signal from analysis (28). During infection with Bas27 at high MOI (10), we observed growth arrest and cell lysis during the timeframe of the experiment (Fig. 5, Supplementary Video). Most *E. coli* cells treated with daunorubicin showed normal growth during infection, yet some demonstrated growth arrest, which coincided with accumulation of intracellular daunorubicin signal during the first 15 min (fourth row, red cells). At the same time, Hoechst 33258 was accumulated in the half of cells (third row, blue cells) and partially delayed cell division (Fig. 5). We therefore conclude that similar to recently proposed model (20), phage injection and transmembrane channel opening enable entry of fluorescent dyes into a subpopulation of cells, while some cells potentially remain protected without DNA binders’ entry if MOI 10 was enough to infect every single cell.

**Figure 5.**
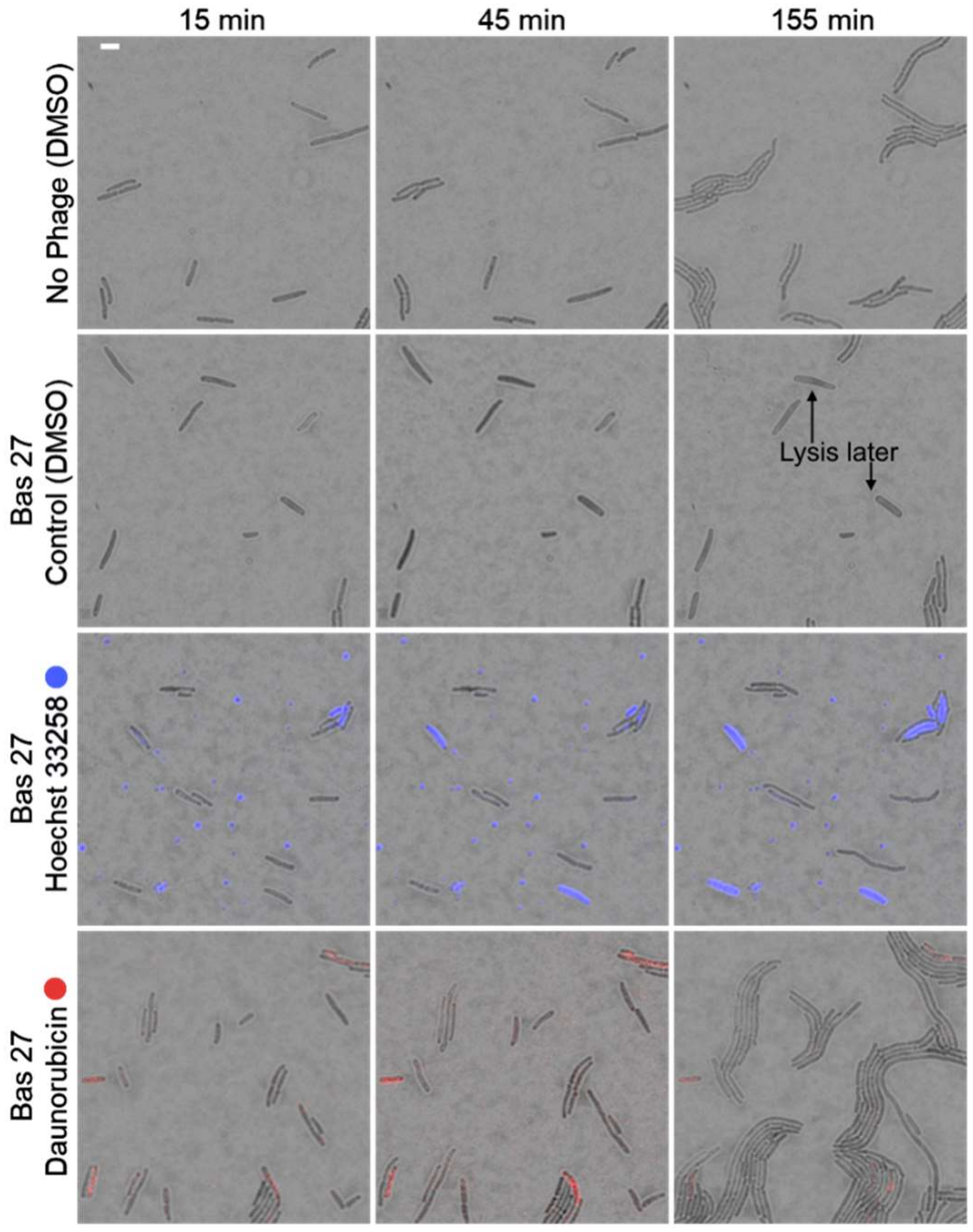
Time-lapse fluorescent microscopy of cells during Bas27 phage infection in the presence of daunorubicin and Hoechst 33258. Infection (MOI of 10) was started in liquid culture (OD of 0.4), then cells were applied on the LB agarose pads with addition of daunorubicin (222 μM, red), Hoechst 33258 (25 μM, blue) or DMSO (1%) and imaged during 3-hours intervals in the 30 °C chamber. Samples without phage infection were also monitored (the bottom panel). The sample was imaged using phase-contrast for cell recognition and fluorescent filters: G-2A for daunorubicin (red emission), UV-2A for Hoechst 33258 (blue emission). Scale bar is 4 μm.

## Discussion

The study of small molecules affecting viral infection in eukaryotes and prokaryotes is important both for understanding molecular mechanisms of virus-host interactions and for their application in biotechnology and therapy. For over 80 years, small DNA binders have been known to block bacteriophage infection. Natural compounds tested in the mid-20th century were recently rediscovered as components of bacterial immune systems (2). Synthetic chemicals, including both intercalating and non-intercalating DNA-binding compounds, have also been identified as phage inhibitors (1,2,19,29). Yet the mechanism of their action remains unclear.

In this work, we demonstrated the ability of the widely used DNA dye Hoechst 33258 to shut down viral infection in *E. coli* and compared the mode of its action with novel synthetic compounds and daunorubicin, previously shown to inhibit phage infection. Among the tested compounds, Hoechst 33258 was the most active while its derivatives demonstrated narrower ranges of antiviral activity. Future modifications would be necessary to obtain a molecule with a broader selectivity.

To uncover the mechanism of antiviral activity of DNA-binding dyes, we performed analysis of Hoechst 33258 in comparison with daunorubicin in solid and liquid media assays using a panel of phages. Our population-level microbiological analysis revealed that free viruses were not inactivated by the studied compounds, but that phage infection was blocked at an early stage of infection, likely before or during the DNA injection step. Notably, we found that Hoechst 33258 or daunorubicin protected the cells from several phages even at high virus titers (MOI 10), with all the cells infected in the absence of these compounds in these conditions. These observations argue against an abortive infection mechanism caused by the dye itself and suppose that it’s activity prevents accumulation of the toxic phage products (1). A single-cell microscopy study recently suggested that daunorubicin entered the cells during λ phage injection at early steps, thus leading to cell death and blocking subsequent steps of infection (20). Importantly, however, we detected dividing viable cells that lacked phage DNA and intracellular dye in the cultures infected at high MOI, indicating that both dyes can provide protection without entering the cells by themselves.

We speculate that daunorubicin and Hoechst 33258 may act by physically impeding the DNA translocation step, arresting infection between membrane engagement and successful genome entry, or impede viral replication directly. It remains unknown whether the interactions of daunorubicin or Hoechst 33258 with viral DNA are required for their action. Several small molecules that do not interact with DNA (RU.521, kanamycin) were recently shown to inhibit viral infection (19). Potentially, DNA binders may also synergize with other types of immunity, such as restriction-modification systems, as recently shown for daunorubicin (1).

Intercalating and minor groove binding dyes are widely used to distinguish alive from dead cells during phage infection. Our results show that some of these dyes can by themselves act as inhibitors of infection, and their use can thus affect the experimental output. Identifying dyes that stain cells efficiently without affecting phage infection would be important in studying phage–host interactions. On the opposite, efficient DNA binders or other small molecules with a broad antiphage specificity would also be useful for studying viral infections and treating bacterial cultures in biotechnology.

In conclusion, our work demonstrated that both synthetic and natural DNA staining agents that do not prevent bacterial growth can be highly efficient as antiviral compounds acting at early steps of viral injection. The study of small molecule compounds that affect viral infection is important for understanding the simplest population-level immune mechanisms based on secreted molecules preserved over billions of years and widespread in some clinically and biotechnologically important phyla, such as *Streptomyces*.

## Supporting information

and its Supplementary Information

## Funding

This study was supported by the Ministry of Science and Higher Education of the Russian Federation (topic no. FFEW-2024-0009, viral experiments in liquid and solid media, single-cell analysis) and the Russian Science Foundation (grant 24-44-00107 to AK, measurements of phage infection dynamics using RT-PCR).

## Acknowledgements

Microscopy was performed using the equipment of IGB RAS facilities supported by the Ministry of Science and Higher Education of the Russian Federation.

## Author Contribution

Vladimir Panteleev – experiments in liquid and solid media, visualization, writing – review and editing, conceptualization, experiments design; Nika F. Petrova – experiments in liquid and solid media; Ivan Moldon – single-cell experiments and microscopy, writing – review and editing, microscopy visualization; Alena I. Sidorova – synthesis of Hoechst 33258 derivatives, writing – review and editing; Artem Isaev – providing phage collections, writing – review and editing; Aleksandr Andriianov – isolation of MSK phages; Albert Arutyunyan – synthesis of original Hoechst 33258 and it’s derivatives; Andrei Moiseenko – microscopy; Pavel N. Solyev – synthesis of original Hoechst 33258, chemicals analysis and description; Andrey Kulbachinskiy – project administration, funding acquisition, writing – review and editing, Daria Gelfenbein – conceptualization, project administration, supervision, funding acquisition, writing original draft.

## Conflict of interest

The authors declare no conflict of interests.

## Data availability

All data supporting the findings of this study are available within the paper and its Supplementary Information.

## Notes

### Competing Interest Statement

The authors have declared no competing interest.

