## Supplementary material for "Inhibition of Bacteriophage Infection by Natural and Synthetic DNA-binding Compounds": and its Supplementary Information

**Supplementary Table 1.** Primers used for qPCR assay experiments

| Primer name | 5’-3’ sequence | Description |
| --- | --- | --- |
| gapA_F | GGAACGCCATACCAGTCAGT | Primer for qPCR amplification of rpoB. |
| gapA_R | AAAGGCGCTAACTTCGACAA | Primer for cDNA synthesis and qPCR amplification of rpoB. |
| Lambda_F [Lam_q_4_F] | AAACGTCCTTCTCGGTGCAT | Primer for qPCR amplification of Lambda genome targeting the proximity of ea31 gene |
| Lambda_R [Lam_q_4_R] | CAGTGCCTAGCAAACTCGGA | Primer for qPCR amplification of Lambda genome targeting the proximity of ea31 gene |
| Lambda_early_F [Lam_q_1_F] | ATCGCACCATCAGCCAGAAA | Primer for qPCR amplification of Lambda early gene transcript (replication protein O) |
| Lambda_early_R [Lam_q_1_R] | GCCTGCCTGTTGCTTGTTAC | Primer for qPCR amplification of Lambda early gene transcript (replication protein O) |
| Lambda_late_F [Lam_q_5_F] | ACGGCTACTCCGTGTTTGAG | Primer for qPCR amplification of Lambda late gene transcript (portal protein B) |
| Lambda_late_R [Lam_q_5_R] | CTCTTCCAGCCAGCACAGAA | Primer for qPCR amplification of Lambda late gene transcript (portal protein B) |

Section 1. Supplementary figures

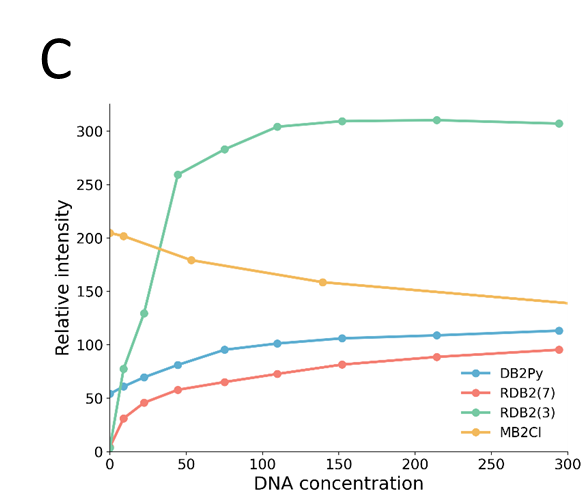

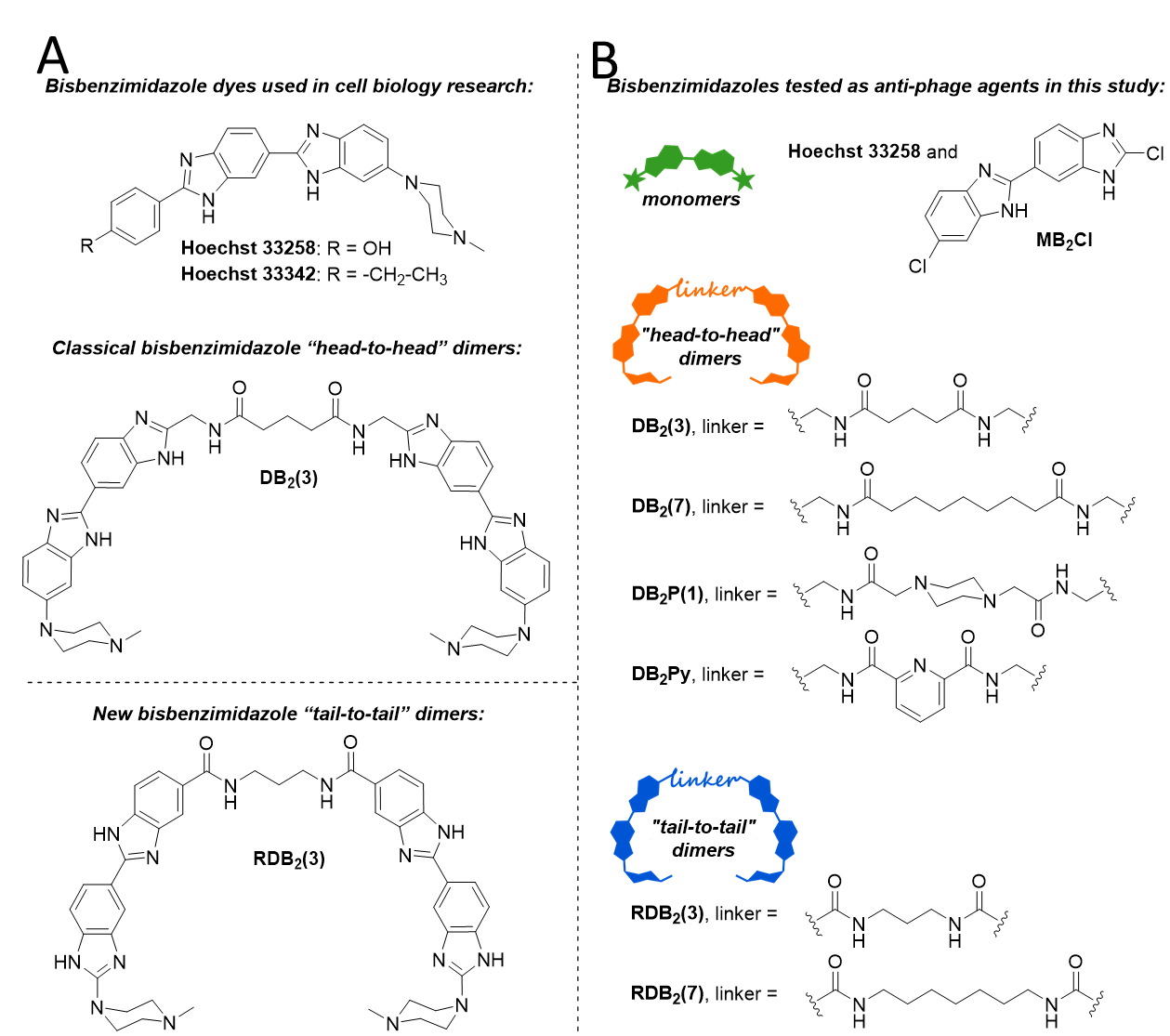

**Figure S1.** Chemical structures of Hoechst-like compounds. (A) Structure of Hoechst dyes used in cell biology research. (B) Hoechst-based compounds used in the study. (C) Comparison of fluorescence intensity of new bisbenzimidazoles when binding to dsDNA.

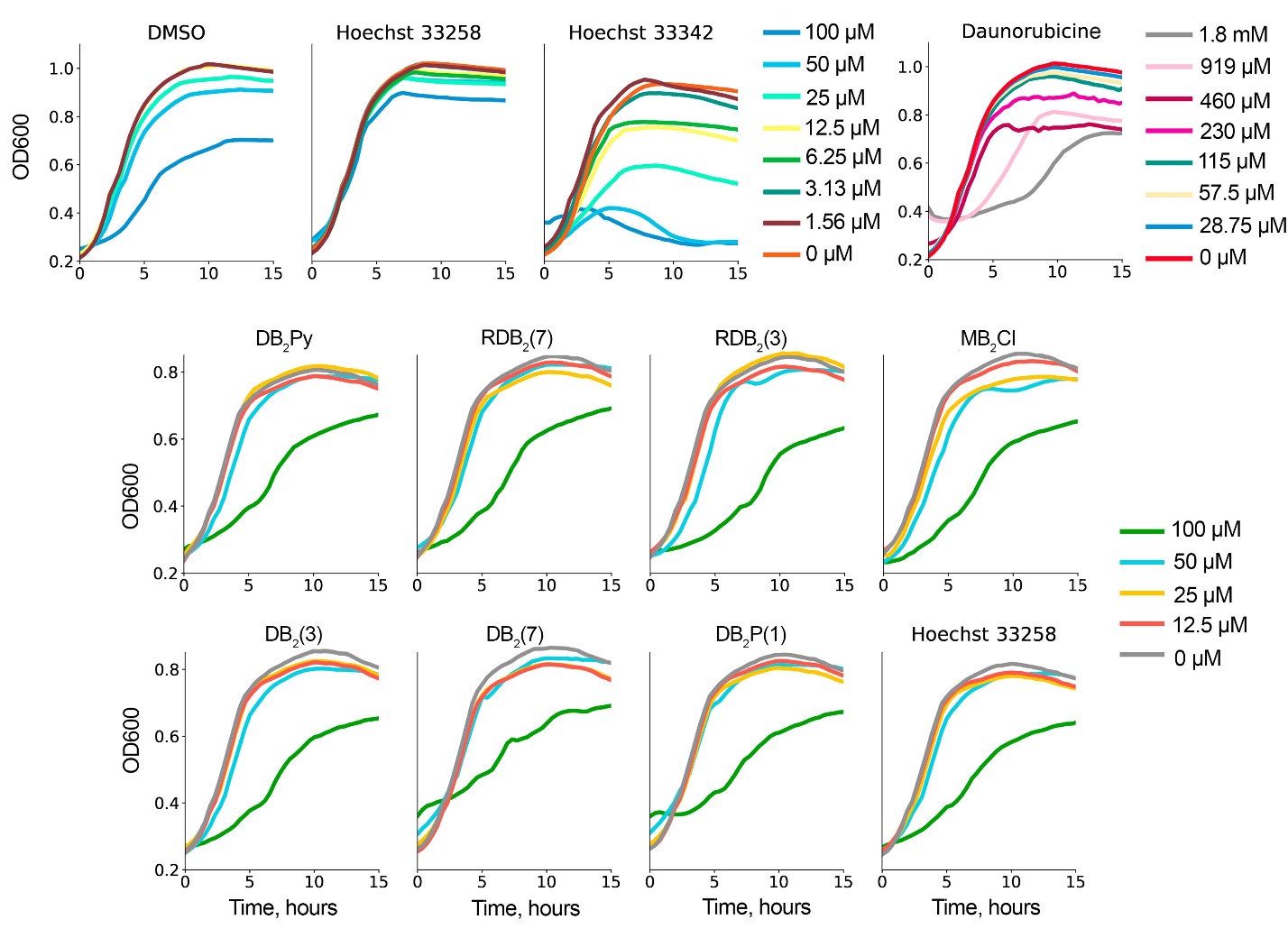

**Figure S2.** Cell growth in the presence of different concentrations of Hoecht-like compounds and daunorubicin. The highest concentrations that did not impair cell growth were 25 uM for Hoechst 33258, RDB_2_(7), RDB_2_(3), MB_2_Cl, DB_2_(3), DB_2_(7), DB_2_P(1), DB_2_Py, 115 uM for daunorubicin, and 3.13 uM for Hoechst 33342. These concentrations are referred to as working concentrations and were used in experiments if not noted otherwise.

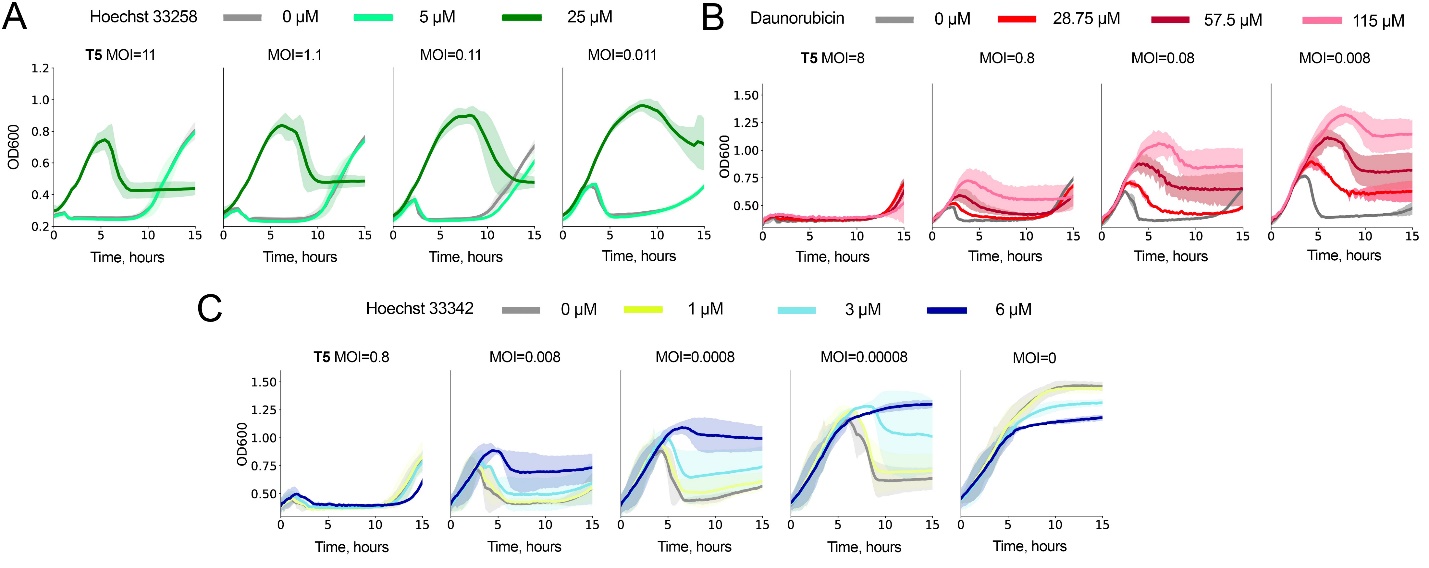

**Figure S3.** Growth curves of T5-infected bacteria at three MOIs in the presence of three concentrations of Hoechst 33258 (A), daunorubicin (B), and Hoechst 33342 (C).

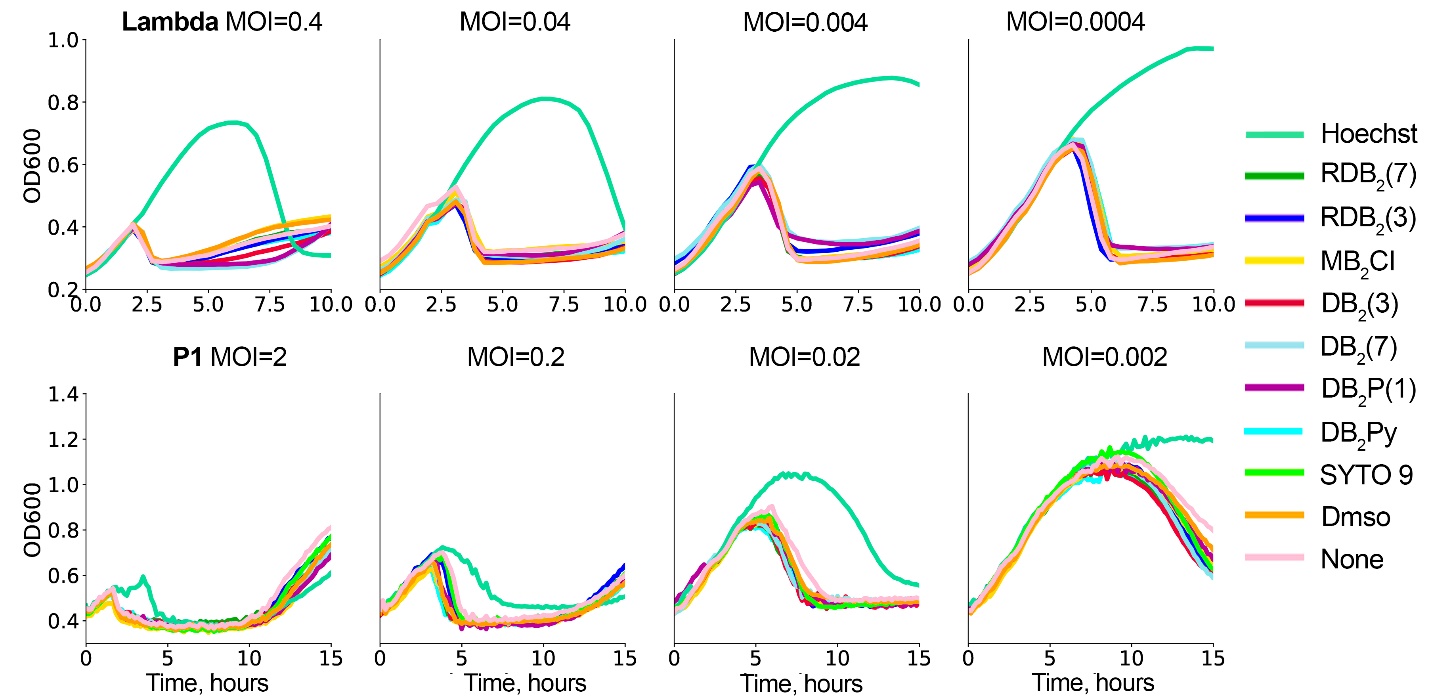

**Figure S4.** Growth curves of P1 and Lambda-infected bacteria at four MOIs in the presence of Hoechst-like compounds, SYTO 9, or DMSO.

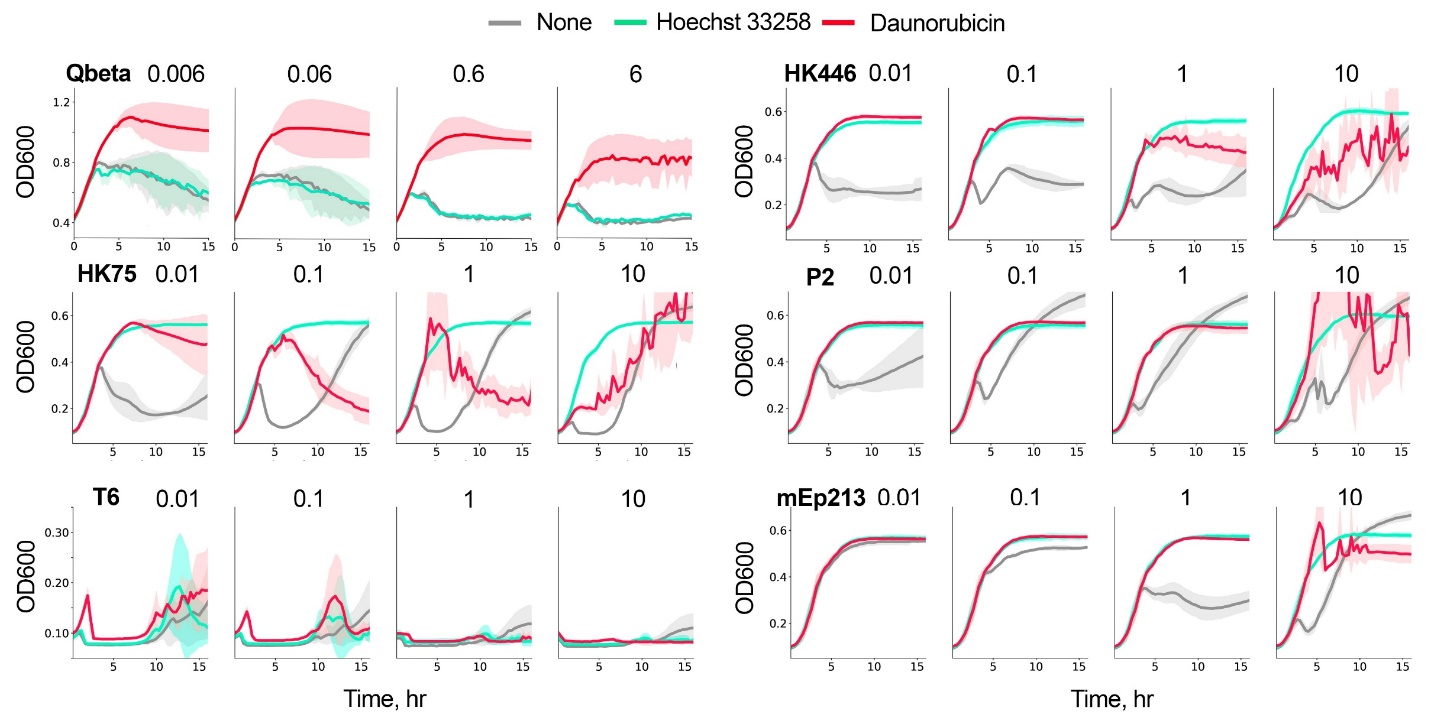

**Figure S5.** Growth curves of Qbeta, HK75, T6, HK446, P2, and mEp213-infected bacteria at four MOIs in the presence of Hoechst 33258 and daunorubicin (result of 3 experiments).

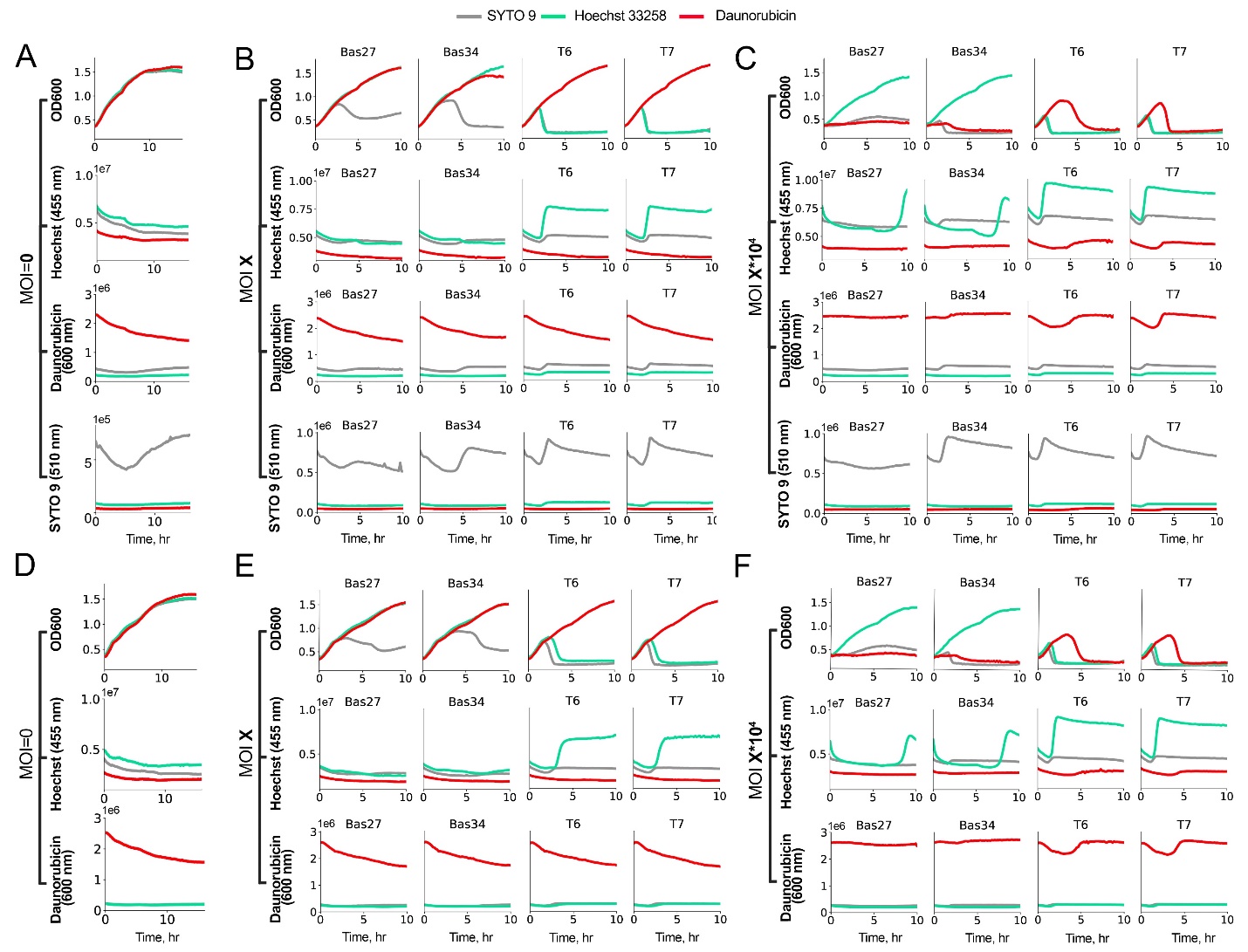

**Figure S6.** Detection of DNA-binder penetration into cells by fluorescence measurements during Bas27, Bas34, T6, and T7 phage infections. For each bacterial culture, OD600 and fluorescence signals from Hoechst 33258, daunorubicin, and SYTO 9 were monitored. Panels **A-C** and **D-F** correspond to the first and second replicates of the experiment, respectively. Panels **A** and **D** show non-infected control cultures; panels **B** and **E** show infections at MOI=0.0013, 0.002, 2e-4, and 1.2e-3 for phages Bas27, Bas34, T6, and T7, respectively; and panels **C** and **F** show infections at 10,000-fold higher MOI for each phage.

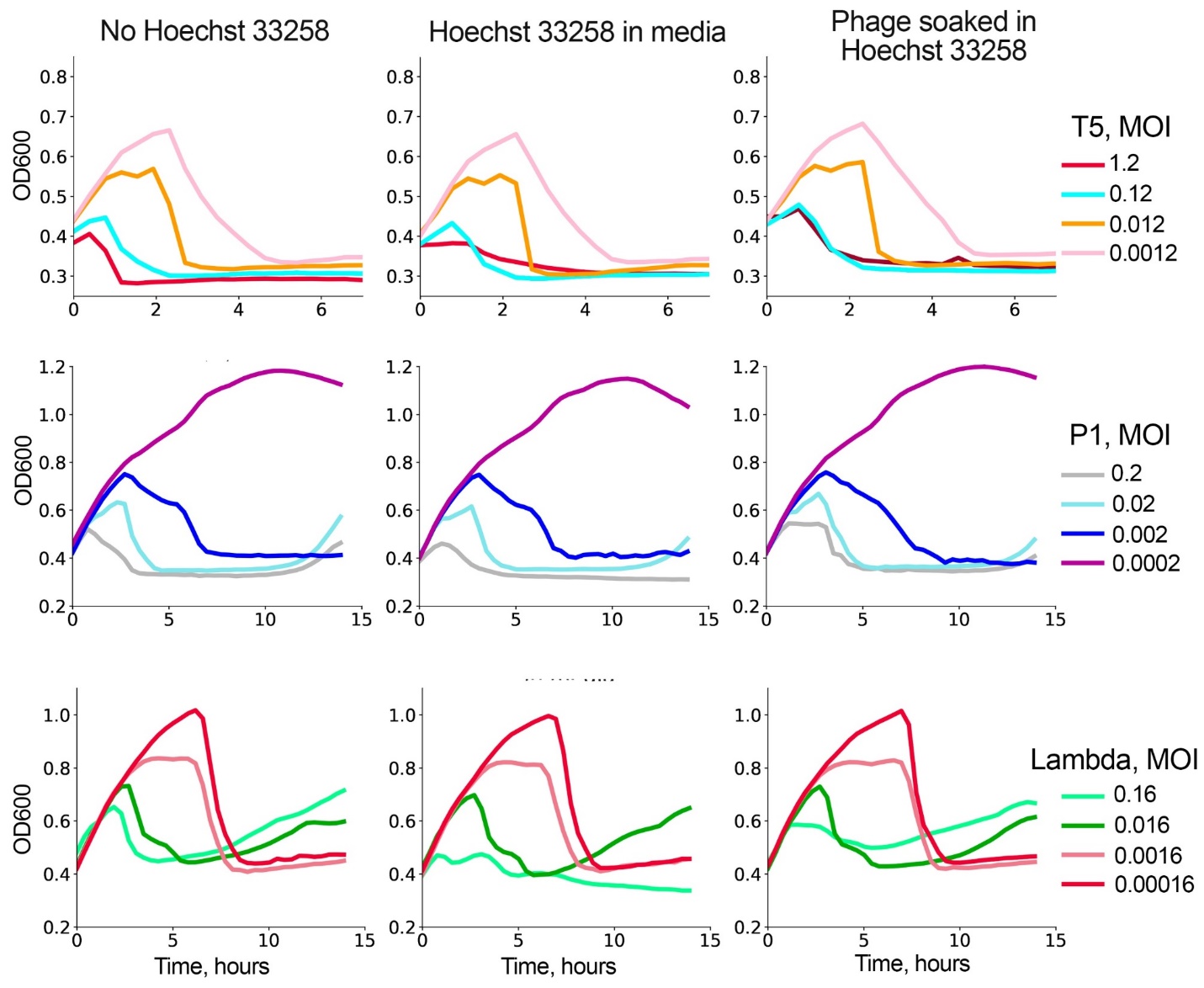

**Figure S7.** Effect of pre-soaking T5, P1 and Lambda phages in Hoechst 33258 on phage infectivity in liquid culture across several MOIs

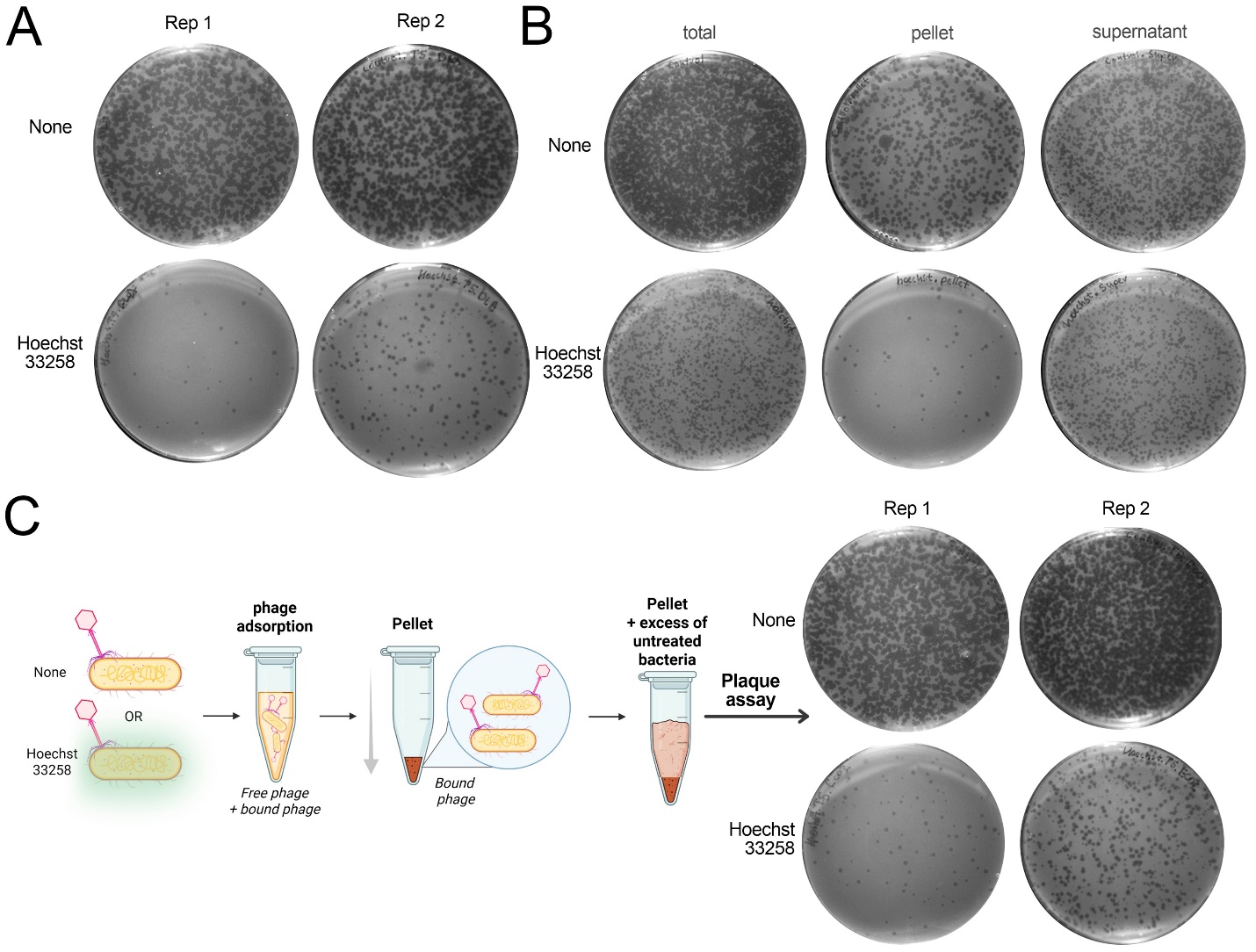

**Figure S8.** Plaque formation in the presence of DNA-binding compounds. **A)** Second replicate of the assay measuring Т5 phage adsorption efficiency and loss of plaque formation in the presence of Hoechst 33258. After adsorption in the presence of Hoechst 33258, samples were collected either directly from the culture for plaque formation analysis (“total” sample) or after centrifugation from the pellet and supernatant fractions. **В)** Efficiency of the center-of-infection assay in the presence of Hoechst 33258. Bacteria were infected with T5 phage in the presence of Hoechst 33258, pelleted, supplemented with an excess of untreated bacteria, and then applied to top agar. **С)** Plaque formation efficiency of T5 phage in top agar supplemented with Hoechst 33258.

Section 2. Synthesis and analysis of new bisbenzimidazoles

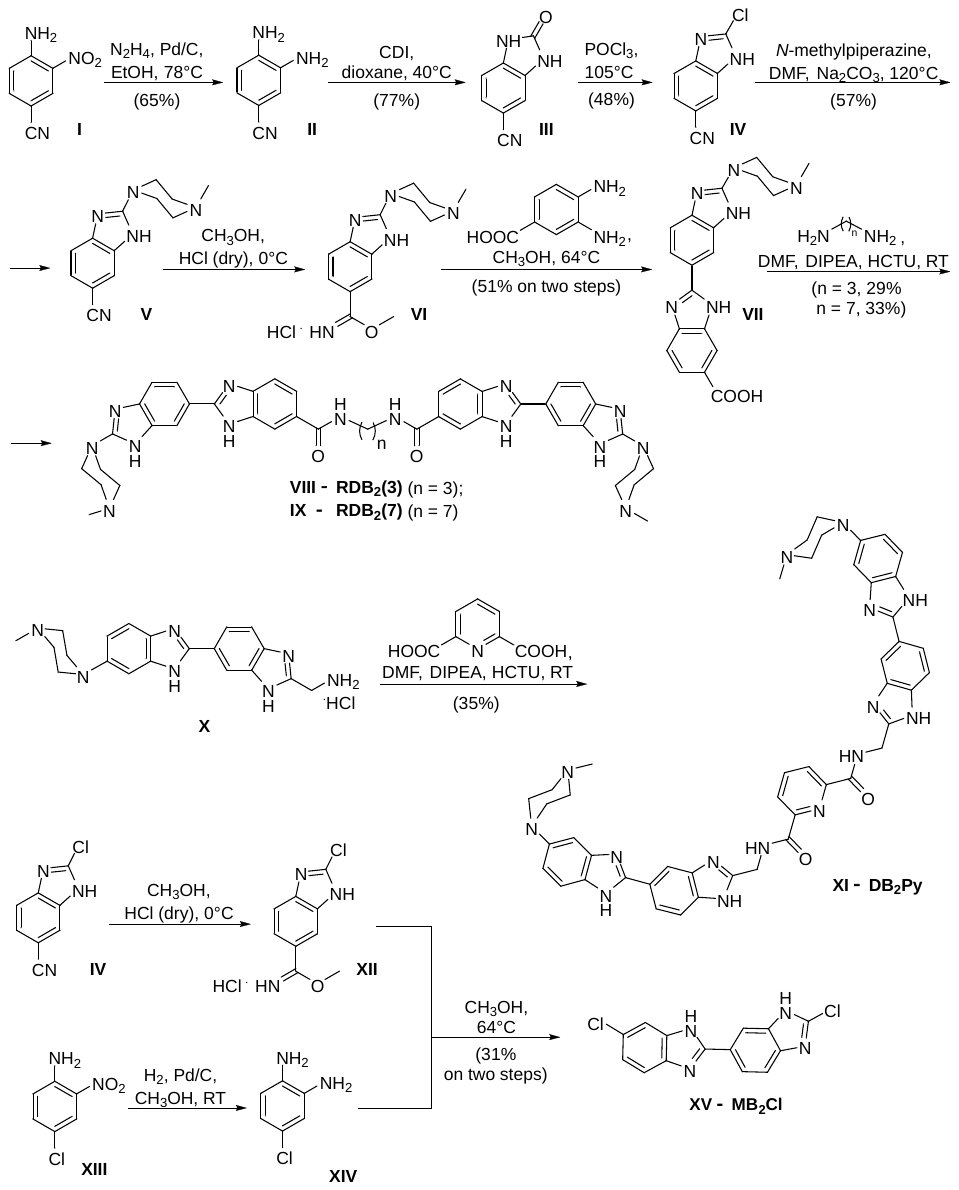

Scheme 1. Synthesis of new bisbenzimidazoles

**General**

All the starting materials were of reagent grade and higher purity, purchased from Acros Organics (Geel, Belgium) and Sigma-Aldrich (Burlington, MA, USA) and used without additional purification, unless otherwise stated. Solvents were dried and purified according to standard procedures; LC-MS grade methanol was purchased from PanReac (Barcelona, Spain) and used without further purification. TLC was performed on Merck Kieselgel 60F_254_ (Darmstadt, Germany) with UV visualization. Column chromatography was performed on silica gel 40–63 μm from Merck in the following systems: A – ethyl acetate/hexane (1:2), B – methylene chloride/methanol/DIPEA (100:20:1), C – ethyl acetate/hexane/ethanol (10:20:1), G – isopropyl alcohol/hydrazine hydrate (6:1), D – methanol/methanolic ammonia (12:1), E – methylene chloride/methanol/acetic acid (16:2:1). The substances in the chromatograms were detected in UV light by absorption at 254 nm, as well as by fluorescence at 365 nm.

^1^H and ^13^C NMR spectra (δ, ppm; *J*, Hz) were registered on an AMX III 300 spectrometer (Bruker BioSpin GmbH, Karlsruhe, Germany) with a working frequency of 300 MHz for ^1^H NMR (Me_4_Si as an internal standard for organic solvents), 75 MHz for ^13^C NMR (with carbon–proton interaction decoupling). High-resolution mass spectra (HRMS) were registered on a micrOTOF-Q II hybrid quadrupole time-of-flight mass spectrometer (Bruker Daltonics GmbH, Bremen, Germany) using electrospray ionization (ESI); the measurements were carried out in positive ion mode. The samples were injected into the mass spectrometer chamber using a syringe injection or from the Agilent 1260 HPLC system (Agilent Technologies, Santa Clara, CA, USA) equipped with an Agilent Poroshell 120 EC-C18 column (3.0 × 50 mm; 2.7 μm) and an identically packed security guard using an autosampler. The samples were in 50% methanol in water (MilliQ ultrapure water; Merck Millipore KGaA, Darmstadt, Germany). The column was eluted with a gradient of acetonitrile (A) concentrations in water (B) with a flow rate of 400 μL/min in the following gradient parameters: 0–15% A for 6 min, 15–85% A for 1.5 min, 85–0% A for 0.1 min, and 0% A for 2.4 min. MALDI-TOF mass spectra were recorded on an AB SCIEX 4800 time-of-flight instrument (AB SCIEX, Framingham, MA, USA) in positive ion recording mode; matrix – 2,5-dihydroxybenzoic acid; N_2_ laser, 337 nm.

**Fluorescence spectra of bisbenzimidazole complexes binding to dsDNA**

The fluorescence spectra were measured in 5 μM solutions of the compounds in 10 mM sodium phosphate buffer (pH 7.5) supplemented with 100 mM potassium chloride upon binding to salmon sperm dsDNA on a Cary Eclipse spectrofluorimeter (Agilent Technologies, Santa Clara, CA, USA) at 25°C. With excitation at 340 nm and a slit width of 5 nm, the fluorescence spectra were recorded in the range of 310-600 nm.

**Experimental procedures**

*3,4-Diaminobenzonitrile* **(II)**

Hydrazine hydrate (11.37 g, 227.1 mmol) was added dropwise to a solution of 4-amino-3-nitrobenzonitrile (**I**, 6.37 g, 39.0 mmol) and Pd/C (416 mg, 3.9 mmol) in 100 mL of ethanol under reflux. After 6 hours the reaction mixture was filtered and the solvent was evaporated. The product was dispersed in 100 mL of diethyl ether, then the precipitate was filtered, air-dried, washed with distilled water (1 × 50 mL) and dried over P_2_O_5_. According to TLC, the product was homogeneous in the system B (R*_f_* = 0.5). Yield 65% (3.37 g).

^1^H NMR (300 MHz, DMSO-*d*_6_), δ: 6.80 (dd, *J* = 7.8, 2.4 Hz, 1H), 6.76 (d, *J* = 2.4 Hz, 1H), 6.55 (d, *J* = 7.8 Hz, 1H), 5.42 (s, 2H), 4.84 (s, 2H).

^13^C NMR (75 MHz, DMSO-*d*_6_), δ: 140.68, 135.22, 122.88, 121.54, 116.23, 113.70, 97.37.

HRМS (*m/z*) C_7_H_7_N_3_: calculated for [M+H]^+^ 134.0780, found 134.0780.

*2-Oxo-2,3-dihydro-1H-benzo[d]imidazole-5-carbonitrile* **(III)**

1,1'-Carbonyldiimidazole (1.95 g, 12.0 mmol) was added to a preheated to 40°C solution of 3,4-diaminobenzonitrile (**II**, 1.50 g, 11.3 mmol) in 15 mL of dioxane. After 5 hours the solvent was evaporated, and distilled water (25 mL) was added to the reaction mixture. The resulting precipitate was filtered, washed with distilled water (2 × 20 mL) and dried over P_2_O_5_. According to TLC the product was homogeneous in the system B (R*_f_* = 0.2). Yield 77% (1.39 g).

^1^H NMR (300 MHz, DMSO-*d*_6_), δ: 11.16 (s, 1H), 11.03 (s, 1H), 7.39 (dd, *J* = 8.1, 1.2 Hz, 1H), 7.31 (d, *J* = 1.2 Hz, 1H), 7.07 (d, *J* = 8.1 Hz, 1H).

^13^C NMR (75 MHz, DMSO-*d*_6_), δ: 155.60, 134.13, 130.38, 126.26, 120.25, 111.83, 109.58, 102.73.

HRМS (*m/z*) C_8_H_5_N_3_O: calculated for [2M+H]^+^ 319.0938, found 319.0944.

*2-Chloro-1H-benzo[d]imidazole-6-carbonitrile* **(IV)**

A solution of 2-oxo-2,3-dihydro-1*H*-benzo[*d*]imidazole-5-carbonitrile (**III**, 894 mg, 5.6 mmol) in 12 mL of phosphorus oxychloride was refluxed for 120 hours. The reaction mixture was poured onto ice. The resulting precipitate was filtered, washed with distilled water (3 × 25 mL) and dried over P_2_O_5_. The product was purified by column chromatography using ethyl acetate/hexane (1:4) system. According to TLC the product was homogeneous in the system A (R*_f_* = 0.6). Yield 48% (474 mg).

^1^H NMR (300 MHz, DMSO-*d*_6_), δ: 8.11 (s, 1H), 7.69 (d, *J* = 8.4 Hz, 1H), 7.65 (d, *J* = 1.5 Hz, 1H), 7.62 (dd, *J* = 8.4, 1.5 Hz, 1H).

^13^C NMR (75 MHz, DMSO-*d*_6_), δ: 142.72, 136.20, 126.55, 120.84, 119.99, 115.75, 105.07.

HRМS (*m/z*) C_8_H_4_^35^ClN_3_: calculated for [2M+H]^+^ 355.0260, found 355.0263.

*2-(4-Methylpiperazin-1-yl)-1H-benzo[d]imidazole-6-carbonitrile* **(V)**

Heated to 120°C solution of 2-chloro-1*H*-benzo[*d*]imidazole-6-carbonitrile (**IV**, 500 mg, 2.8 mmol), *N*-methylpiperazine (282 mg, 5.6 mmol) and sodium carbonate (597 mg, 5.6 mmol) in 5 mL of DMF was stirred for 2 hours. The solvent was evaporated and 10 mL of ethanol was added to the product. Dioxane solution of HCl was added dropwise to the product solution until pH = 4.0. The resulting precipitate was filtered, washed with ethanol (2 × 5 mL) and dried over P_2_O_5_/NaOH. According to TLC data the product is homogeneous in the system A (R*_f_* = 0.1). Yield 57% (389 mg).

^1^H NMR (300 MHz, D_2_O), δ: 7.73 (d, *J* = 1.4 Hz, 1H), 7.63 (dd, *J* = 8.4, 1.4 Hz, 1H), 7.49 (d, *J* = 8.4 Hz, 1H), 4.40 - 4.15 (m, 2H), 3.96 – 3.71 (m, 4H), 3.34 – 3.55 (m, 2H), 3.05 (s, 3H).

^13^C NMR (75 MHz, D_2_O), δ: 150.96, 133.36, 129.69, 128.62, 119.43, 116.00, 112.77, 105.74, 51.79, 43.74, 43.25.

HRМS (*m/z*) C_13_H_15_N_5_: calculated for [M+H]^+^ 242.1401, found 242.1401.

*Methyl 2-(4-methylpiperazin-1-yl)-1H-benzo[d]imidazole-6-carbiimidate hydrochloride* **(VI)**

A precooled to 0°C solution of 2-(4-methylpiperazin-1-yl)-1*H*-benzo[*d*]imidazole-6-carbonitrile **V** (350 mg, 1.45 mmol) in 10 mL of methanol was saturated with dry hydrochloric acid. The reaction mixture was kept sealed for 4 days at +5°C, after that the solvent was evaporated. The product was used immediately in the next step to prepare 2'-(4-methylpiperazin-1-yl)-1*H*,3'*H*-[2,5'-dibenzo[*d*]imidazole]-6-carboxylic acid (**VII**).

*2'-(4-Methylpiperazin-1-yl)-1H,3'H-[2,5'-dibenzo[d]imidazole]-6-carboxylic acid* **(VII)**

A solution of methyl 2-(4-methylpiperazin-1-yl)-1*H*-benzo[*d*]imidazole-6-carbiimidate hydrochloride (**VI**, 300 mg, 0.97 mmol) and 3,4-diaminobenzoic acid (167 mg, 1.1 mmol) in 10 mL of methanol was refluxed for 1 hour. The formed precipitate was filtered off, washed with methanol cooled to 0°C (3 × 10 mL) and dried over P_2_O_5_. According to TLC the product was homogeneous in the system D (R*_f_* = 0.4). Yield 51% (186 mg).

^1^H NMR (300 MHz, DMSO-*d*_6_), δ: 8.03 – 8.00 (m, 1H), 7.93 – 7.89 (m, 1H), 7.60 – 7.48 (m, 2H), 7.18 (d, *J* = 8.2 Hz, 1H), 6.94 (d, *J* = 9.2 Hz, 1H), 3.51 - 3.33 (m, 4H), 2.42 – 2.30 (m, 4H), 2.20 (s, 3H).

^13^C NMR (75 MHz, DMSO-*d*_6_), δ: 174.20, 174.12, 167.78, 148.59, 147.36, 144.30, 132.48, 129.52, 122.46, 120.60, 116.36, 115.66, 112.66, 111.55 111.45, 55.39, 47.06, 46.65.

HRМS (*m/z*) C_20_H_20_N_6_O_2_: calculated for [M+H]^+^ 377.1721, found 377.1716.

*N,N'-(Propan-1,3-diyl)-bis(2'-(4-methylpiperazin-1-yl)-1H,3'H-[2,5'-dibenzo[d]imidazole]-6-carboxamide)* **(VIII – RDB_2_(3))**

DIPEA (75.5 mg, 0.585 mmol) and HCTU (132 mg, 0.318 mmol) were added to a solution of 2'-(4-methylpiperazin-1-yl)-1*H*,3'*H*-[2,5'-dibenzo[*d*]imidazole]-6-carboxylic acid (**VII**, 100 mg, 0.266 mmol) in 5 mL of DMF. The reaction mixture was stirred for 15 min, then propane-1,3-diamine (9.85 mg, 0.133 mmol) was added. After 1 hour the solvent was evaporated. The product was dispersed in 5 ml of chloroform, the precipitate was filtered and dried over P_2_O_5_. The product was purified by column chromatography in a methanol/methanolic ammonia system (25:1). According to TLC the product was homogeneous in the system D (R*_f_* = 0.7). Yield 29% (122 mg).

^1^H NMR (300 MHz, DMSO-*d*_6_ ), δ: 8.53 (t, *J* = 3 Hz, 2H), 8.12 – 8.00 (m, 4H), 7.85 (dd, *J* = 8.4, 1.3 Hz, 2H), 7.68 (dd, *J* = 8.4, 1.3 Hz, 2H), 7.54 (d, *J* = 8.4 Hz, 2H), 7.28 (d, *J* = 7.1 Hz, 2H), 3.61 – 3.53 (m, 8H), 3.39 (m, 4H), 2.49 – 2.40 (m, 8H), 2.24 (s, 3H), 1.80 (m, 2H).

^13^C NMR (75 MHz, DMSO-*d*_6_), δ: 167.55, 162.81, 161.62, 161.34, 157.11, 155.18, 153.34, 151.72, 141.98, 130.56, 128.54, 128.33, 124.70, 120.81, 110.27, 53.67, 36.46, 25.47.

MALDI МS (*m/z*) C_43_H_46_N_14_O_2_: calculated for [M] 790.3928, found – 790.3514.

*N,N'-(Heptane-1,7-diyl)-bis(2'-(4-methylpiperazin-1-yl)-1H,3'H-[2,5'-dibenzo[d]imidazole]-6-carboxamide)* **(IX – RDB_2_(7))**

DIPEA (60.4 mg, 0.468 mmol) and HCTU (106 mg, 0.255 mmol) were added to a solution of 2'-(4-methylpiperazin-1-yl)-1*H*,3'*H*-[2,5'-dibenzo[*d*]imidazole]-6-carboxylic acid (**VII**, 80 mg, 0.213 mmol) in 5 mL of DMF. The reaction mixture was stirred for 15 min, after that heptane-1,7-diamine (13.8 mg, 0.106 mmol) was added. After 1 hour the solvent was evaporated. The product was dispersed in 5 ml of chloroform, the precipitate was filtered and dried over P_2_O_5_. The product was purified by column chromatography in a methanol/methanolic ammonia system (50/1). According to TLC the product was homogeneous in the system D (R*_f_* = 0.8). Yield 33% (30 mg).

^1^H NMR (300 MHz, DMSO-*d*_6_), δ: 8.38 (t, *J* = 5.5 Hz, 2H), 8.06 (d, *J* = 1.5 Hz, 2H), 8.00 (d, *J* = 1.5 Hz, 2H), 7.83 (dd, *J* = 8.4, 1.5 Hz, 2H), 7.69 (dd, *J* = 8.4, 1.5 Hz, 2H), 7.55 (d, *J* = 8.4 Hz, 2H), 7.31 (d, *J* = 8.4 Hz, 2H), 3.59 – 3.50 (m, 8H), 3.33 – 3.25 (m, 4H), 2.48 – 2.40 (m, 8H), 2.24 (s, 6H), 1.64 - 1.52 (m, 4H), 1.43 – 1.33 (m, 6H).

^13^C NMR (75 MHz, DMSO-*d*_6_), δ: 167.18, 157.90, 155.31, 142.22, 141.97, 139.189, 138.81, 128.75, 121.71, 121.55, 119.73, 114.23, 113.05, 110.40, 54.36, 46.27, 29.74, 29.13, 27.08, 19.01.

HRМS (*m/z*) C_47_H_54_N_14_O_2_: calculated for [M+H]^+^ 847.4627, found 847.4617.

MALDI MS (*m/z*) C_47_H_54_N_14_O_2_: calculated for [M] – 846.4554, found – 846.3526.

*N^2^,N^6^-bis((6-(4-methylpiperazin-1-yl)-1H,3’H-[2,5’-dibenzo[d]imidazol]-2’-yl)methyl)pyridine-2,6-dicarboxamide* **(XI – DB_2_Py)**

DIPEA (19.7 mg, 0.152 mmol) and HCTU (34.3 mg, 0.083 mmol) were added to a solution of pyridine-2,6-dicarboxylic acid (11.6 mg, 0.069 mmol) in 5 mL of DMF. The reaction mixture was stirred for 15 min, after that (6-(4-methylpiperazin-1-yl)-1*H*,3’*H*-[2,5’-dibenzo[*d*]imidazol]-2’-yl)methylamine dihydrochloride (**X**, 60 mg, 0.138 mmol) was added. After 1 h the solvent was evaporated. The product was dispersed in 5 ml of chloroform, the precipitate was filtered off and dried over P_2_O_5_. The product was purified by column chromatography in the methylene chloride/methanolic ammonia system (2/1). According to TLC the product was homogeneous in the system E (R*_f_* = 0.7). Yield 35% (83 mg).

^1^H NMR (300 MHz, D_2_O), δ: 8.33 – 8.12 (m, 7H), 7.98 (m, 2H), 7.63 (d, *J* = 9 Hz, 2H), 7.42 (d, *J* = 9 Hz, 2H), 7.01 – 6.88 (м, 2H), 4.85 (d, *J* = 6.0 Hz, 4H), 3.23 – 2.93 (m, 8H), 2.66 - 2.54 (m, 8H), 2.30 (s, 6H).

^13^C NMR (75 MHz, D_2_O), δ: 169.80, 153.74, 152.22, 149.73, 146.24, 144.98, 139.02, 137.71, 137.61, 137.54, 137.48, 137.39, 132.63, 127.86, 125.92, 122.24, 116.43, 112.42, 95.98, 52.44, 46.57, 43.85, 34.46.

HRМS (*m/z*) C_47_H_47_N_15_O_2_: calculated for [M+H]^+^ 854.4110, found 854.4101.

MALDI MS (*m/z*) C_47_H_47_N_15_O_2_: calculated for [M] – 853.4037, found 853.1904.

*2-Chloro-1H-benzo[d]imidazole-6-carbimidate hydrochloride* **(XII)**

A precooled to 0°C solution of 2-chloro-1*H*-benzo[*d*]imidazole-6-carbonitrile (**IV**, 100 mg, 0.563 mmol) in 10 mL of methanol was saturated with dry hydrochloric acid. The reaction mixture was kept sealed for 4 days at +5°C, after that the solvent was evaporated. The reaction product was used immediately in the next step to obtain 2',6-dichloro-1*H*,3'*H*-2,5'-bibenzo[*d*]imidazole (**XV**).

*4-Chlorobenzene-1,2-diamine* **(XIV)**

A suspension of Pd/C (5.97 mg, 0.056 mmol) in 20 mL of methanol was stirred under a hydrogen atmosphere for 10 min, after that 4-chloro-2-nitroaniline (**XIII**, 80 mg, 0.561 mmol) was added and the mixture was hydrogenated for 3 h until hydrogen uptake ceased completely. The reaction mixture was filtered and the filtrate was evaporated. The precipitate was re-evaporated with H_2_O (10 mL) and isopropyl alcohol (10 mL). The product was immediately used for the synthesis of 2',6-dichloro-1*H*,3'*H*-2,5'-bibenzo[*d*]imidazole (***XV***).

*2',6-Dichloro-1H,3'H-2,5'-bibenzo[d]imidazole* **(XV – MB_2_Cl)**

A solution of methyl 2-chloro-1*H*-benzo[*d*]imidazole-6-carbimidate hydrochloride (**XII**, 100 mg, 0.406 mmol) and 4-chlorobenzene-1,2-diamine (**XIV**, 58 mg, 0.406 mmol) in 20 mL of methanol was refluxed for 6 hours. The solvent was evaporated from the reaction mixture, the product was purified by column chromatography using an ethyl acetate/methanol gradient. According to TLC the product was homogeneous in the system D (R*_f_* = 0.8). Yield 31% (38 mg).

^1^H NMR (300 MHz, DMSO-*d*_6_), δ: 8.31 (d, *J* = 1.6 Hz, 1H), 8.23 (d, *J* = 1.5 Hz, 1H), 8.13 (dd, *J* = 8.4, 1.7 Hz 1H), 7.93 (dd, *J* = 8.5, 1.6 Hz, 1H), 7.74 (d, *J* = 8.5 Hz, 1H), 7.58 (d, *J* = 8.4 Hz, 1H).

^13^C NMR (75 MHz, DMSO-*d*_6_), δ: 167.80, 148.39, 147.30, 144.34, 132.27, 129.31, 128.40, 122.37, 120.66, 116.28, 115.71, 112.58, 111.66, 111.21.

**^1^H NMR spectrum of 3,4-diaminobenzonitrile (II)**

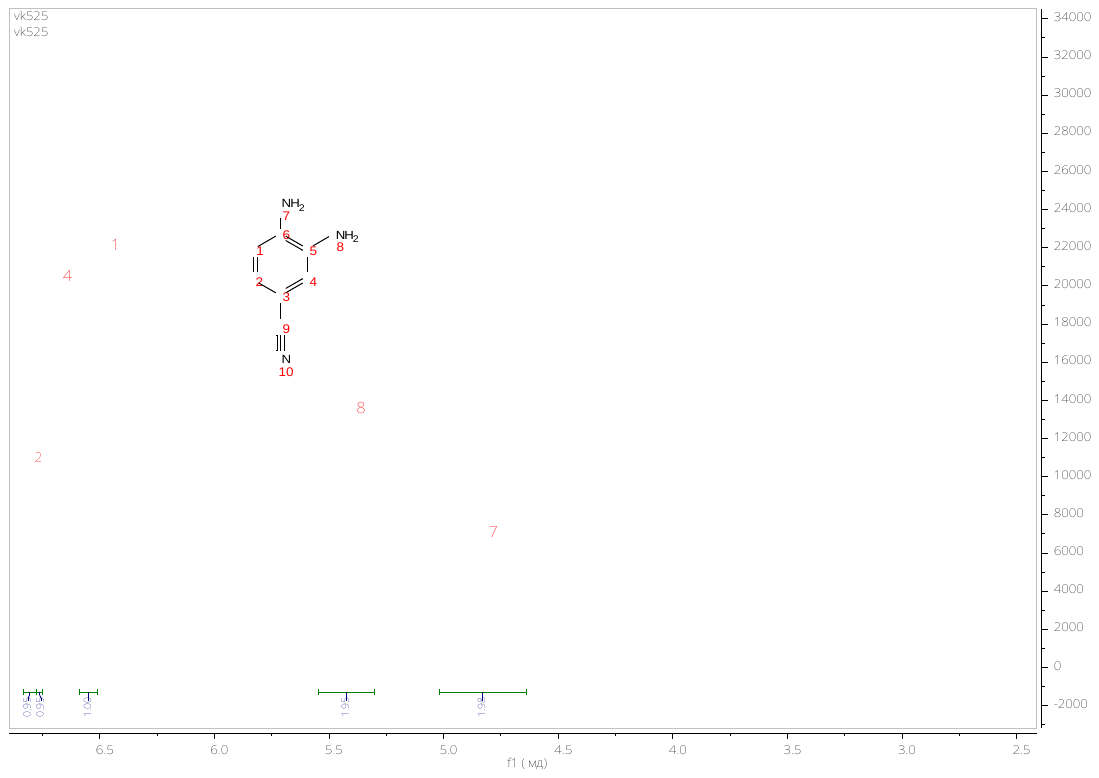

**^13^C NMR spectrum of 3,4-diaminobenzonitrile (II)**

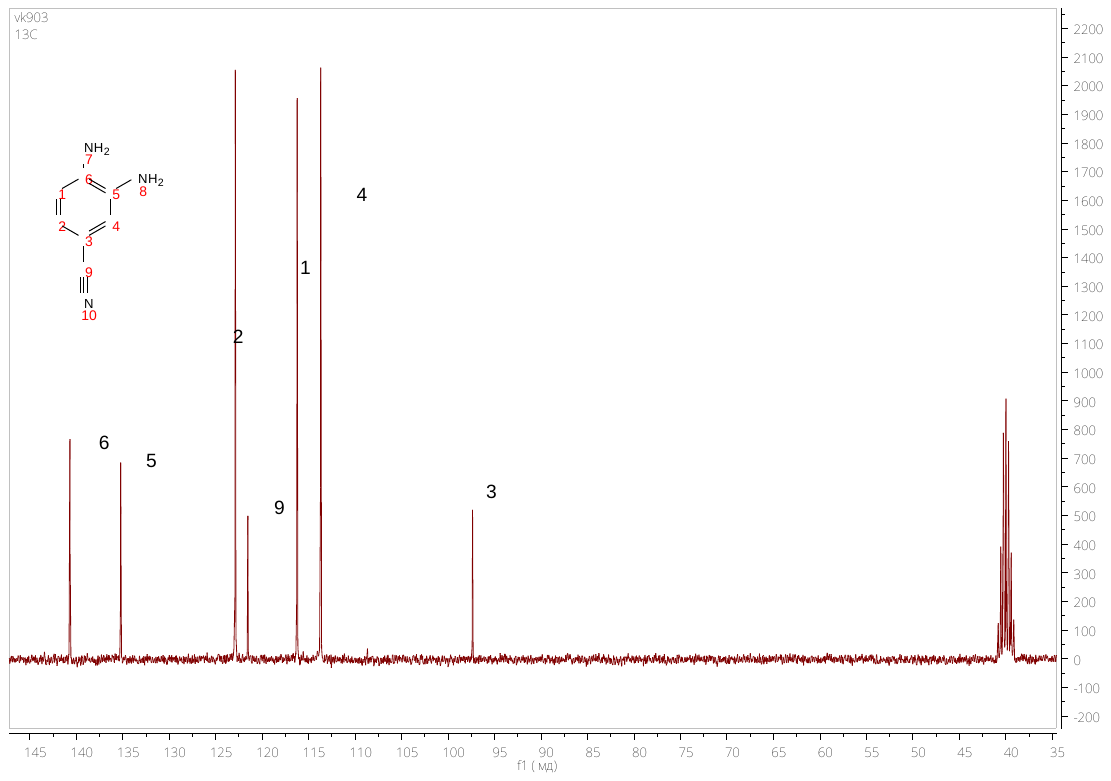

**HRMS spectrum of 3,4-diaminobenzonitrile (II)**

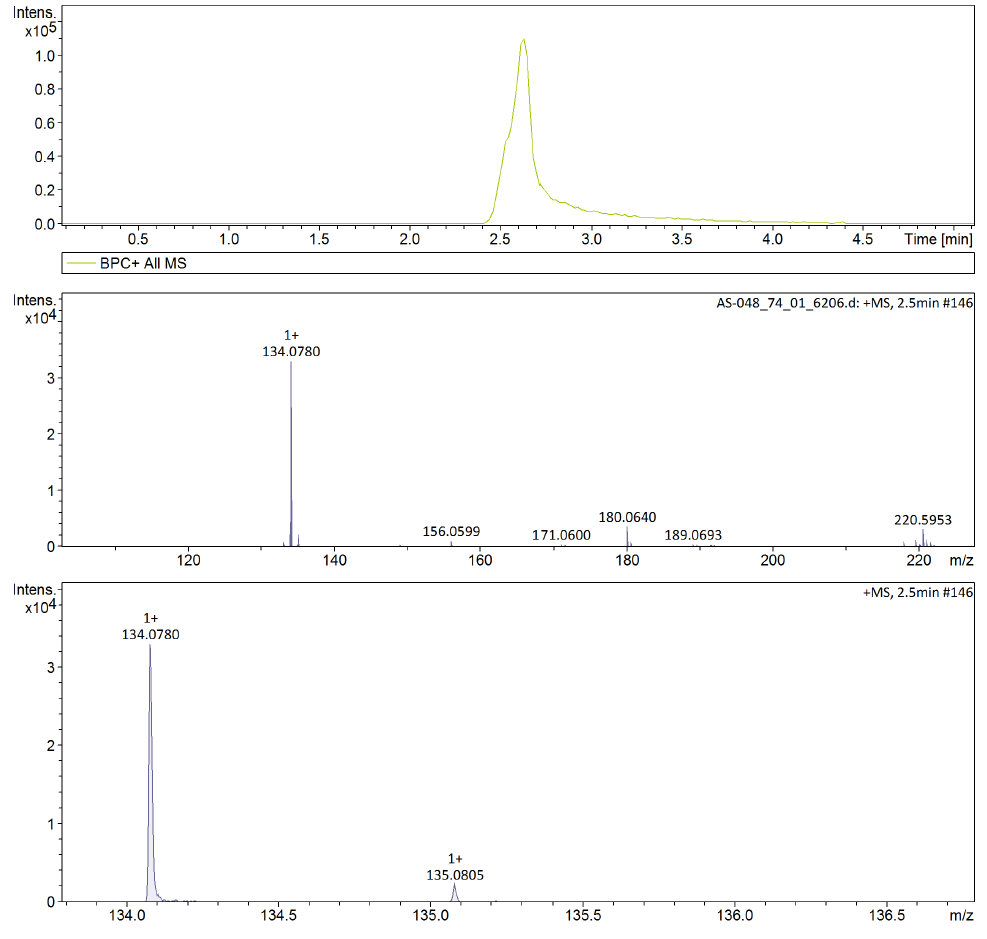

**^1^H NMR spectrum of 2-oxo-2,3-dihydro-1*H*-benzo[*d*]imidazole-5-carbonitrile (III)**

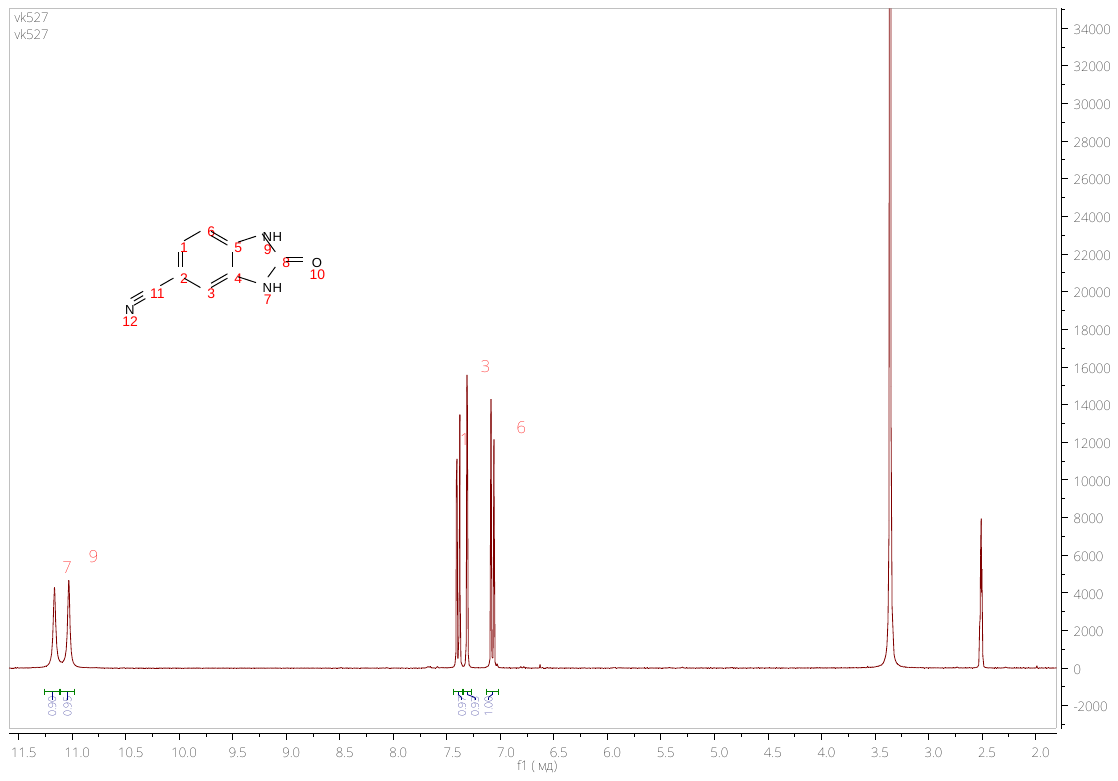

**^13^C NMR spectrum of 2-oxo-2,3-dihydro-1*H*-benzo[*d*]imidazole-5-carbonitrile (III)**

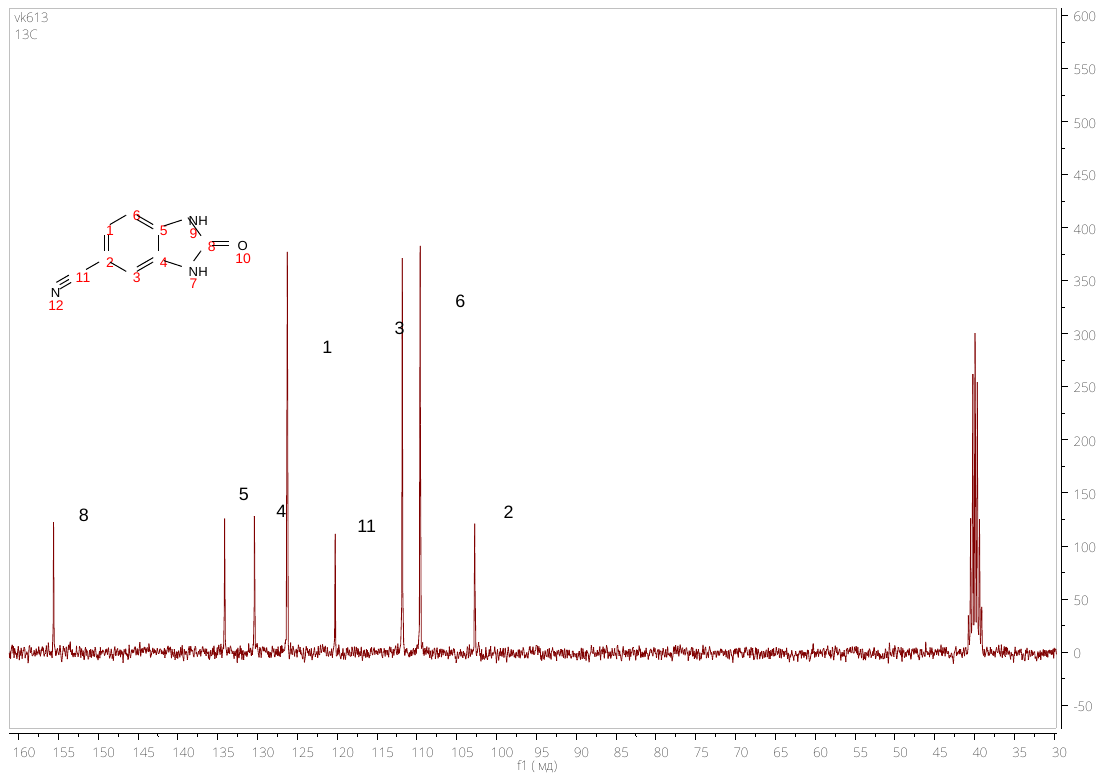

**HRMS spectrum of 2-oxo-2,3-dihydro-1*H*-benzo[*d*]imidazole-5-carbonitrile (III)**

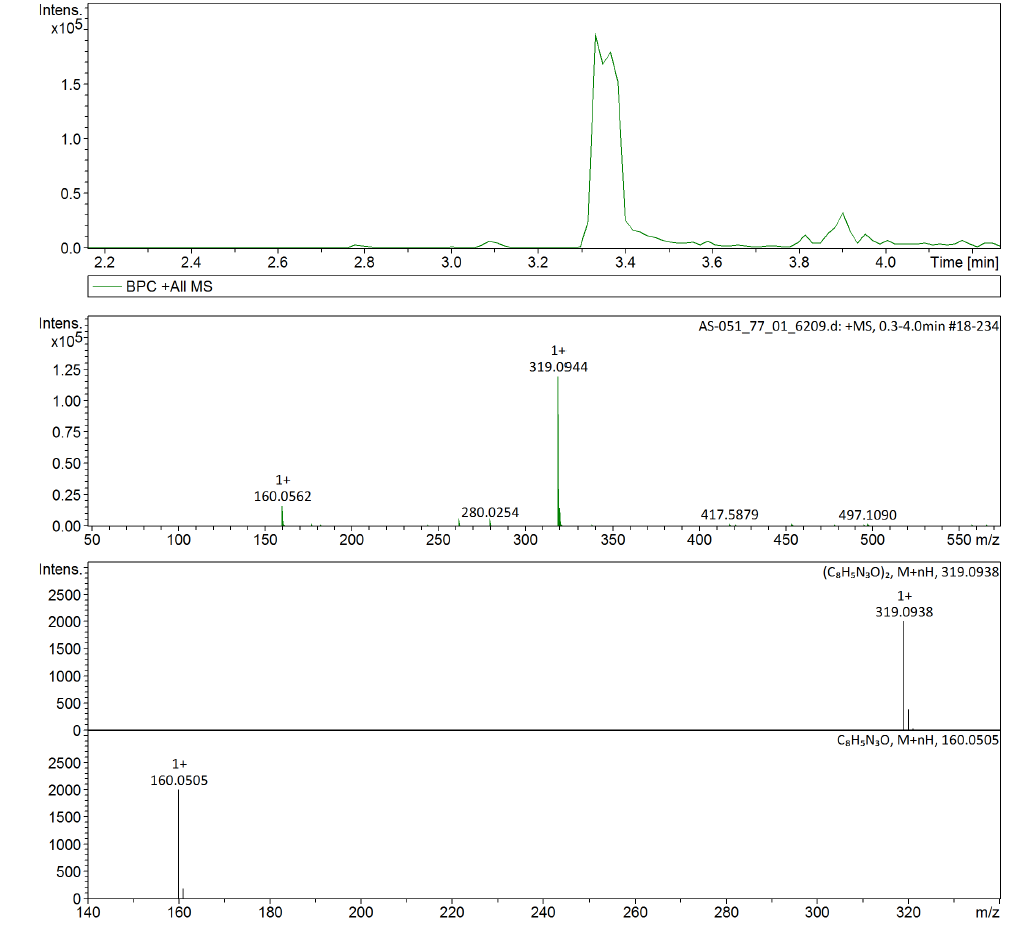

**^1^H NMR spectrum of 2-chloro-1*H*-benzo[*d*]imidazole-6-carbonitrile (IV)**

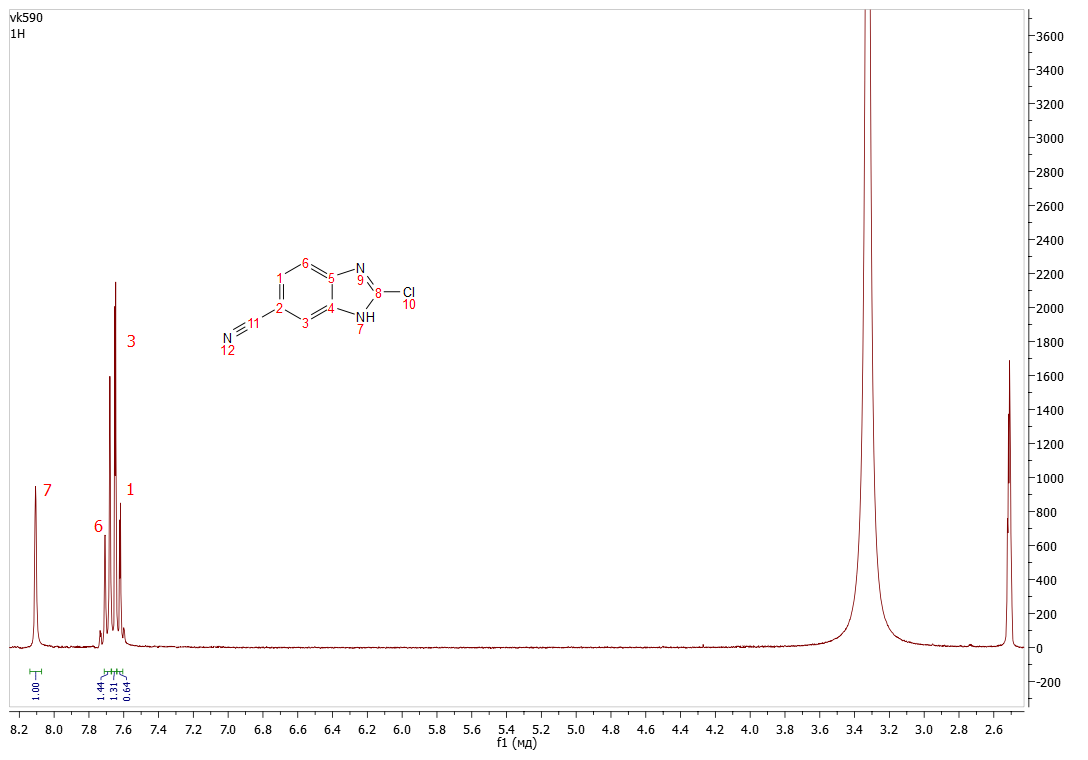

**^13^C NMR spectrum of 2-chloro-1*H*-benzo[*d*]imidazole-6-carbonitrile (IV)**

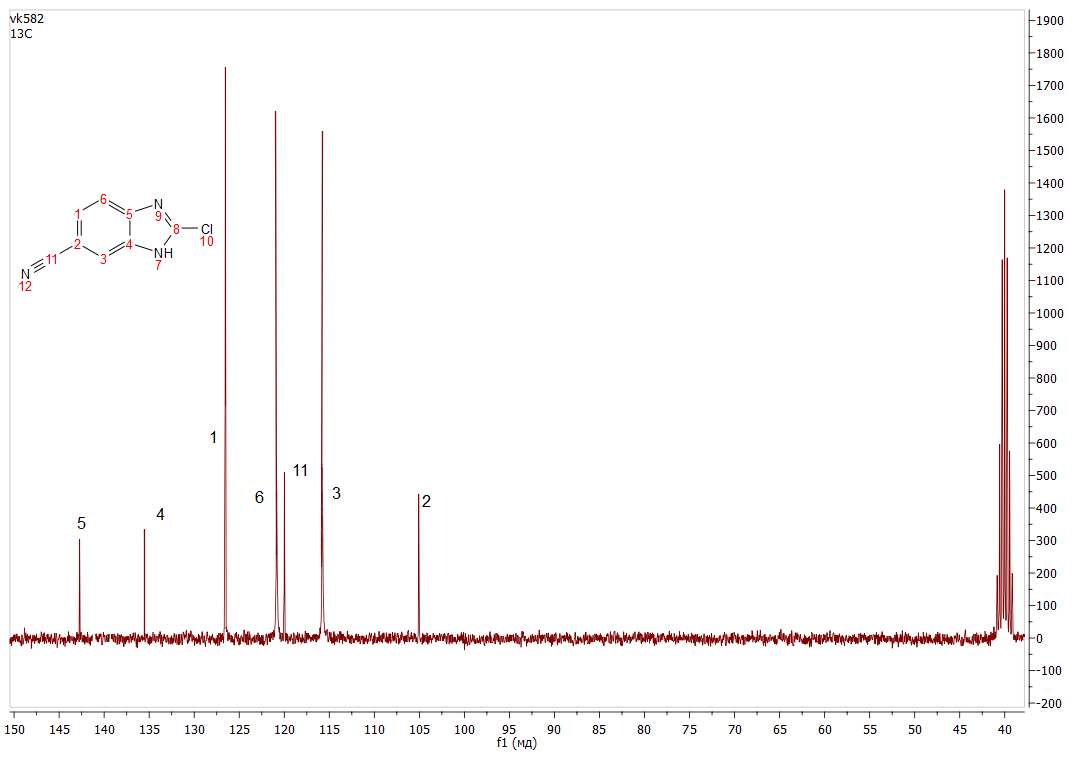

**HRMS spectrum of 2-chloro-1*H*-benzo[*d*]imidazole-6-carbonitrile (IV)**

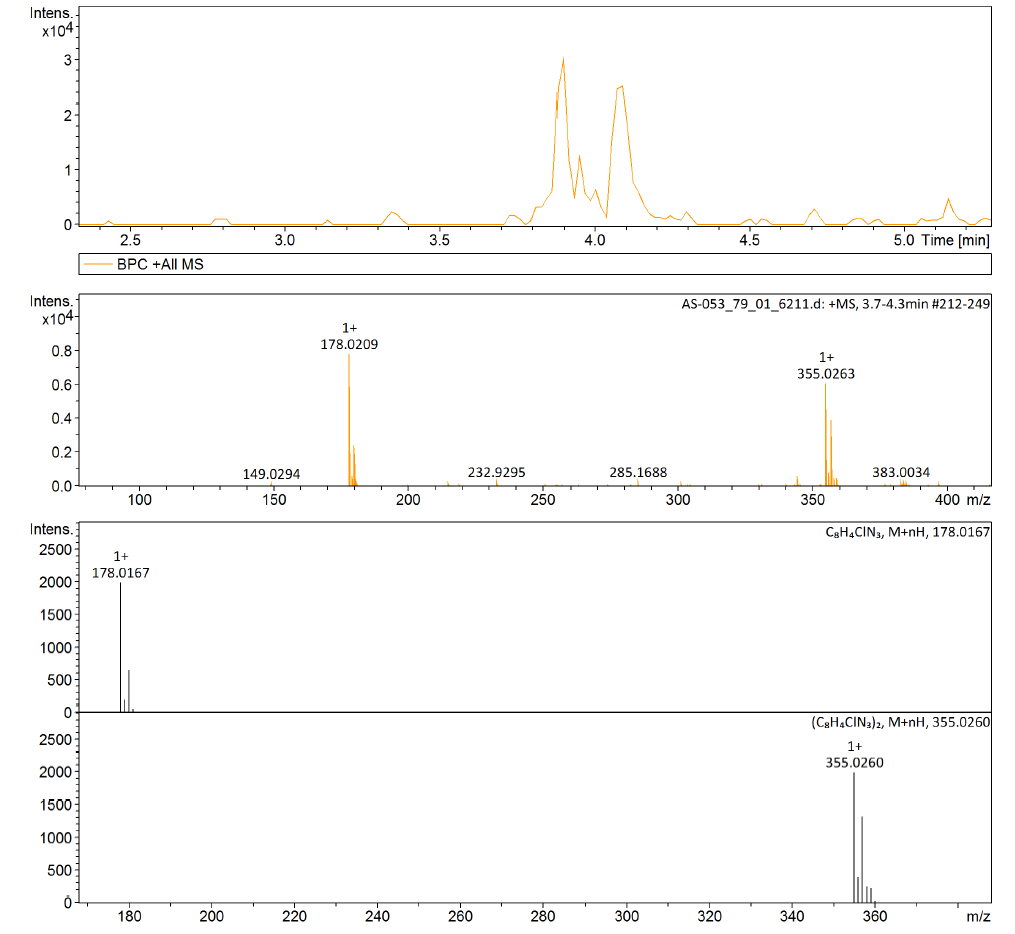

**^1^H NMR spectrum of 2-(4-methylpiperazin-1-yl)-1*H*-benzo[*d*]imidazole-6-carbonitrile (V)**

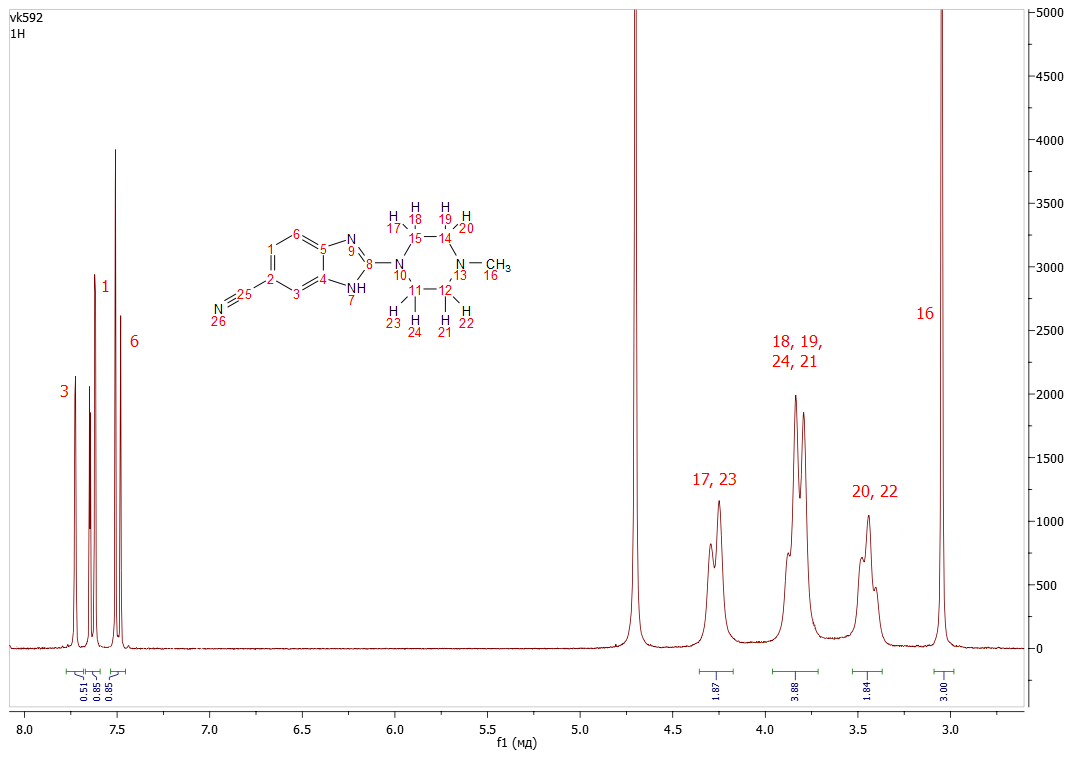

**^13^C NMR spectrum of 2-(4-methylpiperazin-1-yl)-1*H*-benzo[*d*]imidazole-6-carbonitrile (V)**

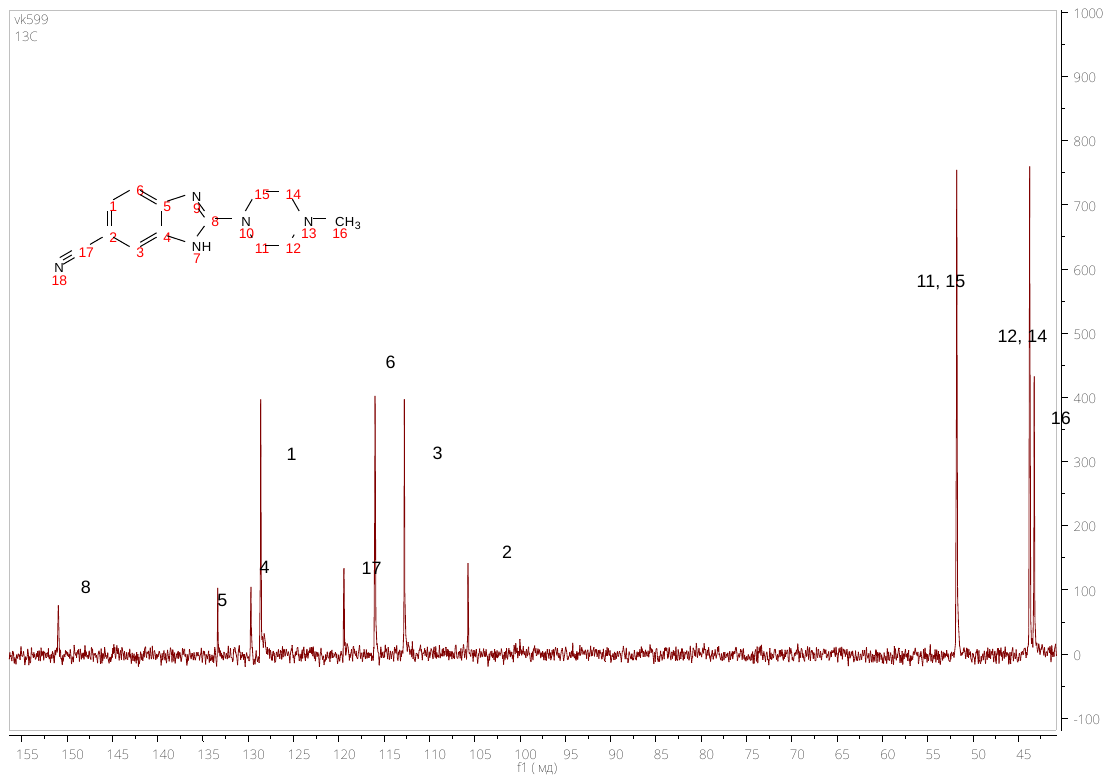

**HRMS spectrum of 2-(4-methylpiperazin-1-yl)-1*H*-benzo[*d*]imidazole-6-carbonitrile (V)**

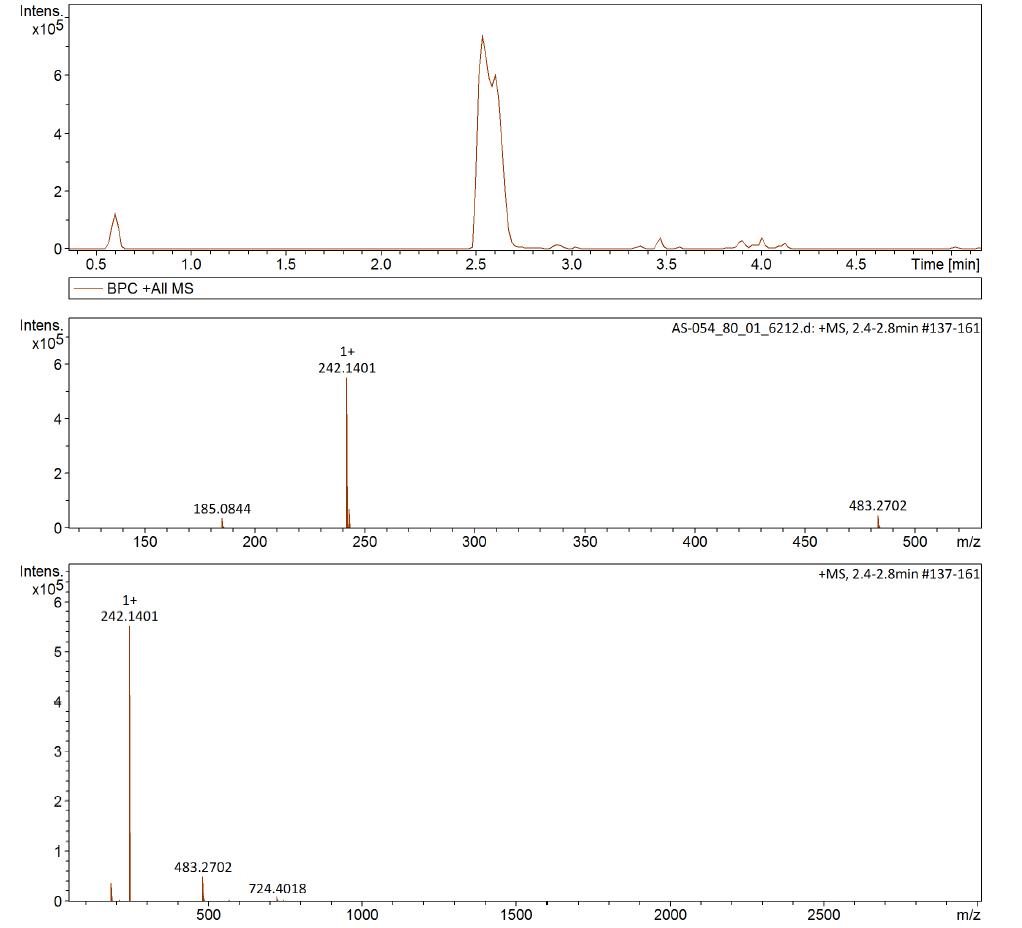

**^1^H NMR spectrum of 2'-(4-methylpiperazin-1-yl)-1*H*,3'*H*-[2,5'-dibenzo[*d*]imidazole]-6-carboxylic acid (VII)**

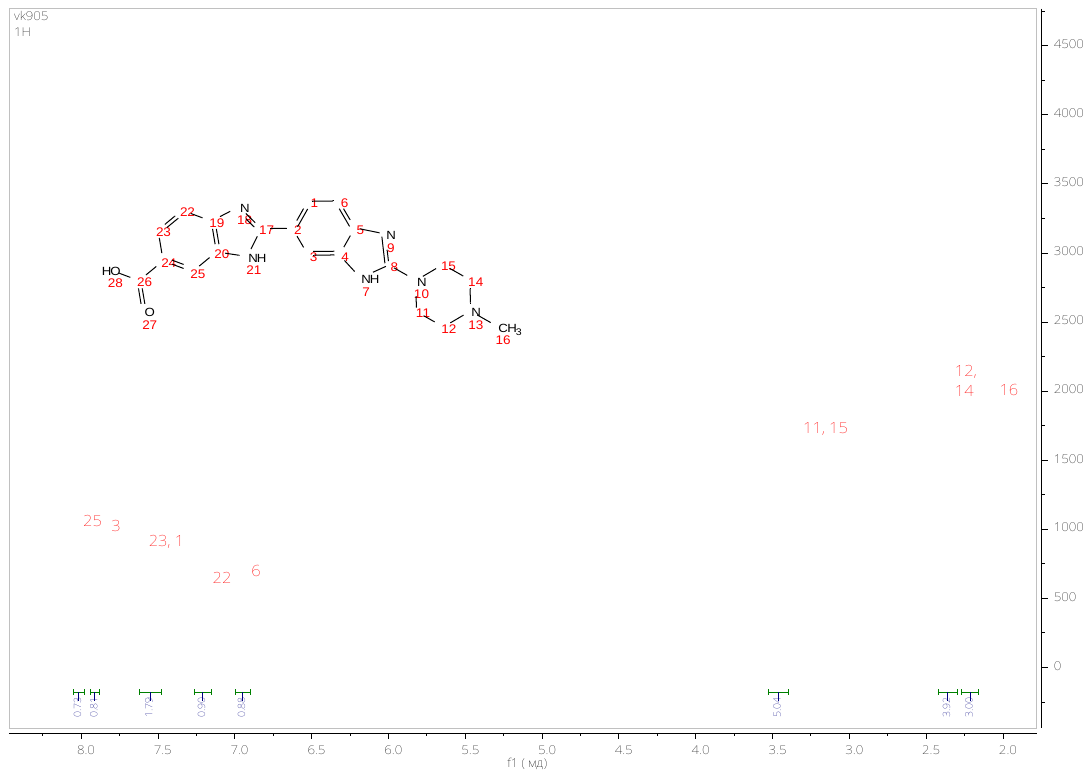

**^13^C NMR spectrum of 2'-(4-methylpiperazin-1-yl)-1*H*,3'*H*-[2,5'-dibenzo[*d*]imidazole]-6-carboxylic acid (VII)**

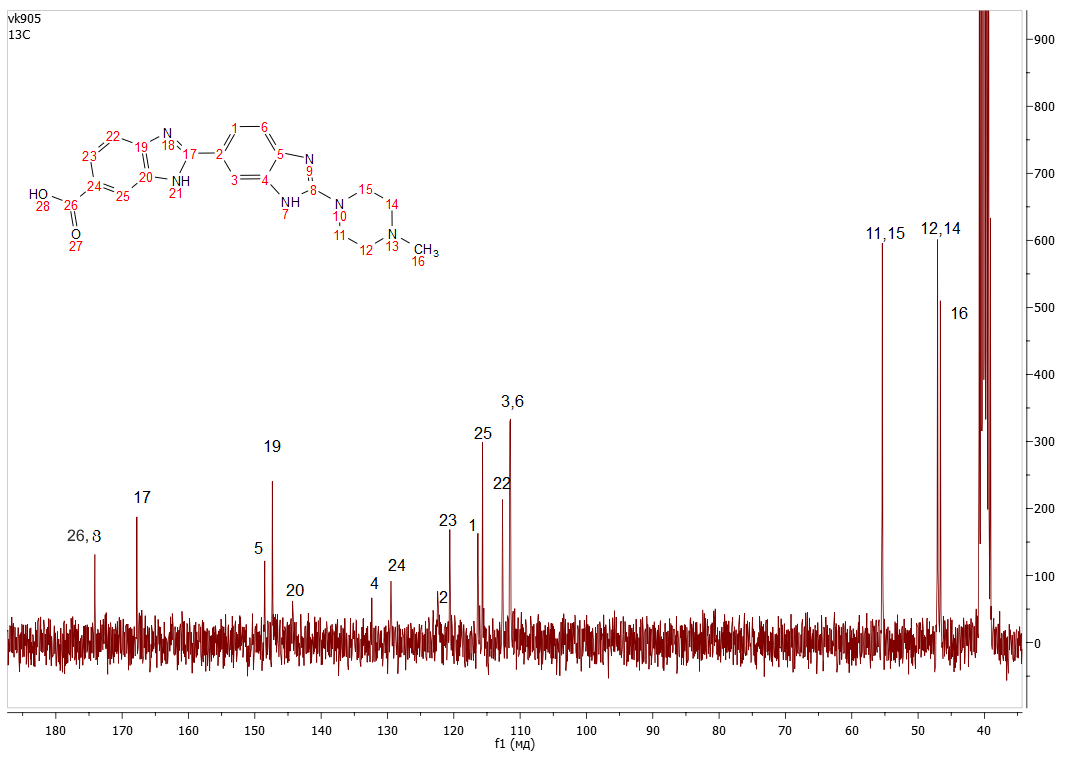

**HRMS spectrum of 2'-(4-methylpiperazin-1-yl)-1*H*,3'*H*-[2,5'-dibenzo[*d*]imidazole]-6-carboxylic acid (VII)**

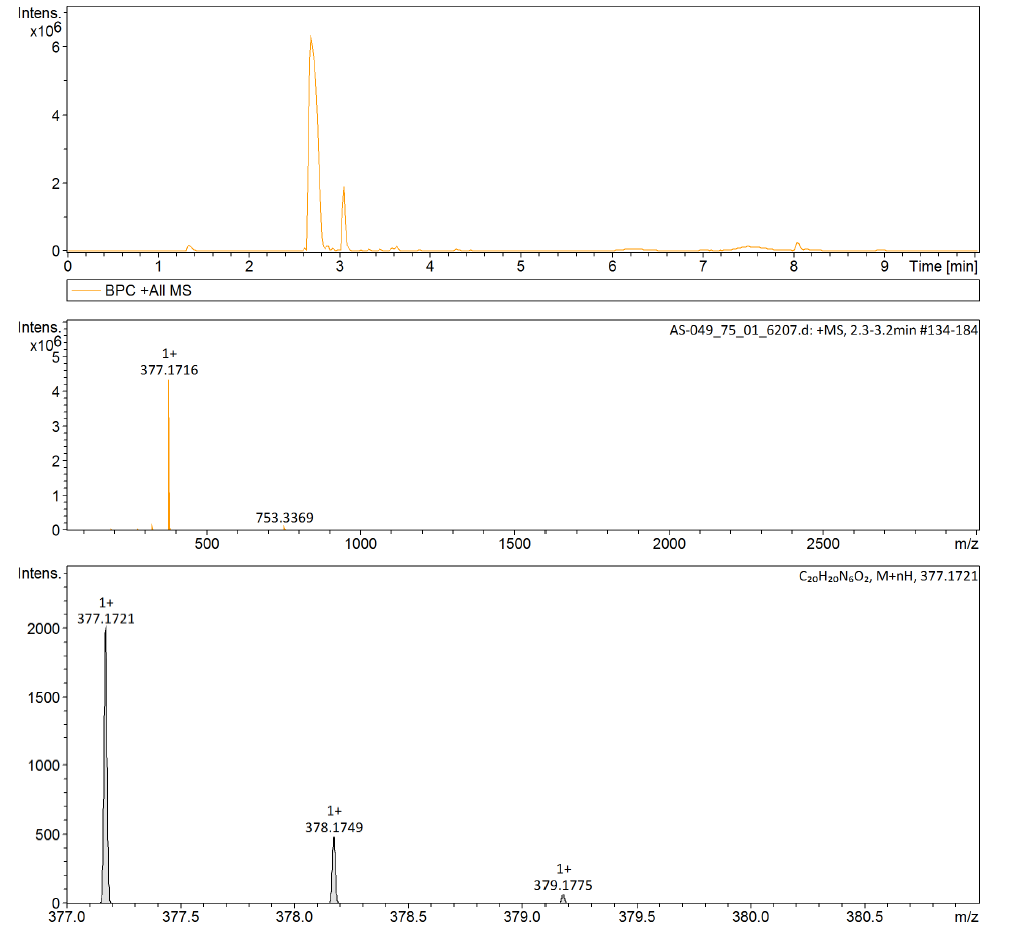

**^1^H NMR spectrum of N,N'-(propan-1,3-diyl)bis(2'-(4-methylpiperazin-1-yl)-1*H*,3'*H*-[2,5'-dibenzo[*d*]imidazole]-6-carboxamide) (VIII – RDB_2_(3))**

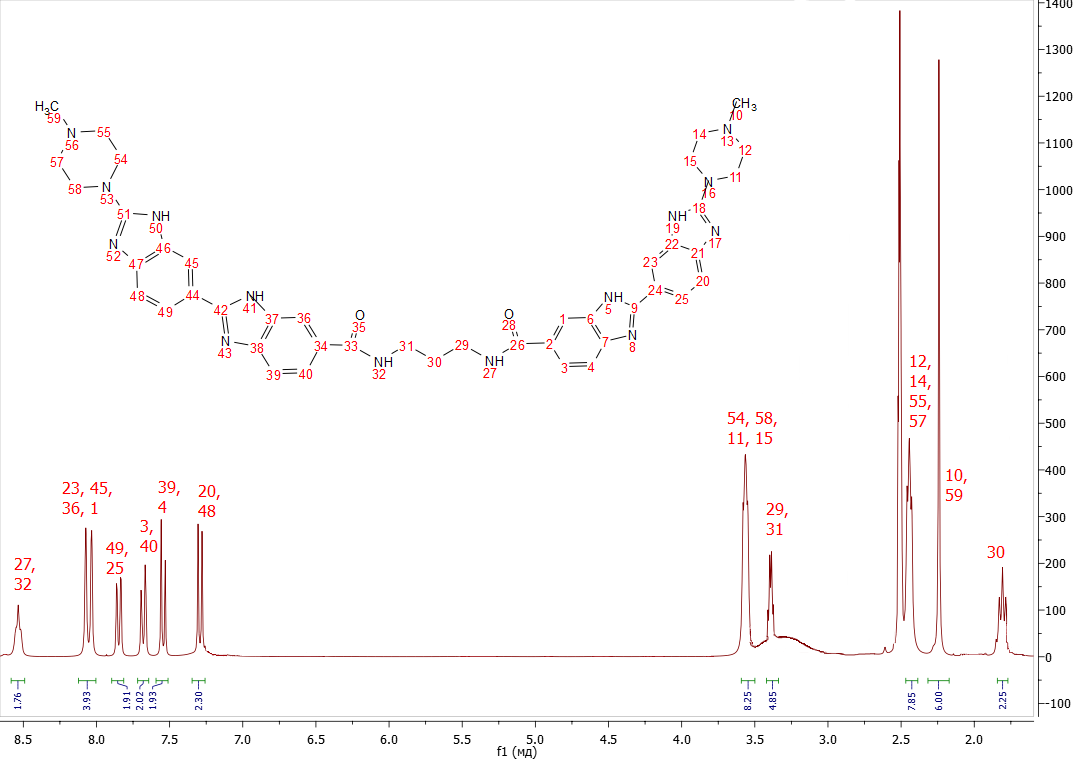

**^13^C NMR spectrum of N,N'-(propan-1,3-diyl)bis(2'-(4-methylpiperazin-1-yl)-1*H*,3'*H*-[2,5'-dibenzo[*d*]imidazole]-6-carboxamide) (VIII – RDB_2_(3))**

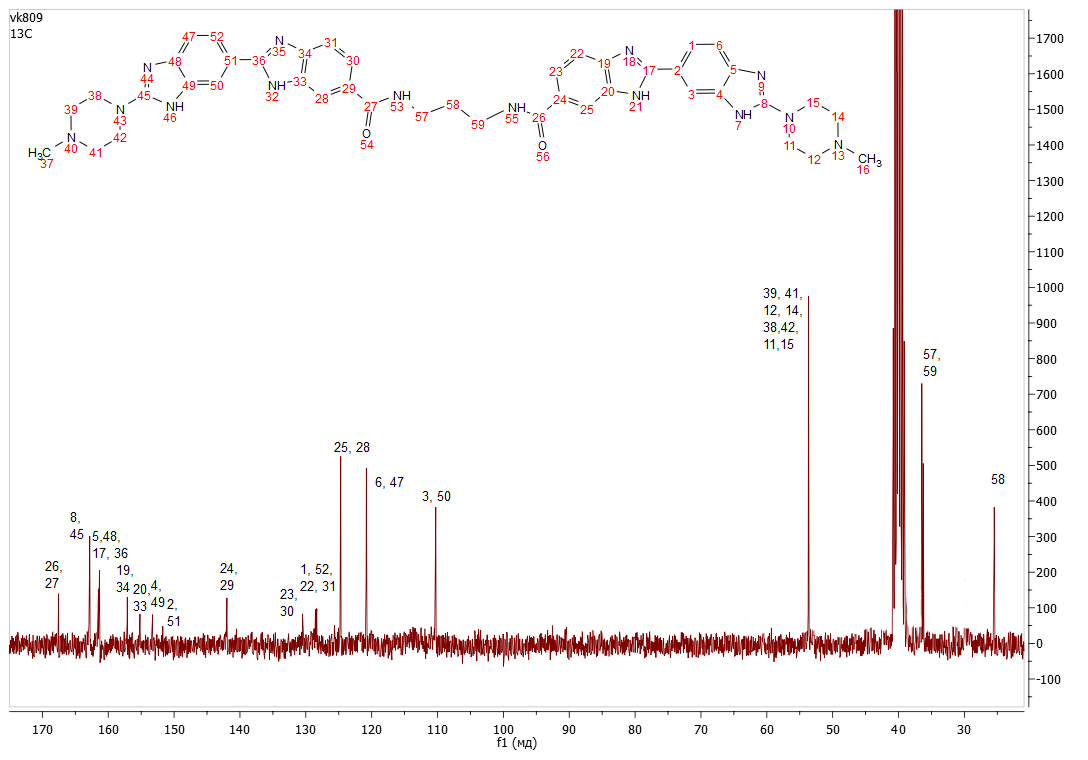

**MALDI MS spectrum of N,N'-(propan-1,3-diyl)bis(2'-(4-methylpiperazin-1-yl)-1*H*,3'*H*-[2,5'-dibenzo[*d*]imidazole]-6-carboxamide) (VIII – RDB_2_(3))**

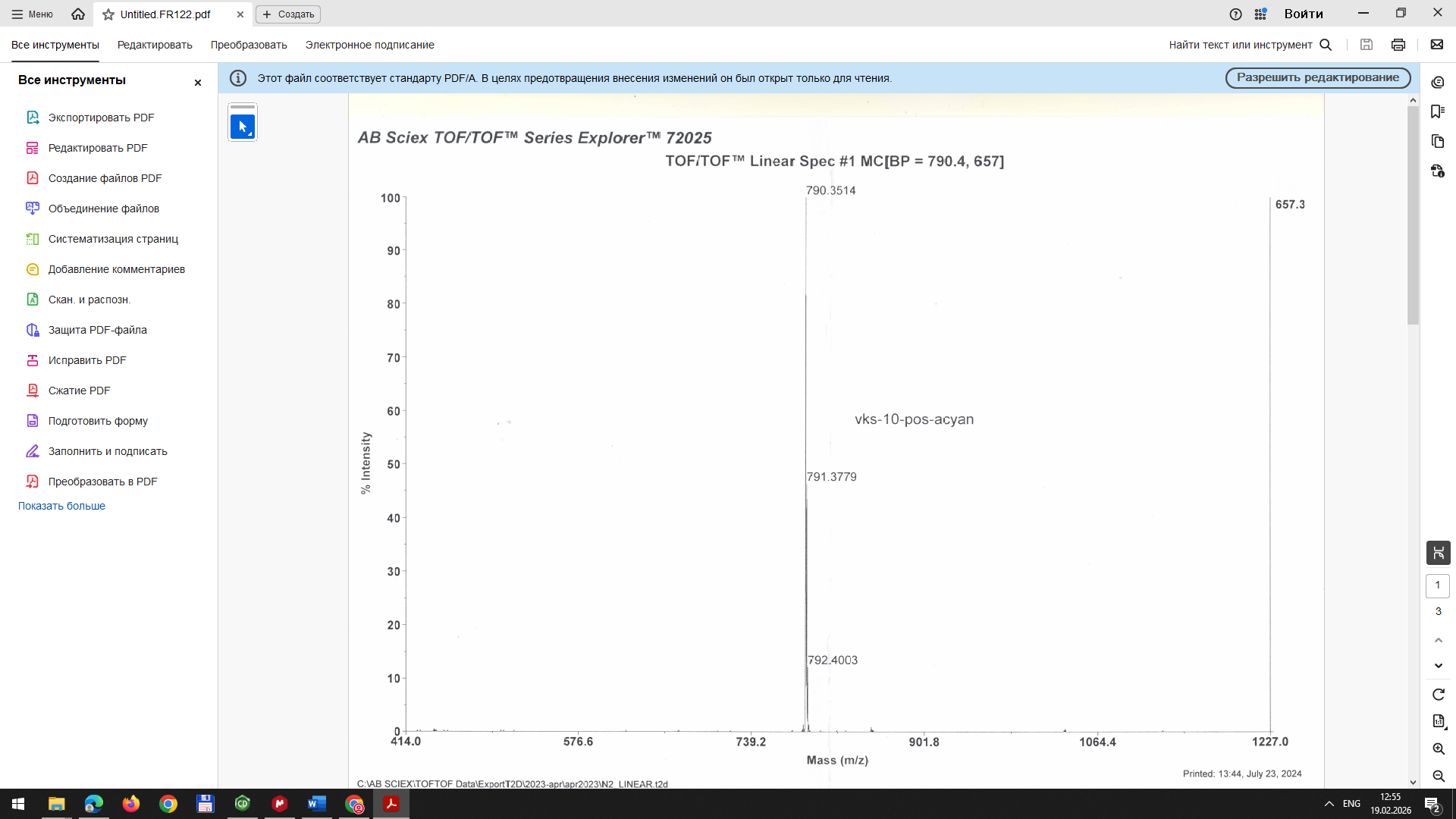

**^1^H NMR spectrum of N,N'-(heptan-1,7-diyl)bis(2'-(4-methylpiperazin-1-yl)-1*H*,3'*H*-[2,5'-dibenzo[*d*]imidazole]-6-carboxamide) (IX – RDB_2_(7))**

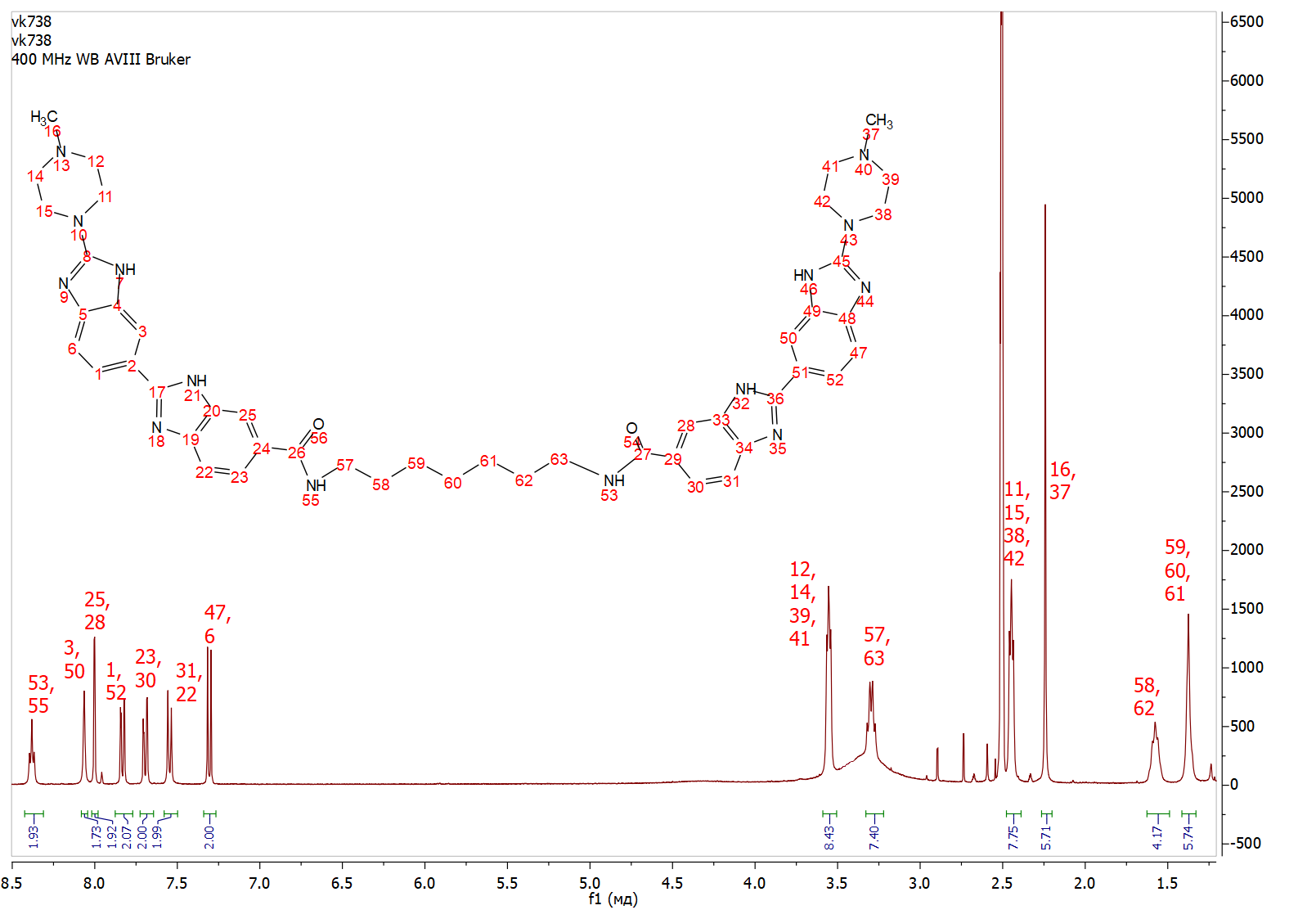

**^13^C NMR spectrum of N,N'-(heptan-1,7-diyl)bis(2'-(4-methylpiperazin-1-yl)-1*H*,3'*H*-[2,5'-dibenzo[*d*]imidazole]-6-carboxamide) (IX – RDB_2_(7))**

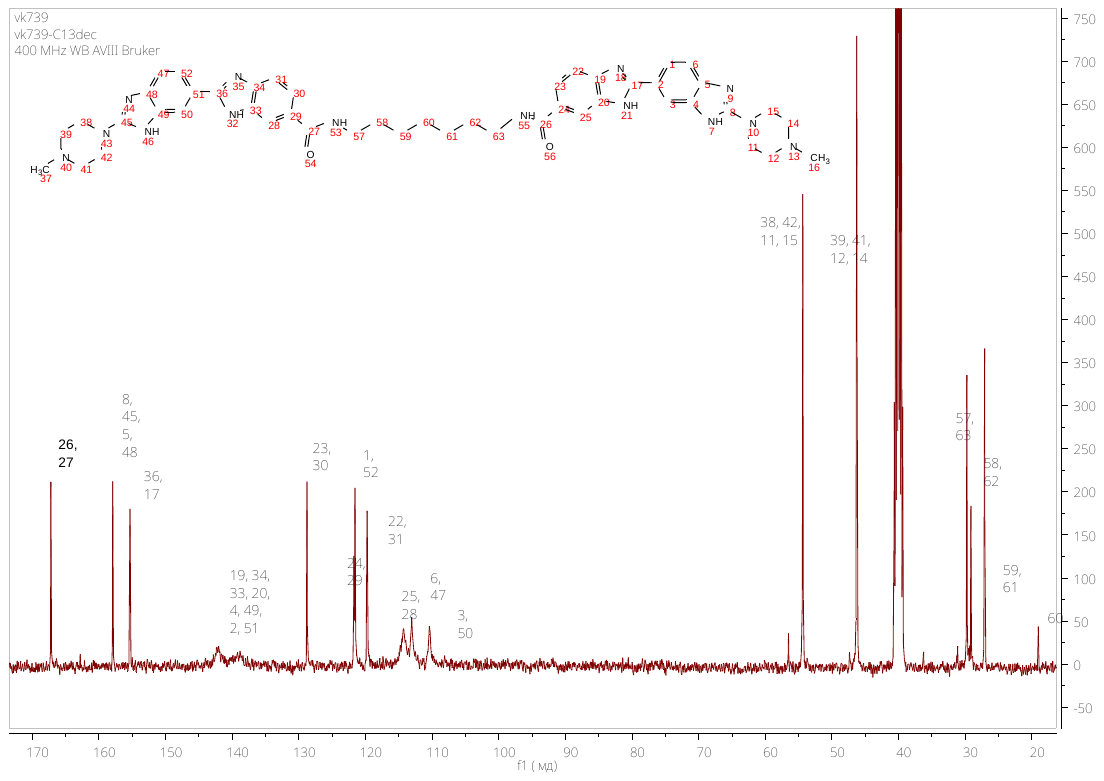

**MALDI MS spectrum of N,N'-(heptan-1,7-diyl)bis(2'-(4-methylpiperazin-1-yl)-1*H*,3'*H*-[2,5'-dibenzo[*d*]imidazole]-6-carboxamide) (IX – RDB_2_(7))**

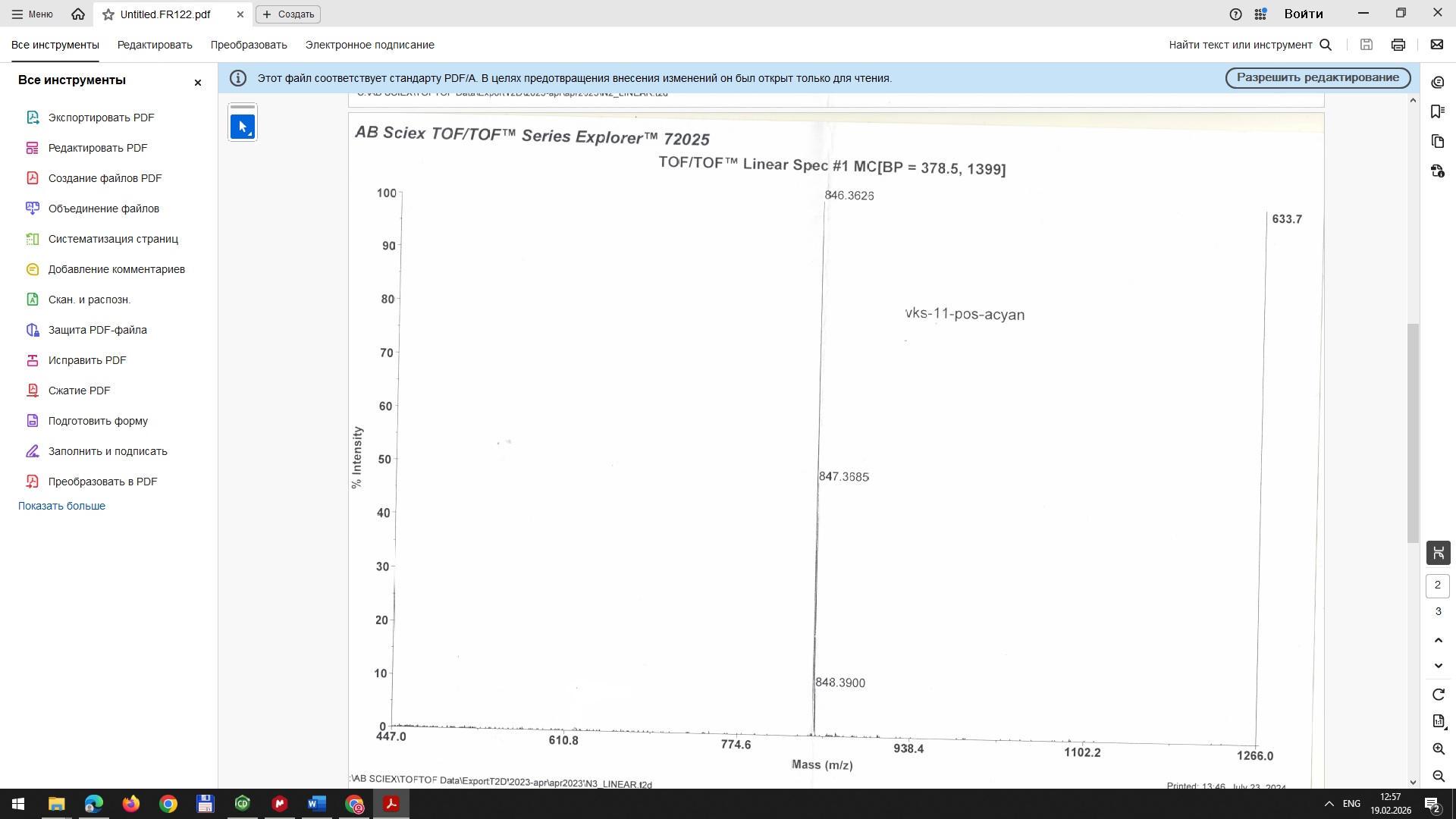

**HRMS spectrum of N,N'-(heptan-1,7-diyl)bis(2'-(4-methylpiperazin-1-yl)-1*H*,3'*H*-[2,5'-dibenzo[*d*]imidazole]-6-carboxamide) (IX – RDB_2_(7))**

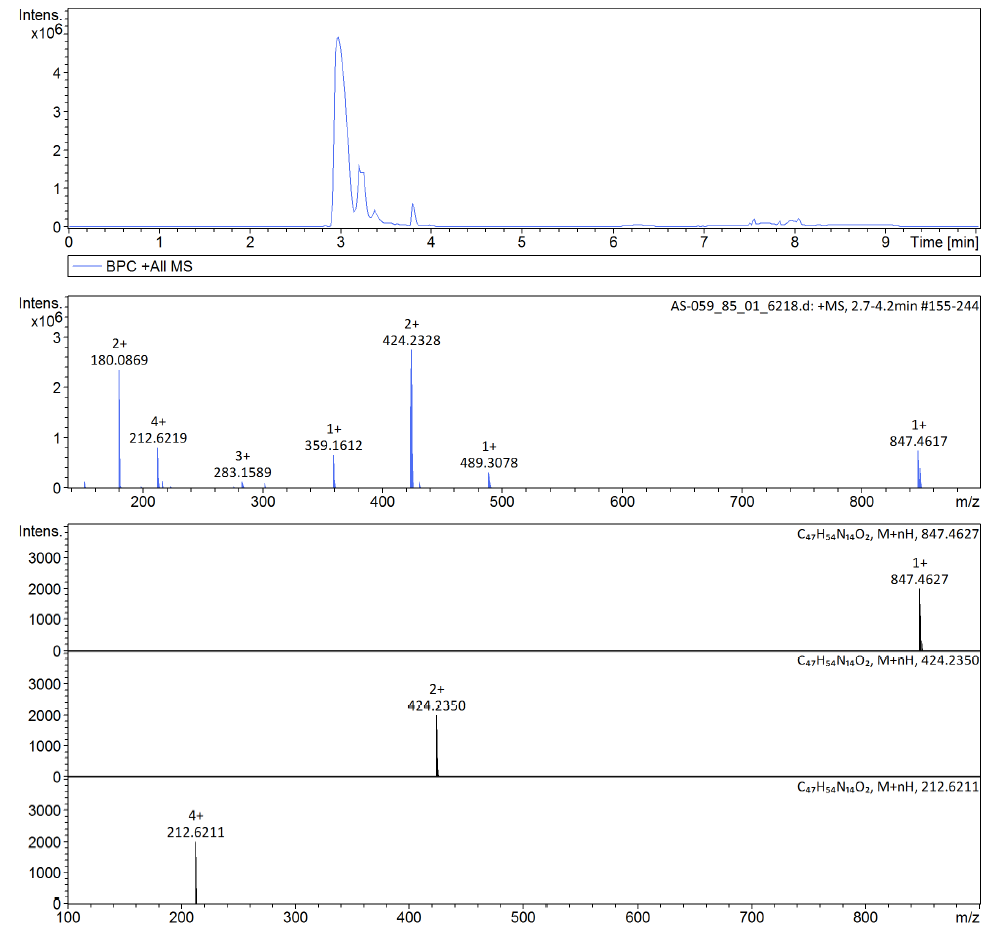

**^1^H NMR spectrum of N^2^,N^6^-bis((6-(4-methylpiperazin-1-yl)-1*H*,3'*H*-[2,5'-dibenzo[*d*]imidazole]-2'-yl)methyl)pyridine-2,6-dicarboxamide (XI - DB_2_Py)**

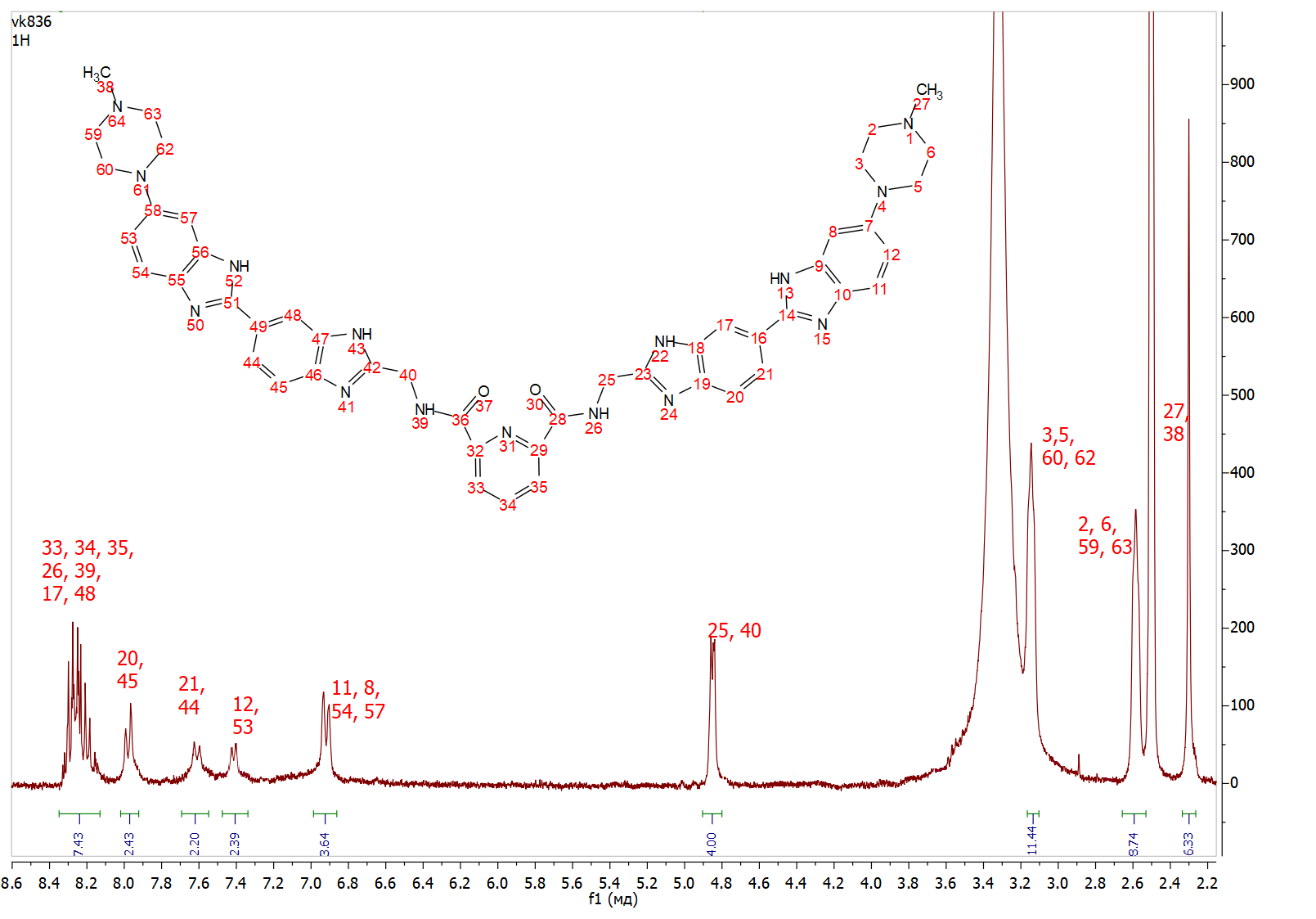

**^13^C NMR spectrum of N^2^,N^6^-bis((6-(4-methylpiperazin-1-yl)-1*H*,3'*H*-[2,5'-dibenzo[*d*]imidazole]-2'-yl)methyl)pyridine-2,6-dicarboxamide (XI - DB_2_Py)**

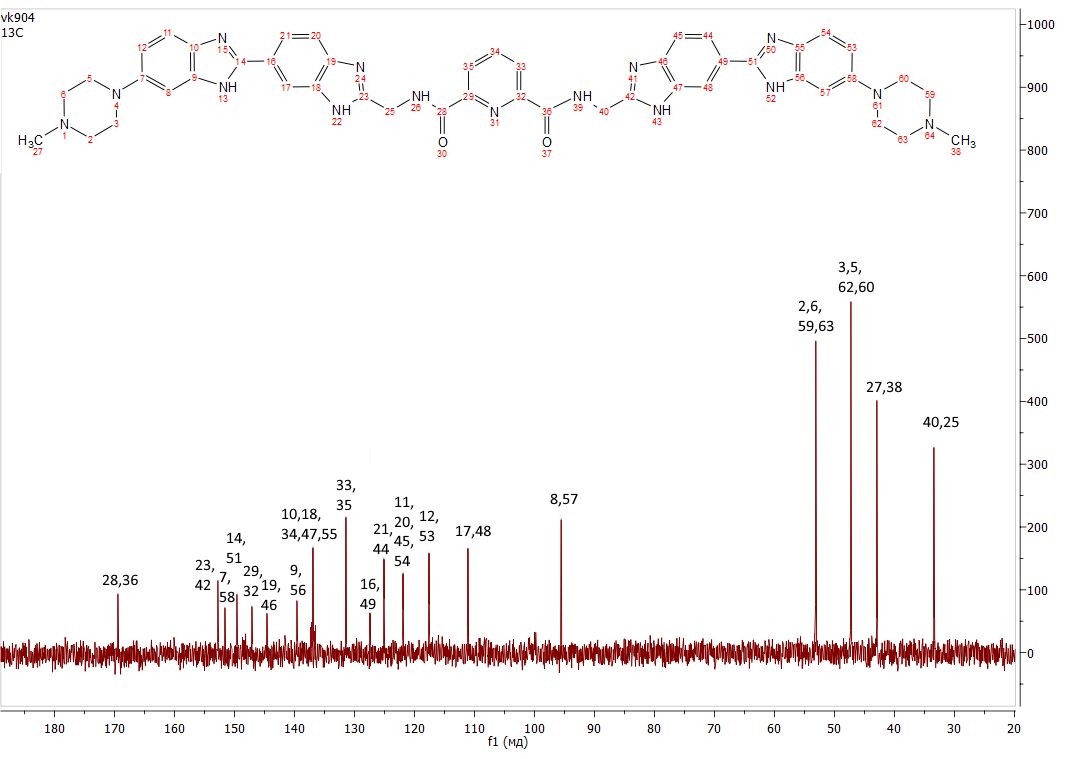

**MALDI MS spectrum of N^2^,N^6^-bis((6-(4-methylpiperazin-1-yl)-1*H*,3'*H*-[2,5'-dibenzo[*d*]imidazole]-2'-yl)methyl)pyridine-2,6-dicarboxamide (XI - DB_2_Py)**

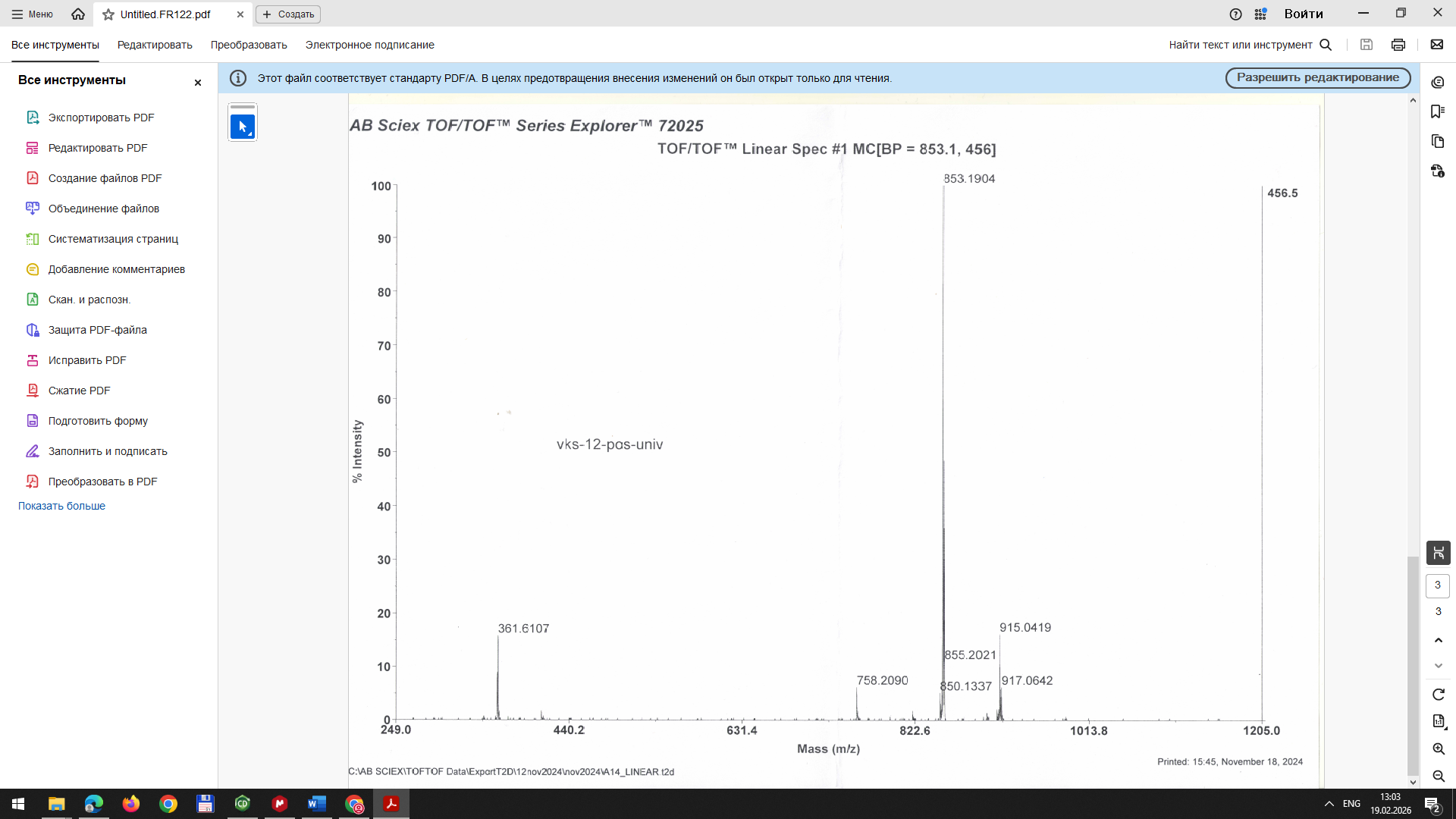

**HRMS spectrum of N^2^,N^6^-bis((6-(4-methylpiperazin-1-yl)-1*H*,3'*H*-[2,5'-dibenzo[*d*]imidazole]-2'-yl)methyl)pyridine-2,6-dicarboxamide (XI – DB_2_Py)**

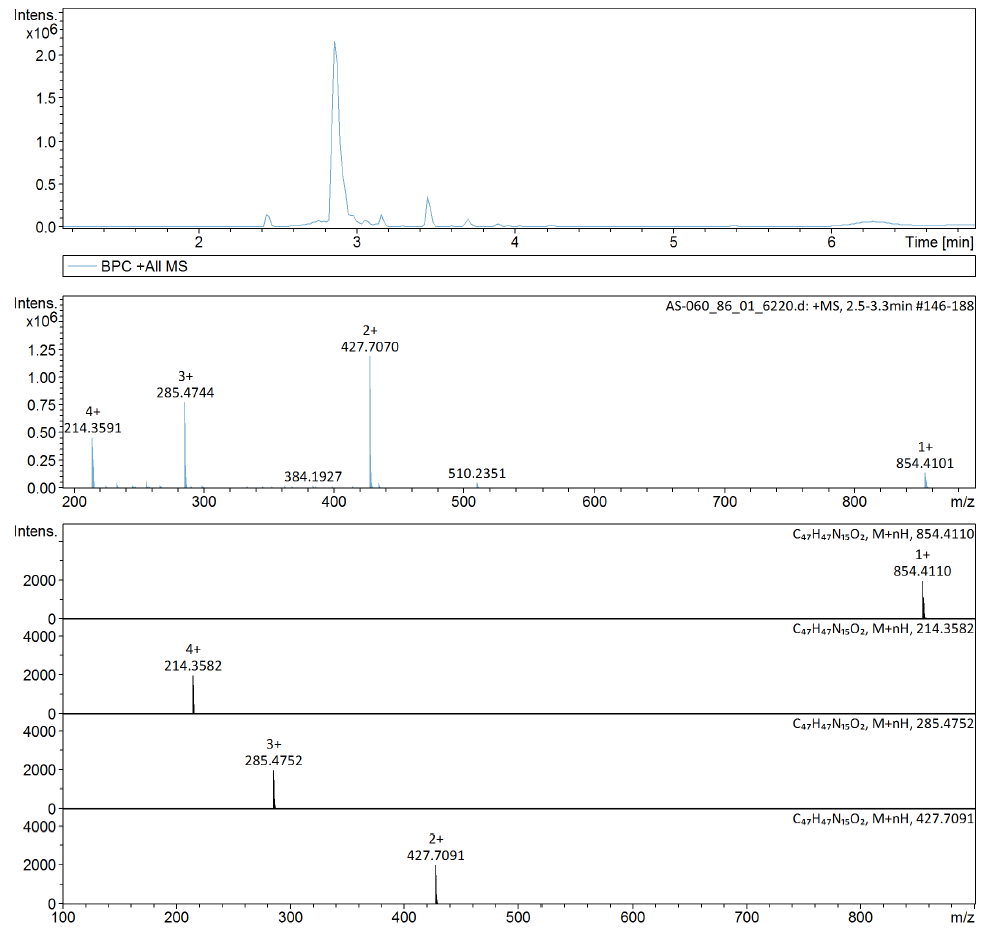

**^1^H NMR spectrum of 2',6-dichloro-1*H*,3'*H*-2,5'-bibenzo[*d*]imidazole (XV – MB_2_Cl)**

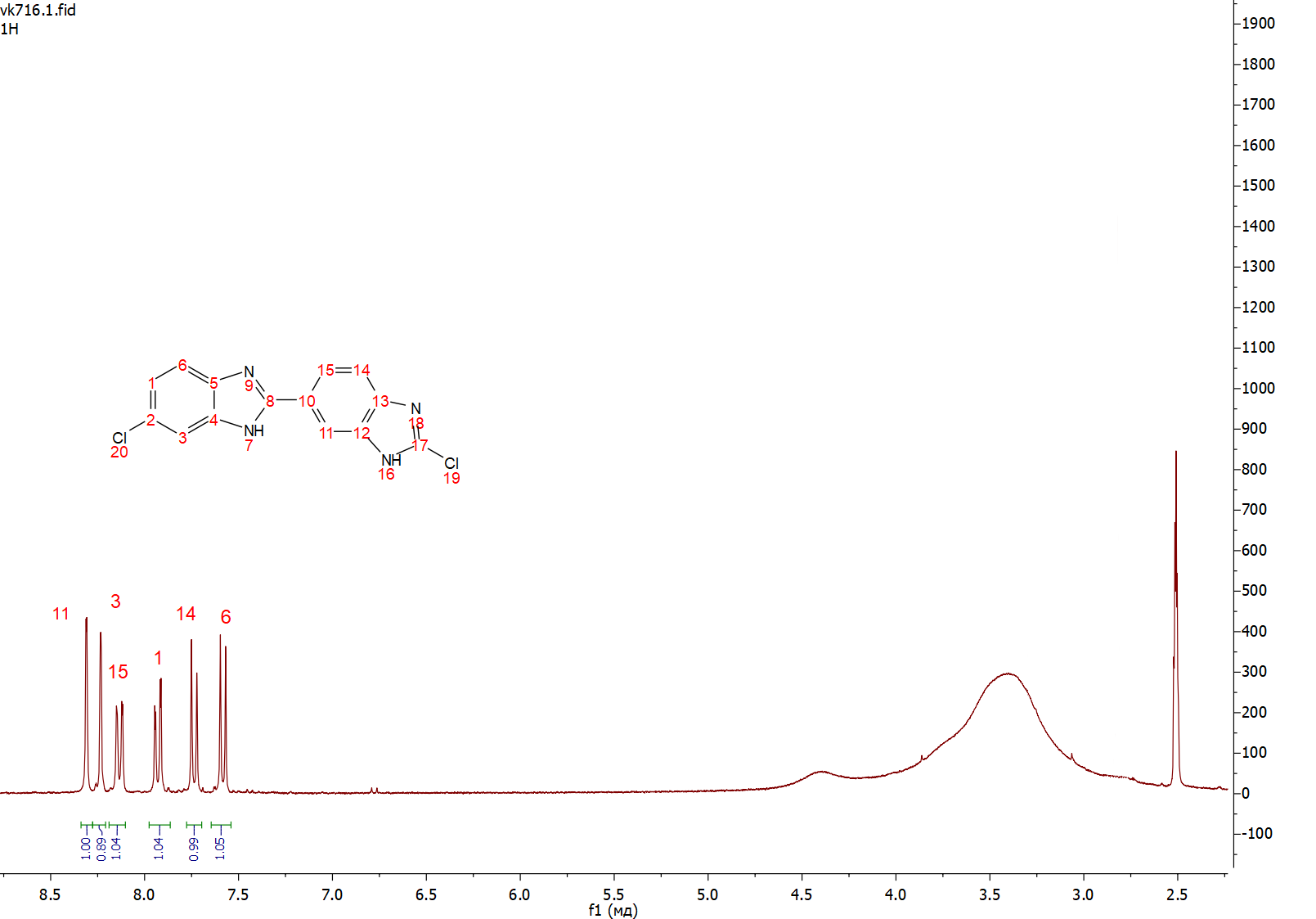

**^13^C NMR spectrum of 2',6-dichloro-1*H*,3'*H*-2,5'-bibenzo[*d*]imidazole (XV – MB_2_Cl)**

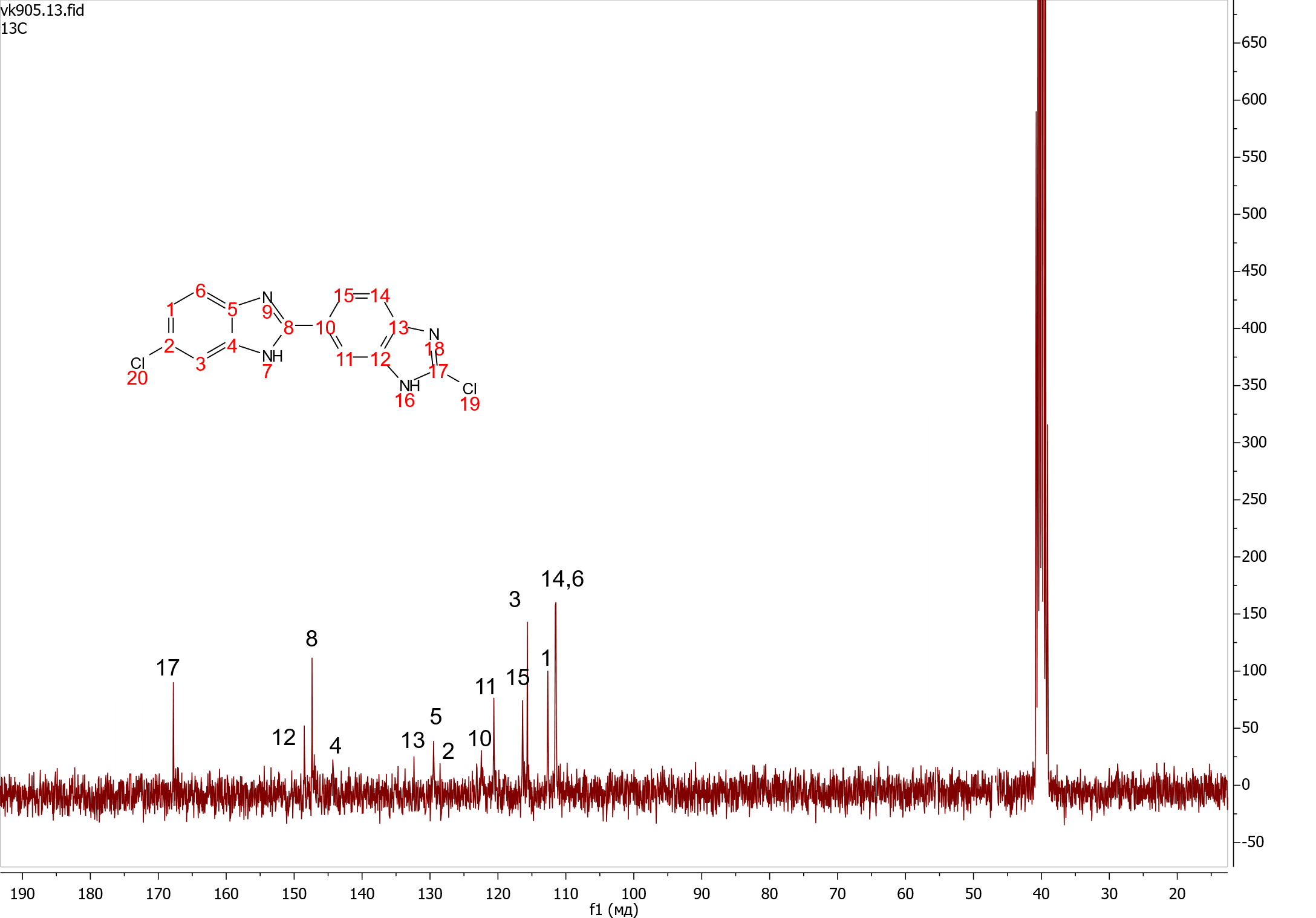

**Fluorescence spectra of *N*,*N*'-(propan-1,3-diyl)bis(2'-(4-methylpiperazin-1-yl)-1*H*,3'*H*-[2,5'-dibenzo[*d*]imidazole]-6-carboxamide) (VIII – RDB_2_(3)) binding to dsDNA**

RDB2(3)

Intensity, rel. units

| Wavelength, nm | 0 | 9 | 22 | 45 | 75 | 110 | 152 | 214 | 294 |
| --- | --- | --- | --- | --- | --- | --- | --- | --- | --- |
|  | 0 | 2 | 5 | 10 | 17 | 25 | 35 | 50 | 70 |
| 350 | 0.560957 | 1.56495 | 2.59119 | 2.42311 | 2.72215 | 2.82273 | 3.2381 | 3.24056 | 3.93212 |
| 351.07 | 0.36036 | 1.11338 | 2.88684 | 4.26309 | 3.59712 | 5.82072 | 4.40088 | 5.02232 | 4.43213 |
| 352 | 0.375375 | 1.53492 | 3.45888 | 5.56586 | 5.69958 | 5.34548 | 6.09176 | 5.81607 | 6.06175 |
| 353.07 | 0.204583 | 2.26543 | 3.17965 | 5.92394 | 5.94553 | 6.39153 | 5.32439 | 5.95871 | 7.28256 |
| 354 | 0.573066 | 3.31772 | 2.83876 | 7.8304 | 7.10945 | 8.1936 | 8.59179 | 7.24346 | 8.71249 |
| 355.07 | 0.583771 | 3.68074 | 5.66687 | 8.47458 | 10.8879 | 9.86842 | 12.0031 | 10.9174 | 10.1781 |
| 356 | 1.1325 | 3.4662 | 5.49867 | 10.5777 | 12.286 | 12.7339 | 13.8651 | 13.4814 | 12.6529 |
| 357.07 | -0.207039 | 3.77572 | 6.67059 | 13.2406 | 14.3411 | 15.9826 | 15.8176 | 16.1136 | 15.4811 |
| 358 | 0.768049 | 5.9581 | 7.59681 | 15.9114 | 18.2532 | 19.4909 | 19.176 | 19.4247 | 19.1681 |
| 359.07 | 0.39604 | 5.10204 | 10.5481 | 21.5185 | 20.8883 | 21.9573 | 23.0396 | 24.4735 | 22.5263 |
| 360 | 0.568074 | 7.11538 | 11.7173 | 22.6753 | 25.0926 | 28.4504 | 28.968 | 28.7495 | 29.2447 |
| 361.06 | 2.12725 | 8.56403 | 14.1403 | 28.9006 | 30.8112 | 32.2646 | 34.7975 | 33.6186 | 34.5703 |
| 361.96 | 1.7452 | 9.22509 | 17.575 | 33.6212 | 37.4662 | 38.0541 | 39.4099 | 39.454 | 40.8205 |
| 363.03 | 1.54589 | 10.8559 | 18.0879 | 38.5808 | 41.6667 | 46.489 | 48.14 | 45.982 | 45.7984 |
| 363.93 | 1.16279 | 12.1236 | 21.7051 | 42.1651 | 48.8565 | 54.4301 | 53.1772 | 52.6115 | 52.715 |
| 365 | 0.403063 | 14.7651 | 25.4187 | 48.7053 | 54.4271 | 60.2238 | 61.9679 | 61.6123 | 61.689 |
| 366.06 | 0 | 16.9523 | 29.0002 | 57.6592 | 61.407 | 68.5703 | 69.6877 | 71.2871 | 71.0575 |
| 366.96 | 1.18812 | 18.8462 | 32.498 | 62.6541 | 70.6198 | 75.3451 | 79.4675 | 78.521 | 77.7912 |
| 368.03 | 0.193237 | 21.2133 | 36.1368 | 71.9234 | 78.1341 | 85.7538 | 86.982 | 87.7736 | 87.2443 |
| 368.93 | 0.970685 | 21.0506 | 38.4158 | 76.6395 | 84.0184 | 90.768 | 94.572 | 95.7082 | 95.1925 |
| 370 | 2.0024 | 25.2024 | 40.1404 | 84.5295 | 92.0351 | 103.594 | 104.091 | 105.374 | 106.229 |
| 371.07 | 1.25654 | 25.8458 | 46.5919 | 93.9953 | 103.176 | 113.613 | 115.49 | 115.9 | 113.942 |
| 372 | 1.88537 | 28.6724 | 48.7495 | 102.916 | 110.337 | 124.52 | 123.926 | 127.142 | 125.683 |
| 373.07 | 2.18427 | 32.1465 | 54.6095 | 111.685 | 118.182 | 133.673 | 136.869 | 135.022 | 133.046 |
| 374 | 2.3175 | 33.4507 | 57.6264 | 116.13 | 129.339 | 141.946 | 141.987 | 144.37 | 144.754 |
| 375.07 | 1.38559 | 35.0808 | 63.4466 | 124.213 | 136.873 | 149.646 | 155.94 | 153.876 | 153.503 |
| 376 | 1.57449 | 38.7692 | 67.5837 | 134.853 | 145.359 | 158.414 | 162.603 | 162.711 | 162.51 |
| 377.07 | 1.6768 | 40.7464 | 70.5782 | 141.771 | 153.494 | 170.664 | 174.543 | 173.736 | 175.069 |
| 378 | 1.17142 | 44.788 | 76.3052 | 149.665 | 162.332 | 179.104 | 181.702 | 186.86 | 182.779 |
| 379.07 | 2.23486 | 46.9156 | 80.9133 | 156.98 | 172.394 | 188.382 | 192.717 | 196.305 | 195.942 |
| 380 | 1.76852 | 48.2701 | 81.0466 | 161.733 | 179.133 | 194.938 | 198.981 | 199.724 | 199.254 |
| 381.06 | 2.33943 | 51.7936 | 85.7602 | 174.389 | 186.37 | 207.423 | 206.697 | 211.906 | 206.756 |
| 381.96 | 2.58193 | 51.068 | 87.7262 | 179.527 | 196.689 | 211.76 | 219.241 | 219.654 | 213.868 |
| 383.03 | 1.68508 | 54.9659 | 93.812 | 187.337 | 202.157 | 223.956 | 227.167 | 227.686 | 226.136 |
| 383.93 | 3.13611 | 57.9536 | 94.5593 | 194.539 | 208.653 | 228.303 | 235.896 | 234.46 | 237.919 |
| 385 | 3.27806 | 57.8416 | 98.8655 | 196.813 | 217.306 | 240.514 | 239.891 | 250.718 | 241.781 |
| 386.06 | 4.0753 | 61.11 | 100.894 | 208.481 | 219.124 | 242.863 | 252.489 | 251.508 | 247.076 |
| 386.96 | 3.59856 | 62.6147 | 105.2 | 211.352 | 229.508 | 255.538 | 257.776 | 259.709 | 252.538 |
| 388.03 | 2.82805 | 63.6487 | 107.853 | 219.447 | 237.801 | 261.769 | 263.452 | 265.444 | 263.466 |
| 388.93 | 3.24118 | 66.8433 | 109.67 | 226.099 | 246.451 | 265.923 | 267.697 | 272.783 | 267.305 |
| 390 | 3.39457 | 67.9325 | 114.64 | 230.523 | 249.796 | 273.538 | 275.64 | 280.921 | 275.262 |
| 391.06 | 3.93623 | 70.5767 | 113.393 | 231.61 | 257.919 | 277.268 | 282.327 | 283.635 | 285.12 |
| 391.96 | 2.49377 | 70.632 | 120.656 | 235.678 | 256.648 | 282.451 | 287.075 | 294.224 | 286.901 |
| 393.03 | 2.51451 | 73.2303 | 120.926 | 246.329 | 262.812 | 286.611 | 286.585 | 291.468 | 287.234 |
| 393.93 | 3.54122 | 72.4019 | 123.038 | 250.191 | 268.061 | 289.157 | 297.988 | 295.775 | 292.844 |
| 395 | 3.5074 | 75.5444 | 121.838 | 250.73 | 272.709 | 296.377 | 296.152 | 298.788 | 297.028 |
| 396.06 | 3.77929 | 75.6506 | 125.051 | 250.685 | 273.477 | 296.304 | 302.632 | 307.016 | 299.692 |
| 396.96 | 2.28137 | 74.3443 | 126.66 | 253.157 | 272.608 | 298.762 | 307.208 | 303.633 | 300.521 |
| 398.03 | 4.31203 | 76.5006 | 124.5 | 258.871 | 276.535 | 300.567 | 307.285 | 308.94 | 306.272 |
| 398.93 | 3.93848 | 76.7068 | 129.766 | 259.931 | 276.977 | 304.314 | 312.151 | 308.466 | 304.785 |
| 400 | 3.53883 | 77.3269 | 129.467 | 259.236 | 282.857 | 304.026 | 309.257 | 310.165 | 306.979 |
| 401.07 | 3.39175 | 79.0244 | 129.768 | 262.128 | 283.111 | 307.296 | 309.281 | 310.141 | 303.622 |
| 402 | 3.04356 | 81.332 | 129.43 | 263.136 | 287.337 | 308.504 | 311.495 | 313.451 | 303.31 |
| 403.07 | 3.42922 | 81.8534 | 132.171 | 265.294 | 282.878 | 309.23 | 308.036 | 309.405 | 307.482 |
| 404 | 3.70447 | 81.3316 | 129.749 | 263.264 | 282.756 | 304.48 | 311.133 | 311.59 | 305.721 |
| 405.07 | 4.48179 | 80.6673 | 128.629 | 265.18 | 279.629 | 306.538 | 307.997 | 311.073 | 303.132 |
| 406 | 5.05807 | 80.0625 | 127.995 | 260.896 | 280.535 | 302.92 | 304.25 | 310.17 | 304.536 |
| 407.07 | 5.04952 | 81.2641 | 129.427 | 255.48 | 277.721 | 300.727 | 306.429 | 311.927 | 303.528 |
| 408 | 4.38262 | 78.5163 | 123.736 | 258.671 | 281.007 | 300.098 | 303.15 | 300.474 | 298.356 |
| 409.07 | 5.14188 | 78.1309 | 126.316 | 257.809 | 274.106 | 299.176 | 304.468 | 297.268 | 299.313 |
| 410 | 3.31319 | 79.5529 | 124.845 | 256.477 | 273.597 | 296.846 | 298.551 | 300.923 | 293.824 |
| 411.06 | 2.42764 | 79.7651 | 125.163 | 250.757 | 273.641 | 289.794 | 295.515 | 303.144 | 288.965 |
| 411.96 | 3.49379 | 79.4919 | 122.725 | 250.936 | 268.327 | 291.735 | 296.223 | 294.829 | 291.192 |
| 413.03 | 6.1574 | 76.8892 | 125.056 | 245.104 | 265.724 | 286.097 | 287.914 | 290.913 | 285.714 |
| 413.93 | 4.57529 | 78.6997 | 120.612 | 244.873 | 265.905 | 287.305 | 284.947 | 287.895 | 283.841 |
| 415 | 4.65567 | 76.0105 | 121.327 | 243.174 | 260.582 | 279.984 | 281.606 | 281.767 | 281.104 |
| 416.06 | 6.16592 | 74.0118 | 119.791 | 239.019 | 254.707 | 275.449 | 276.125 | 280.016 | 274.395 |
| 416.96 | 5.30806 | 74.9632 | 118.182 | 233.14 | 252.137 | 273.181 | 269.906 | 272.492 | 270.088 |
| 418.03 | 6.75248 | 72.4212 | 116.229 | 231.817 | 248.112 | 268.732 | 266.967 | 269.555 | 263.495 |
| 418.93 | 5.38815 | 71.9712 | 112.751 | 226.059 | 248.598 | 260.117 | 262.142 | 268.644 | 260.504 |
| 420 | 5.39707 | 71.7442 | 108.992 | 226.237 | 239.374 | 256.405 | 257.335 | 261.871 | 259.321 |
| 421.06 | 5.49138 | 71.1279 | 111.182 | 217.047 | 238.154 | 251.757 | 251.435 | 254.367 | 253.417 |
| 421.96 | 5.72519 | 67.3016 | 105.68 | 212.358 | 230.112 | 248.117 | 247.575 | 247.959 | 244.303 |
| 423.03 | 5.8117 | 67.9049 | 102.976 | 208.841 | 222.864 | 237.689 | 238.929 | 239.801 | 233.479 |
| 423.93 | 6.31662 | 66.7783 | 102.045 | 203.438 | 215.193 | 235.673 | 238.945 | 238.371 | 232.59 |
| 425 | 6.21247 | 62.0032 | 99.802 | 196.066 | 216.78 | 225.775 | 232.275 | 229.21 | 226.611 |
| 426.06 | 5.32725 | 62.2903 | 96.6443 | 193.555 | 208.148 | 221.327 | 223.509 | 219.769 | 220.387 |
| 426.96 | 5.91949 | 60.3025 | 94.693 | 186.21 | 199.479 | 214.743 | 216.275 | 217.518 | 216.562 |
| 428.03 | 5.8651 | 58.6287 | 91.3726 | 182.12 | 191.471 | 208.53 | 211.901 | 209.526 | 205.308 |
| 428.93 | 6.19628 | 57.2643 | 89.5141 | 178.932 | 189.173 | 198.545 | 205.607 | 199.786 | 200.123 |
| 430 | 4.41968 | 55.9105 | 83.8386 | 169.112 | 184.162 | 197.778 | 195.433 | 196.817 | 191.182 |
| 431.04 | 5.81613 | 54.0431 | 83.1566 | 167.254 | 177.182 | 191.243 | 190.226 | 186.468 | 189.313 |
| 431.94 | 4.14313 | 53.5369 | 81.2568 | 161.258 | 174.575 | 188.117 | 185.133 | 183.753 | 181.152 |
| 432.98 | 4.07609 | 51.7491 | 77.906 | 158.019 | 169.1 | 181.873 | 178.656 | 179.072 | 177.645 |
| 434.02 | 4.53291 | 47.452 | 77.2699 | 148.141 | 159.887 | 171.981 | 172.089 | 167.331 | 168.975 |
| 435.07 | 5.33281 | 47.6384 | 74.3058 | 148.189 | 159.438 | 166.328 | 166.908 | 169.711 | 163.325 |
| 435.97 | 6.35961 | 48.2315 | 72.0313 | 140.222 | 154.8 | 160.699 | 162.262 | 160.986 | 158.193 |
| 437.01 | 4.89428 | 46.5436 | 68.9178 | 135.189 | 144.858 | 155.474 | 155.659 | 156.605 | 154.998 |
| 438.05 | 4.66019 | 43.4349 | 69.946 | 135.394 | 141.315 | 148.841 | 151.727 | 151.147 | 147.745 |
| 438.95 | 5.14192 | 43.2565 | 65.6818 | 127.723 | 137.565 | 146.675 | 147.983 | 146.884 | 143.536 |
| 440 | 5.43584 | 42.2846 | 63.2172 | 124.439 | 134.322 | 138.465 | 139.733 | 138.602 | 139.653 |
| 441.06 | 4.98107 | 40.3696 | 62.2425 | 118.568 | 128.462 | 135.682 | 136.344 | 132.853 | 135.399 |
| 441.96 | 4.84211 | 38.5408 | 58.7641 | 114.954 | 125.315 | 133.062 | 132.996 | 130.725 | 127.119 |
| 443.03 | 5.21739 | 38.4927 | 59.6162 | 112.016 | 121.244 | 129.841 | 128.552 | 128.169 | 126.524 |
| 443.93 | 4.99904 | 38.4314 | 57.2927 | 109.216 | 117.525 | 120.675 | 122.01 | 126.109 | 120.157 |
| 445 | 5.63948 | 35.8184 | 54.2194 | 101.238 | 112.216 | 118.221 | 118.743 | 118.137 | 112.515 |
| 446.06 | 5.63764 | 36.1928 | 55.4287 | 100.422 | 107.459 | 113.769 | 110.901 | 113.255 | 112.575 |
| 446.96 | 3.38174 | 34.4899 | 51.2552 | 98.2234 | 103.183 | 109.057 | 111.598 | 111.473 | 107.868 |
| 448.03 | 4.99168 | 33.1192 | 49.2809 | 93.8628 | 99.3115 | 107.711 | 106.434 | 107.187 | 105.397 |
| 448.93 | 4.52781 | 32.413 | 47.8469 | 92.6609 | 100.486 | 103.655 | 103.525 | 103.629 | 101.788 |
| 450 | 4.20084 | 32.3944 | 46.798 | 92.1404 | 96.7866 | 100.908 | 100.67 | 101.639 | 97.5559 |
| 451.04 | 4.78755 | 29.7281 | 46.759 | 89.36 | 90.5398 | 98.0312 | 97.7659 | 97.503 | 97.2164 |
| 451.94 | 6.1127 | 30.0403 | 45.0136 | 86.4833 | 91.293 | 96.0725 | 96.4563 | 95.0813 | 90.6 |
| 452.98 | 4.62428 | 29.5082 | 40.8951 | 79.1889 | 88.0883 | 92.2563 | 90.2889 | 90.9464 | 89.2193 |
| 454.02 | 4.22103 | 28.084 | 41.1909 | 80.0511 | 84.6123 | 88.5898 | 88.504 | 89.4359 | 86.9043 |
| 455.07 | 4.85814 | 27.3277 | 40.8756 | 78.3505 | 83.111 | 85.6536 | 84.802 | 84.1806 | 85.1553 |
| 455.97 | 5.62888 | 27.6035 | 40.0319 | 75.6027 | 81.6199 | 83.3158 | 83.6086 | 81.1613 | 80.9524 |
| 457.01 | 4.32982 | 27.0991 | 38.0071 | 71.2998 | 76.7713 | 78.3047 | 79.603 | 79.9527 | 78.3965 |
| 458.05 | 5.25394 | 25.9377 | 36.4039 | 69.1524 | 73.8848 | 78.9414 | 78.5867 | 77.957 | 78.1918 |
| 458.95 | 4.18243 | 24.9951 | 35.7456 | 67.0758 | 71.6024 | 74.2845 | 74.4471 | 74.2212 | 74.6237 |
| 460 | 3.03203 | 23.6416 | 35.7143 | 64.822 | 70.0826 | 70.7537 | 69.8611 | 70.2341 | 70.7346 |
| 461.06 | 4.93733 | 23.6349 | 32.8947 | 60.6575 | 66.6667 | 70.4012 | 70.8019 | 68.7611 | 69.9372 |
| 461.96 | 4.35956 | 23.3885 | 32.5219 | 62.8781 | 62.8029 | 67.7475 | 67.8973 | 67.4319 | 64.048 |
| 463.03 | 3.98331 | 22.4333 | 30.9538 | 56.3199 | 61.9645 | 64.3544 | 64.6312 | 65.9898 | 61.266 |
| 463.93 | 3.92805 | 22.6474 | 31.2051 | 55.6138 | 60.8013 | 63.6807 | 61.9208 | 62.3608 | 61.9414 |
| 465 | 4.84215 | 21.0861 | 30.8872 | 54.3501 | 56.6735 | 60.0166 | 61.7608 | 58.7393 | 58.9028 |
| 466.06 | 6.34982 | 20.529 | 27.1863 | 52.9248 | 56.7478 | 58.6713 | 55.9544 | 56.9249 | 55.8308 |
| 466.96 | 4.90092 | 19.2457 | 26.8904 | 49.5733 | 54.7394 | 57.4869 | 54.7497 | 56.6184 | 56.0556 |
| 468.03 | 3.79696 | 19.9661 | 27.5774 | 49.726 | 50.5729 | 55.1948 | 53.6094 | 53.1401 | 53.5538 |
| 468.93 | 3.86432 | 18.178 | 26.1866 | 46.042 | 49.3827 | 51.4512 | 50.9704 | 50.6256 | 50.4285 |
| 470 | 5.03018 | 17.7109 | 24.5885 | 46.287 | 47.0635 | 48.8056 | 48.3384 | 49.0697 | 49.3135 |
| 471.04 | 4.57095 | 15.9751 | 23.8963 | 44.0582 | 46.1028 | 46.0554 | 49.2161 | 47.8844 | 45.9163 |
| 471.94 | 3.97803 | 17.0112 | 22.6562 | 43.3616 | 44.0493 | 45.5097 | 46.9839 | 48.6663 | 44.5822 |
| 472.98 | 3.4116 | 15.9585 | 21.38 | 39.8849 | 43.9382 | 44.8769 | 44.3473 | 43.654 | 42.1521 |
| 474.02 | 3.9801 | 16.2055 | 21.3096 | 38.3102 | 41.3512 | 42.2906 | 42.3259 | 41.1823 | 42.0479 |
| 475.07 | 4.329 | 15.9784 | 21.0192 | 36.8555 | 39.3095 | 38.7725 | 39.2278 | 41.7638 | 40.6151 |
| 475.97 | 3.3626 | 13.5729 | 19.1726 | 34.7149 | 38.4062 | 38.4538 | 37.5624 | 39.0753 | 37.1595 |
| 477.01 | 3.21092 | 12.8902 | 20.5671 | 32.6563 | 35.4754 | 36.7432 | 38.2121 | 37.4975 | 37.6075 |
| 478.05 | 3.48646 | 14.3087 | 18.9554 | 31.7659 | 36.1667 | 35.8789 | 36.4372 | 34.8762 | 33.6067 |
| 478.95 | 3.57143 | 11.267 | 18.5374 | 33.0379 | 34.6672 | 33.6251 | 34.4064 | 33.7193 | 33.9844 |
| 480 | 3.98963 | 10.9504 | 18.341 | 29.5119 | 32.5156 | 34.0862 | 31.4895 | 33.6217 | 31.3385 |
| 481.04 | 3.70851 | 11.3919 | 18.4458 | 29.4799 | 30.8927 | 30.5421 | 31.8313 | 33.1263 | 32.3048 |
| 481.94 | 4.2245 | 10.2342 | 15.4146 | 28.3165 | 28.344 | 31.4292 | 28.8714 | 31.023 | 31.3099 |
| 482.98 | 3.03092 | 12.5712 | 15.3082 | 26.9424 | 27.3339 | 29.1837 | 28.341 | 27.8229 | 27.8996 |
| 484.02 | 4.04367 | 9.96873 | 15.6773 | 24.2954 | 28.5889 | 28.3368 | 27.1903 | 26.4571 | 25.8207 |
| 485.07 | 3.47435 | 10.5459 | 14.8431 | 24.808 | 26.2253 | 26.9714 | 28.2051 | 25.1674 | 25.8241 |
| 485.97 | 3.12695 | 10.7505 | 13.6519 | 24.1698 | 26.6123 | 24.9221 | 26.4447 | 24.5059 | 24.4379 |
| 487.01 | 2.4948 | 8.27586 | 14.9724 | 23.5194 | 24.5919 | 24.4473 | 23.7727 | 24.1497 | 23.6393 |
| 488.05 | 3.09777 | 9.75513 | 12.5235 | 21.2553 | 22.0531 | 24.1084 | 24.2972 | 24.0405 | 21.2089 |
| 488.95 | 2.03874 | 8.01644 | 10.8805 | 21.0118 | 21.335 | 22.9654 | 20.9847 | 22.655 | 21.9667 |
| 490 | 3.17902 | 8.9123 | 11.7421 | 20.3707 | 20.5021 | 21.0275 | 20.4446 | 21.8075 | 22.1545 |
| 491.04 | 3.02358 | 8 | 10.9322 | 21.0276 | 18.5374 | 20.2177 | 21.0093 | 20.9668 | 20.3025 |
| 491.94 | 2.90879 | 8.22079 | 9.91061 | 17.9267 | 20.3116 | 18.064 | 20.3035 | 19.2694 | 19.6565 |
| 492.98 | 2.9458 | 6.99732 | 12.1399 | 17.2621 | 19.2539 | 17.8288 | 18.7587 | 19.5552 | 19.0856 |
| 494.02 | 3.861 | 8.85126 | 9.96483 | 15.3628 | 19.163 | 20.4042 | 17.276 | 18.8178 | 19.5476 |
| 495.07 | 2.01005 | 7.72986 | 10.3134 | 15.3846 | 17.5843 | 17.2306 | 18.5484 | 17.5773 | 17.2975 |
| 495.97 | 3.27582 | 6.34128 | 10.3563 | 15.4872 | 14.9373 | 16.7142 | 16.7356 | 17.8859 | 15.366 |
| 497.01 | 2.07598 | 5.50459 | 9.98668 | 16.5452 | 16.2184 | 16.3065 | 15.7938 | 16.0739 | 16.4559 |
| 498.05 | 2.42131 | 8.32149 | 8.21479 | 14.697 | 15.6417 | 15.4377 | 14.9555 | 16.8759 | 14.9798 |
| 498.95 | 2.86592 | 7.80608 | 9.25034 | 16.0741 | 16.2866 | 15.5211 | 14.5755 | 14.1575 | 13.0895 |
| 500 | 2.17649 | 6.70755 | 8.29456 | 12.955 | 14.0472 | 15.74 | 13.7958 | 13.7517 | 14.9988 |
| 501.04 | 1.76367 | 5.79305 | 7.38007 | 13.3279 | 11.7139 | 12.955 | 14.6303 | 14.8683 | 12.6743 |
| 501.94 | 4.10317 | 5.31371 | 6.55611 | 13.4267 | 13.7624 | 13.2989 | 12.1611 | 13.2424 | 13.1978 |
| 502.98 | 2.54723 | 5.72557 | 6.87702 | 12.987 | 12.2407 | 12.878 | 12.5423 | 11.3469 | 13.1999 |
| 504.02 | 2.42082 | 4.73747 | 6.88538 | 10.2963 | 11.8901 | 12.9674 | 10.93 | 11.5061 | 11.5212 |
| 505.07 | 2.18949 | 5.72301 | 5.95116 | 10.6447 | 12.1951 | 12.5273 | 11.5037 | 10.0369 | 11.1673 |
| 505.97 | 2.75591 | 5.29601 | 4.7547 | 11.2862 | 11.2196 | 11.6959 | 11.4847 | 10.8499 | 9.33984 |
| 507.01 | 1.80796 | 5.73463 | 7.11595 | 10.0096 | 10.252 | 12.6216 | 10.0355 | 10.8268 | 9.44534 |
| 508.05 | 2.41692 | 4.79478 | 5.68311 | 7.91766 | 9.92282 | 10.7811 | 9.52744 | 11.7695 | 10.4634 |
| 508.95 | 3.62551 | 4.04367 | 5.95238 | 9.11736 | 8.88797 | 10.1392 | 8.42408 | 9.95552 | 9.32063 |
| 510 | 3.45556 | 5.59217 | 6.55542 | 8.76024 | 8.34465 | 9.81162 | 9.76928 | 9.94431 | 7.7951 |
| 511.04 | 2.10128 | 4.47974 | 5.92467 | 8.75 | 8.05477 | 8.61362 | 9.27388 | 8.84397 | 9.61969 |
| 511.94 | 2.40336 | 4.37724 | 4.64037 | 8.03922 | 8.59775 | 7.98036 | 8.41544 | 7.57576 | 8.94806 |
| 512.98 | 1.67926 | 2.80056 | 5.45613 | 8.13796 | 6.79378 | 6.59868 | 9.03862 | 9.05979 | 7.67367 |
| 514.02 | 2.52771 | 4.70636 | 4.70991 | 7.00253 | 7.71056 | 6.90496 | 7.20265 | 7.74732 | 7.89422 |
| 515.07 | 0.999201 | 3.24741 | 4.16254 | 7.3471 | 8.1483 | 7.34441 | 7.76247 | 7.8399 | 6.39659 |
| 515.97 | 2.29214 | 3.82263 | 5.10308 | 6.91017 | 7.56461 | 6.6 | 7.1885 | 7.75031 | 7.22175 |
| 517.01 | 1.73745 | 4.40107 | 4.86567 | 6.05693 | 6.55738 | 6.9472 | 6.52174 | 8.08123 | 6.50054 |
| 518.05 | 1.17925 | 3.32096 | 4.48814 | 7.32238 | 5.37968 | 6.89655 | 5.41951 | 8.03958 | 6.96864 |
| 518.95 | 1.42857 | 3.45556 | 3.53283 | 5.56472 | 7.49625 | 6.78931 | 5.43478 | 4.68241 | 6.42398 |
| 520 | 2.20397 | 2.05915 | 4.65426 | 6.15973 | 6.07507 | 5.48804 | 6.82947 | 4.95903 | 5.7717 |
| 521.02 | 2.20088 | 4.18911 | 3.14607 | 6.34726 | 5.87401 | 6.26346 | 5.46891 | 6.32426 | 5.93472 |
| 522.05 | 0.582524 | 4.13142 | 4.75698 | 6.14105 | 6.16302 | 5.8117 | 5.40645 | 4.41014 | 5.34442 |
| 522.94 | 1.37336 | 2.80794 | 4.04776 | 5.73504 | 5.07507 | 4.18827 | 6.01942 | 3.57293 | 4.70773 |
| 523.97 | 2.14718 | 3.67505 | 3.73709 | 4.61095 | 4.46338 | 5.52159 | 6.00663 | 5.30721 | 5.52147 |
| 525 | 1.40478 | 3.84986 | 2.77541 | 4.10718 | 3.51312 | 5.17928 | 3.61736 | 5.55784 | 5.22749 |
| 526.02 | 1.96117 | 3.38443 | 3.75548 | 4.82253 | 4.87391 | 4.75796 | 6.13748 | 5.2417 | 3.3966 |
| 527.05 | 2.65849 | 3.4338 | 3.14536 | 3.58499 | 5.33881 | 4.95427 | 3.71021 | 3.15989 | 4.19497 |
| 527.94 | 2.07078 | 1.32125 | 2.84339 | 2.8186 | 3.82448 | 3.924 | 4.61847 | 4.40723 | 5.16055 |
| 528.97 | 1.79211 | 2.56158 | 3.3051 | 3.73134 | 3.27511 | 4.48256 | 2.96033 | 3.32005 | 3.74532 |
| 530 | 1.02229 | 2.32243 | 3.01447 | 3.39847 | 3.14005 | 4.82606 | 5.35861 | 4.74121 | 2.85016 |
| 531.04 | 0.202757 | 3.19549 | 4.62088 | 3.31126 | 5.16209 | 3.61836 | 4.40705 | 3.92562 | 3.69506 |
| 531.94 | 0.779271 | 2.41009 | 3.2888 | 4.73118 | 4.55166 | 3.89788 | 2.89196 | 3.04149 | 3.67789 |
| 532.98 | 1.88009 | 3.58494 | 1.62272 | 3.14005 | 3.59569 | 3.37771 | 3.54684 | 3.39487 | 3.62538 |
| 534.02 | 1.77655 | 3.23748 | 1.72302 | 2.55553 | 3.19808 | 3.70294 | 2.82372 | 4.02542 | 3.67347 |
| 535.07 | 1.92827 | 1.22001 | 1.9202 | 4.73738 | 1.91775 | 1.89394 | 3.69565 | 2.56904 | 2.40385 |
| 535.97 | 0.774293 | 1.83824 | 2.96804 | 4.4999 | 1.77096 | 4.2203 | 3.16581 | 3.62319 | 4.29017 |
| 537.01 | 0.798244 | 1.34615 | 3.19013 | 3.22928 | 3.66654 | 3.25203 | 2.43309 | 2.1182 | 2.44549 |
| 538.05 | 0.190006 | 2.32423 | 3.003 | 0.60988 | 2.92455 | 2.41838 | 3.63481 | 0.62448 | 2.19912 |
| 538.95 | 1.24224 | 2.9458 | 2.37171 | 2.82885 | 3.77433 | 2.7191 | 3.47862 | 3.21474 | 3.53288 |
| 540 | 1.75884 | 2.31839 | 1.60353 | 2.66612 | 3.22971 | 2.91262 | 3.57218 | 2.87179 | 2.54777 |
| 541.02 | 0.778362 | 1.60933 | 1.25 | 2.57898 | 3.35902 | 4.28397 | 2.41158 | 2.39664 | 1.38532 |
| 542.05 | 0.987752 | 1.53644 | 0.619067 | 3.0248 | 3.14713 | 2.55503 | 2.21953 | 1.8649 | 1.72944 |
| 542.94 | 0.39557 | 2.56259 | 2.92626 | 3.43938 | 1.47121 | 2.2805 | 3.05188 | 2.46858 | 2.22853 |
| 543.97 | 0.960246 | 1.42566 | 2.51256 | 1.95122 | 2.36127 | 2.61833 | 2.43408 | 1.22449 | 1.51025 |
| 545 | 1.20506 | 2.04248 | 1.55945 | 2.53165 | 1.726 | 2.35748 | 3.09789 | 1.19832 | 1.96464 |
| 546.02 | 1.16347 | 1.35896 | 2.03758 | 1.39137 | 2.09604 | 2.29024 | 2.45499 | 2.86709 | 0.814166 |
| 547.05 | 1.73177 | 2.61044 | 2.60992 | 2.45329 | 2.886 | 2.11987 | 1.20797 | 2.69318 | 1.57884 |
| 547.94 | 0.201126 | 1.64541 | 1.80099 | 1.59236 | 2.33873 | 2.03583 | 1.69312 | 2.27964 | 2.33645 |

**Fluorescence spectra of *N*,*N*'-(heptane-1,7-diyl)bis(2'-(4-methylpiperazin-1-yl)-1*H*,3'*H*-[2,5'-dibenzo[*d*]imidazole]-6-carboxamide) (IX – RDB_2_(7)) binding to dsDNA**

RDB2(7)

Intensity, rel. units

| Wavelength, nm | 0 |
| --- | --- |
| 350 | -0.39557 |
| 351.07 | -0.413223 |
| 352 | -0.581959 |
| 353.07 | 0.588697 |
| 354 | 0.781097 |
| 355.07 | 0.449539 |
| 356 | 0.231054 |
| 357.07 | -0.738825 |
| 358 | 1.10327 |
| 359.07 | 0.882029 |
| 360 | 0.798882 |
| 361.06 | 1.01895 |
| 361.96 | -0.593354 |
| 363.03 | 0.205804 |
| 363.93 | 0.588005 |
| 365 | 0.213721 |
| 366.06 | 0 |
| 366.96 | 0.83455 |
| 368.03 | 0.207555 |
| 368.93 | 1.45349 |
| 370 | 1.83262 |
| 371.07 | 0 |
| 372 | 0.209776 |
| 373.07 | 0.599042 |
| 374 | 0.61627 |
| 375.07 | -0.205888 |
| 376 | -0.81367 |
| 377.07 | 1.84729 |
| 378 | 1.65494 |
| 379.07 | 1.76022 |
| 380 | 0 |
| 381.06 | 0.406421 |
| 381.96 | 1.27065 |
| 383.03 | 0 |
| 383.93 | 2.52154 |
| 385 | 1.07596 |
| 386.06 | 1.68173 |
| 386.96 | 1.72117 |
| 388.03 | 0.609013 |
| 388.93 | 2.3678 |
| 390 | 3.13087 |
| 391.06 | 3.21268 |
| 391.96 | 1.94091 |
| 393.03 | 2.56082 |
| 393.93 | 1.86451 |
| 395 | 3.36834 |
| 396.06 | 1.82778 |
| 396.96 | 3.04216 |
| 398.03 | 2.85598 |
| 398.93 | 2.76596 |
| 400 | 2.27179 |
| 401.07 | 2.69934 |
| 402 | 4.16667 |
| 403.07 | 3.82194 |
| 404 | 3.96246 |
| 405.07 | 4.13394 |
| 406 | 4.3535 |
| 407.07 | 4.458 |
| 408 | 3.74532 |
| 409.07 | 3.48432 |
| 410 | 4.31965 |
| 411.06 | 4.72407 |
| 411.96 | 5.15243 |
| 413.03 | 5.48871 |
| 413.93 | 5.74485 |
| 415 | 5.08958 |
| 416.06 | 4.53466 |
| 416.96 | 5.23455 |
| 418.03 | 5.77 |
| 418.93 | 6.11995 |
| 420 | 7.2329 |
| 421.06 | 5.31915 |
| 421.96 | 5.68679 |
| 423.03 | 6.19923 |
| 423.93 | 5.87248 |
| 425 | 5.50423 |
| 426.06 | 4.26309 |
| 426.96 | 4.53515 |
| 428.03 | 8.09544 |
| 428.93 | 7.55124 |
| 430 | 5.68962 |
| 431.04 | 5.32649 |
| 431.94 | 6.02911 |
| 432.98 | 5.56626 |
| 434.02 | 7.26443 |
| 435.07 | 4.8119 |
| 435.97 | 6.91534 |
| 437.01 | 6.1262 |
| 438.05 | 5.47089 |
| 438.95 | 5.36481 |
| 440 | 4.74496 |
| 441.06 | 6.07533 |
| 441.96 | 4.36015 |
| 443.03 | 5.25677 |
| 443.93 | 3.92238 |
| 445 | 5.17799 |
| 446.06 | 3.68249 |
| 446.96 | 5.50964 |
| 448.03 | 5.24511 |
| 448.93 | 4.89696 |
| 450 | 4.69484 |
| 451.04 | 5.19818 |
| 451.94 | 5.5396 |
| 452.98 | 4.59082 |
| 454.02 | 5.42345 |
| 455.07 | 5.82726 |
| 455.97 | 6.30209 |
| 457.01 | 5.94413 |
| 458.05 | 4.37429 |
| 458.95 | 5.04414 |
| 460 | 7.20865 |
| 461.06 | 4.9613 |
| 461.96 | 4.63499 |
| 463.03 | 6.11685 |
| 463.93 | 5.09704 |
| 465 | 5.58659 |
| 466.06 | 4.95049 |
| 466.96 | 5.58548 |
| 468.03 | 5.01605 |
| 468.93 | 5.77229 |
| 470 | 4.50704 |
| 471.04 | 5.34861 |
| 471.94 | 5.46117 |
| 472.98 | 4.09165 |
| 474.02 | 4.69283 |
| 475.07 | 4.5972 |
| 475.97 | 4.49486 |
| 477.01 | 5.75185 |
| 478.05 | 3.5693 |
| 478.95 | 4.09277 |
| 480 | 3.36434 |
| 481.04 | 3.83065 |
| 481.94 | 4.61014 |
| 482.98 | 3.98565 |
| 484.02 | 2.85831 |
| 485.07 | 3.22191 |
| 485.97 | 4.20252 |
| 487.01 | 4.19832 |
| 488.05 | 5.53541 |
| 488.95 | 4.32986 |
| 490 | 3.72825 |
| 491.04 | 2.44349 |
| 491.94 | 3.7461 |
| 492.98 | 4.24328 |
| 494.02 | 3.64151 |
| 495.07 | 3.7461 |
| 495.97 | 4.45344 |
| 497.01 | 3.58877 |
| 498.05 | 3.37771 |
| 498.95 | 2.13995 |
| 500 | 2.39952 |
| 501.04 | 4.09516 |
| 501.94 | 3.15395 |
| 502.98 | 3.03767 |
| 504.02 | 2.40288 |
| 505.07 | 4.5977 |
| 505.97 | 3.21285 |
| 507.01 | 1.42624 |
| 508.05 | 3.04569 |
| 508.95 | 3.82525 |
| 510 | 3.27735 |
| 511.04 | 2.17817 |
| 511.94 | 1.76991 |
| 512.98 | 3.56436 |
| 514.02 | 2.61359 |
| 515.07 | 1.146 |
| 515.97 | 2.72194 |
| 517.01 | 3.14928 |
| 518.05 | 3.42622 |
| 518.95 | 1.44152 |
| 520 | 2.93317 |
| 521.02 | 1.82334 |
| 522.05 | 2.52932 |
| 522.94 | 3.44012 |
| 523.97 | 2.80838 |
| 525 | 1.92226 |
| 526.02 | 2.20884 |
| 527.05 | 1.42945 |
| 527.94 | 4.38789 |
| 528.97 | 1.01709 |
| 530 | 2.55591 |
| 531.04 | 1.42074 |
| 531.94 | 1.45955 |
| 532.98 | 2.3571 |
| 534.02 | 1.62569 |
| 535.07 | 3.35079 |
| 535.97 | 1.43296 |
| 537.01 | 2.43457 |
| 538.05 | 2.06228 |
| 538.95 | 0.19988 |
| 540 | 2.45449 |
| 541.02 | 0.821524 |
| 542.05 | 1.49573 |
| 542.94 | 1.77866 |
| 543.97 | 2.82302 |
| 545 | 1.22825 |
| 546.02 | 1.24069 |
| 547.05 | 0.395648 |
| 547.94 | 1.24922 |
| 548.97 | 2.01288 |

| Wavelength, nm | 9 | 22 | 45 | 75 | 110 | 152 | 214 | 294 | 372 | 446 |
| --- | --- | --- | --- | --- | --- | --- | --- | --- | --- | --- |
|  | 2 | 5 | 10 | 17 | 25 | 35 | 50 | 70 | 90 | 110 |
| 350 | 1.48231 | 1.14613 | 0 | 0 | 0.191865 | 1.47275 | 1.69078 | -0.381025 | 0.836295 | 0.585709 |
| 351.07 | 1.42974 | 0.569801 | 1.14329 | 0.978474 | 1.02648 | 1.48841 | 0.190114 | 1.12634 | 2.51645 | 2.14362 |
| 352 | 0.570776 | 1.13744 | 0.815494 | 0.953107 | 0.963577 | 0.815827 | 1.12973 | 1.52555 | 2.19338 | 1.99023 |
| 353.07 | 0.761905 | 1.12803 | 0.977135 | 1.36081 | 1.94062 | 1.91131 | 2.81057 | 1.64069 | 3.11412 | 1.46172 |
| 354 | 0.588697 | 1.49198 | 1.35633 | 1.93761 | 1.33766 | 0 | 1.59872 | 2.78862 | 2.79553 | 2.60271 |
| 355.07 | 0.780336 | 2.32423 | 2.17391 | 2.01898 | 1.2163 | 2.19619 | 3.22703 | 1.8615 | 2.49568 | 3.07025 |
| 356 | 1.39804 | 0.37679 | 3.16393 | 2.54105 | 3.64404 | 2.10446 | 4.5854 | 3.48028 | 3.97126 | 3.56013 |
| 357.07 | 1.11462 | 2.56551 | 3.84763 | 3.85356 | 4.50098 | 3.96904 | 3.95406 | 5.88124 | 4.60829 | 5.00481 |
| 358 | 2.48709 | 3.4999 | 3.18091 | 3.11284 | 4.65904 | 3.41168 | 5.01505 | 4.42138 | 6.27075 | 7.43562 |
| 359.07 | 2.27402 | 3.85654 | 4.13223 | 5.87002 | 5.97072 | 5.45596 | 7.46269 | 6.5981 | 6.16362 | 8.40673 |
| 360 | 2.2164 | 2.7528 | 4.13612 | 4.79712 | 5.32755 | 8.25959 | 6.19315 | 7.39617 | 8.16803 | 9.18635 |
| 361.06 | 3.04009 | 4.02222 | 5.03681 | 6.55738 | 6.9958 | 7.77546 | 9.1391 | 11.245 | 10.6195 | 10.5164 |
| 361.96 | 4.64127 | 5.103 | 7.84929 | 8.09637 | 9.24321 | 9.92745 | 10.7832 | 12.4854 | 11.8688 | 12.0458 |
| 363.03 | 5.57157 | 5.75459 | 7.98882 | 8.6711 | 10.6635 | 12.4166 | 12.5564 | 13.4119 | 12.9548 | 14.3667 |
| 363.93 | 4.97703 | 7.74743 | 7.22092 | 11.252 | 12.6909 | 11.8871 | 15.242 | 15.3815 | 17.3863 | 15.2049 |
| 365 | 5.23877 | 7.49136 | 9.71066 | 12.5573 | 14.689 | 16.3529 | 17.0588 | 17.7492 | 18.3639 | 20.2041 |
| 366.06 | 6.82128 | 8.86561 | 11.4899 | 14.2456 | 15.7842 | 16.5531 | 19.308 | 19.5204 | 21.4149 | 23.7762 |
| 366.96 | 7.54717 | 10.2574 | 15.3601 | 16.172 | 17.1429 | 19.0678 | 22.1531 | 22.5386 | 23.9439 | 24.5731 |
| 368.03 | 7.81097 | 10.5993 | 15.5131 | 17.6399 | 18.9465 | 21.5922 | 23.3787 | 25.6309 | 26.8901 | 28.3924 |
| 368.93 | 10.7653 | 13.5528 | 17.5264 | 18.4825 | 20.5954 | 24.5869 | 26.0686 | 29.3663 | 29.9504 | 32.0093 |
| 370 | 10.4778 | 13.8321 | 17.7169 | 22.168 | 24.1763 | 25.9349 | 27.5005 | 32.4779 | 32.4769 | 34.1595 |
| 371.07 | 11.5082 | 15.6154 | 19.9849 | 24.18 | 28.3417 | 30.051 | 31.7398 | 34.6116 | 37.9942 | 38.1347 |
| 372 | 13.4069 | 17.602 | 23.3713 | 25.291 | 28.9977 | 31.5913 | 35.6219 | 37.1642 | 39.7826 | 42.5105 |
| 373.07 | 12.8451 | 18.74 | 26.5506 | 27.3414 | 32.3302 | 35.5225 | 38.3647 | 42 | 43.12 | 44.982 |
| 374 | 15.0602 | 21.0227 | 26.9084 | 30.1733 | 34.6453 | 35.6936 | 41.6503 | 43.3564 | 47.361 | 47.4154 |
| 375.07 | 15.0723 | 23.4493 | 28.469 | 32.4739 | 38.0521 | 39.4388 | 44.1729 | 46.3898 | 49.3151 | 50.2771 |
| 376 | 15.5298 | 21.9671 | 29.0685 | 30.9365 | 39.4684 | 42.4028 | 45.6311 | 51.1773 | 52.3498 | 55.6575 |
| 377.07 | 17.9828 | 25.7222 | 32.7411 | 36.1165 | 39.2385 | 47.4697 | 50.8411 | 54.8523 | 56.5101 | 60.3212 |
| 378 | 17.6064 | 24.4236 | 32.6252 | 38.7597 | 42.8764 | 46.8494 | 52.4873 | 59.0333 | 60.3766 | 61.1252 |
| 379.07 | 19.7707 | 27.5454 | 36.2033 | 39.5525 | 45.1862 | 50.164 | 56.999 | 59.3907 | 61.2929 | 64.9701 |
| 380 | 19.4499 | 29.0788 | 38.4326 | 40.4185 | 47.4009 | 53.5044 | 60.1731 | 61.3245 | 65.3961 | 69.377 |
| 381.06 | 22.2351 | 30.808 | 38.4986 | 44.8819 | 53.6926 | 53.7044 | 60.1533 | 63.9329 | 67.2442 | 70.7753 |
| 381.96 | 21.38 | 32.8975 | 40.9982 | 46.8781 | 53.2142 | 57.7717 | 61.9615 | 68.1909 | 69.3297 | 75.7932 |
| 383.03 | 23.3333 | 32.0973 | 43.088 | 48.8441 | 55.9294 | 59.7428 | 64.4142 | 72.02 | 75.5208 | 78.587 |
| 383.93 | 23.7223 | 34.8101 | 42.6892 | 50.5845 | 58.4179 | 60.2878 | 66.2567 | 74.1876 | 77.8321 | 80.6511 |
| 385 | 24.0964 | 35.2099 | 47.0701 | 52.1466 | 57.9793 | 63.5401 | 71.3076 | 74.5796 | 79.0972 | 82.0622 |
| 386.06 | 25.2887 | 35.5669 | 45.794 | 53.9019 | 60.4923 | 65.3305 | 72.3824 | 76.4936 | 80.4312 | 83.9147 |
| 386.96 | 25.3219 | 36.5784 | 47.7809 | 54.621 | 60.9137 | 66.8957 | 73.7129 | 77.4023 | 83.0325 | 86.8315 |
| 388.03 | 28.3151 | 39.6857 | 49.8401 | 56.9979 | 62.5 | 70.4846 | 75.3821 | 82.0714 | 87.5478 | 91.1533 |
| 388.93 | 28.0106 | 39.0356 | 49.3803 | 57.4577 | 63.5759 | 71.5995 | 78.1728 | 83.5549 | 87.6454 | 90.8566 |
| 390 | 29.9479 | 39.6684 | 52.4981 | 56.7403 | 66.41 | 71.5702 | 79.7594 | 87.0836 | 91.9516 | 93.2746 |
| 391.06 | 30.7627 | 40.8038 | 52.2502 | 58.778 | 67.9327 | 71.7193 | 81.0338 | 88.8805 | 92.207 | 95.1336 |
| 391.96 | 28.8125 | 40.7523 | 54.5559 | 59.6437 | 70.3848 | 74.9257 | 80.0874 | 89.2755 | 90.5082 | 100.124 |
| 393.03 | 30.1811 | 41.7744 | 54.2797 | 61.6952 | 71.8028 | 76.0209 | 82.6217 | 90.7875 | 96.0884 | 98.7306 |
| 393.93 | 30.2843 | 41.6667 | 55.0758 | 60.6356 | 69.8117 | 78.5422 | 86.6172 | 88.9065 | 94.943 | 100.445 |
| 395 | 28.774 | 43.3354 | 56.213 | 61.6547 | 73.0612 | 79.7142 | 85.5188 | 93.5472 | 97.517 | 100.203 |
| 396.06 | 30.0508 | 42.8375 | 55.9723 | 65.3227 | 75.3915 | 77.9483 | 85.4836 | 91.6587 | 98.4037 | 100.524 |
| 396.96 | 31.2902 | 42.9633 | 56.2934 | 64.3105 | 72.5309 | 80.1426 | 87.8653 | 92.1181 | 97.3325 | 102.944 |
| 398.03 | 31.667 | 45.0672 | 55.9686 | 63.1822 | 73.212 | 80.0938 | 88.4739 | 92.6745 | 98.8839 | 104.708 |
| 398.93 | 33.4773 | 44.4399 | 57.7895 | 65 | 73.5871 | 78.4725 | 87.8951 | 96.7125 | 100.04 | 103.86 |
| 400 | 31.0797 | 45.7571 | 57.8299 | 65.1268 | 72.6994 | 81.4628 | 88.6355 | 95.3193 | 100.462 | 105.212 |
| 401.07 | 34.1129 | 45.901 | 57.8171 | 65.8794 | 73.8916 | 83.0578 | 89.2929 | 96.9015 | 98.6028 | 104.492 |
| 402 | 34.0265 | 45.3208 | 57.9432 | 66.6128 | 73.5385 | 83.9145 | 87.9017 | 97.4013 | 98.8852 | 104.583 |
| 403.07 | 32.9261 | 46.1656 | 59.7266 | 64.105 | 75.3271 | 80.7748 | 90.1558 | 95.2772 | 98.6434 | 103.56 |
| 404 | 31.314 | 45.5178 | 57.9512 | 62.6115 | 74.9469 | 81.6243 | 89.3551 | 94.6391 | 99.7856 | 103.956 |
| 405.07 | 30.0524 | 44.5191 | 57.2946 | 65.8767 | 75.6285 | 79.1322 | 88.0491 | 96.1464 | 100 | 101.35 |
| 406 | 33.3866 | 43.9628 | 58.4985 | 66.8826 | 74.4222 | 80.415 | 86.9815 | 91.9169 | 97.9947 | 103.817 |
| 407.07 | 32.3241 | 43.9865 | 56.6465 | 63.8459 | 71.6344 | 79.1941 | 85.1064 | 94.5946 | 95.8654 | 103.316 |
| 408 | 31.6495 | 45.102 | 56.8693 | 63.5152 | 71.9368 | 80.5153 | 85.4456 | 89.8467 | 96.2094 | 99.3415 |
| 409.07 | 31.1656 | 44.2863 | 55.0996 | 62.9186 | 73.1612 | 78.5449 | 86.6318 | 92.8858 | 98.9165 | 100.923 |
| 410 | 30.9239 | 44.2135 | 55.9399 | 62.6877 | 71.0151 | 75.884 | 82.7613 | 92.0175 | 95.4783 | 100.155 |
| 411.06 | 29.6264 | 43.2307 | 56.0897 | 63.3721 | 70.3864 | 75.2437 | 87.2964 | 90.8569 | 94.3659 | 98.8989 |
| 411.96 | 29.2445 | 44.1301 | 55.0477 | 62.0338 | 70.9727 | 76.3068 | 82.4781 | 89.1619 | 95.0128 | 96.6314 |
| 413.03 | 30.1455 | 42.9978 | 54.8051 | 60.9341 | 72.2509 | 75.0894 | 81.044 | 88.4219 | 93.443 | 94.9257 |
| 413.93 | 30.5556 | 41.0648 | 52.2081 | 61.8639 | 69.28 | 72.5146 | 80.6452 | 86.2358 | 90.3088 | 95.3366 |
| 415 | 30.8826 | 40.5434 | 53.2319 | 59.2157 | 67.2679 | 75.9298 | 80.185 | 86.9749 | 88.7143 | 92.7101 |
| 416.06 | 30.3772 | 40.9345 | 52.4375 | 59.7015 | 66.6401 | 72.0916 | 77.1033 | 81.6526 | 90.1528 | 89.9609 |
| 416.96 | 30 | 39.3509 | 52.1838 | 58.9724 | 67.2122 | 71.6523 | 79.0906 | 82.6787 | 87.1506 | 88.9483 |
| 418.03 | 27.2225 | 40.5083 | 51.4038 | 56.739 | 65.3707 | 68.31 | 74.2603 | 81.768 | 84.048 | 88.3956 |
| 418.93 | 29.4944 | 38.4235 | 49.4012 | 54.7596 | 64.5476 | 70.5763 | 74.9263 | 79.7821 | 84.7661 | 88.8007 |
| 420 | 28.1035 | 39.3814 | 49.472 | 54.0073 | 61.6291 | 67.1913 | 75.2918 | 80.6899 | 81.4773 | 84.6517 |
| 421.06 | 26.9608 | 37.037 | 47.7287 | 54.5847 | 58.5548 | 65.3101 | 73.8141 | 75.5483 | 81.3768 | 84.9645 |
| 421.96 | 24.395 | 34.7705 | 45.9343 | 51.6648 | 58.8818 | 64.6671 | 68.6818 | 74.5516 | 77.5116 | 80.2543 |
| 423.03 | 25.0293 | 35.4485 | 47.4354 | 50.8279 | 58.0384 | 61.9628 | 67.546 | 73.7523 | 75.5633 | 79.4118 |
| 423.93 | 26.0686 | 34.8074 | 43.8738 | 48.8962 | 57.561 | 58.4982 | 66.7337 | 71.0927 | 73.2323 | 76.7553 |
| 425 | 25.1559 | 34.0443 | 44.1291 | 47.8469 | 55.7665 | 59.2563 | 66.021 | 68.2977 | 70.2165 | 75.5158 |
| 426.06 | 23.9572 | 33.0594 | 42.9604 | 47.8252 | 53.0849 | 57.3561 | 61.4707 | 67.8978 | 68.2399 | 72.0047 |
| 426.96 | 23.3654 | 29.6821 | 39.6299 | 45.7644 | 50.9905 | 56.266 | 60.9565 | 65.4539 | 66.3395 | 69.6964 |
| 428.03 | 22.805 | 30.8833 | 37.8054 | 43.3532 | 50.0291 | 54.3838 | 57.6731 | 64.4974 | 65.7051 | 68.6616 |
| 428.93 | 22.0514 | 30.98 | 37.4143 | 43.8379 | 49.7339 | 52.1412 | 56.9075 | 62.1022 | 63.8385 | 66.9379 |
| 430 | 21.2559 | 29.9918 | 36.5217 | 41.3105 | 46.0462 | 51.2821 | 54.7136 | 58.6445 | 61.8357 | 63.9857 |
| 431.04 | 20.5785 | 28.6585 | 36.3248 | 40.5545 | 44.0671 | 51.357 | 55.2017 | 59.1054 | 60.2776 | 62.1782 |
| 431.94 | 20.2202 | 28.994 | 33.6614 | 38.9586 | 44.132 | 48.7136 | 50.5189 | 54.1796 | 57.1836 | 60.8763 |
| 432.98 | 19.042 | 24.741 | 33.4884 | 38.2227 | 42.2862 | 45.4824 | 49.8615 | 53.4125 | 55.4448 | 58.5534 |
| 434.02 | 19.7189 | 26.3817 | 31.9907 | 36.7534 | 42.1119 | 45.0432 | 49.3751 | 52.8541 | 53.0667 | 54.9494 |
| 435.07 | 17.2545 | 23.9198 | 32.0334 | 34.7515 | 38.6163 | 44.5434 | 45.6235 | 48.8794 | 51.7373 | 52.8309 |
| 435.97 | 17.2809 | 24.0265 | 29.4762 | 34.1817 | 38.4321 | 41.197 | 44.0495 | 48.1336 | 49.0368 | 53.8862 |
| 437.01 | 16.6961 | 22.8084 | 30.326 | 32.8131 | 37.9473 | 39.7062 | 44.511 | 45.1085 | 49.4221 | 48.3871 |
| 438.05 | 16.3599 | 22.5093 | 27.2602 | 32.9347 | 35.5949 | 38.717 | 41.0178 | 45.3317 | 47.4583 | 49.592 |
| 438.95 | 16.3067 | 21.6886 | 26.5918 | 31.0222 | 35.4932 | 37.7859 | 42.9805 | 44.3709 | 45.3926 | 46.4019 |
| 440 | 15.3051 | 21.9393 | 27.1226 | 29.3844 | 32.4346 | 36.2969 | 38.9352 | 42.9049 | 43.065 | 43.1625 |
| 441.06 | 15.3846 | 19.2487 | 24.0941 | 28.6101 | 31.7583 | 36.1305 | 37.7162 | 41.3091 | 41.4993 | 44.4851 |
| 441.96 | 14.1926 | 20.2079 | 25.1726 | 27.9776 | 30.8432 | 33.8846 | 39.3667 | 38.2795 | 39.5747 | 43.2149 |
| 443.03 | 13.2787 | 18.8024 | 25.1386 | 27.0912 | 30.6224 | 32.6594 | 36.1496 | 37.9432 | 40.1691 | 41.1342 |
| 443.93 | 15.019 | 19.7497 | 23.2059 | 25.4156 | 29.292 | 31.6973 | 35.014 | 35.5716 | 39.0254 | 39.3523 |
| 445 | 12.9819 | 18.0442 | 24.0889 | 24.0421 | 28.3077 | 27.9908 | 32.1054 | 35.4581 | 36.4963 | 37.2683 |
| 446.06 | 14.3334 | 18.8822 | 21.1913 | 23.0844 | 27.5075 | 30.404 | 30.8226 | 32.8629 | 33.3792 | 37.1589 |
| 446.96 | 14.49 | 17.386 | 21.8756 | 22.9949 | 24.5174 | 28.1325 | 31.3144 | 33.2909 | 33.5964 | 33.729 |
| 448.03 | 10.4392 | 16.3902 | 20.3995 | 22.4128 | 25.1864 | 27.0731 | 31.8846 | 30.7598 | 33.0706 | 35.0594 |
| 448.93 | 12.2687 | 15.4154 | 19.4777 | 21.0317 | 24.3471 | 26.6796 | 29.9939 | 31.9678 | 31.2384 | 33.9578 |
| 450 | 11.171 | 15.808 | 20.1581 | 21.6268 | 24.3619 | 24.665 | 26.763 | 29.1658 | 32.7969 | 31.3127 |
| 451.04 | 12.0951 | 14.3296 | 18.8865 | 19.7599 | 21.8835 | 23.8663 | 26.3959 | 27.4212 | 29.783 | 32.0929 |
| 451.94 | 11.9097 | 14.5522 | 17.7077 | 19.6348 | 22.052 | 22.2178 | 26.6747 | 28.6661 | 27.0593 | 30.6737 |
| 452.98 | 9.67867 | 13.2673 | 17.415 | 20.9551 | 21.878 | 22.5518 | 26.1242 | 26.6325 | 28.2362 | 30.467 |
| 454.02 | 10.1282 | 14.6158 | 16.8248 | 20.1937 | 19.865 | 22.4206 | 24.9705 | 27.3101 | 25.8295 | 28.0356 |
| 455.07 | 11.4808 | 12.897 | 17.5722 | 19.5205 | 20.7222 | 22.1282 | 22.7696 | 24.7305 | 25.0104 | 24.9948 |
| 455.97 | 9.06931 | 12.1902 | 15.5776 | 17.0801 | 20.2665 | 22.3152 | 23.7288 | 26.7037 | 23.6422 | 25.9793 |
| 457.01 | 9.20486 | 10.582 | 15.8954 | 16.2752 | 20.1724 | 20.7917 | 22.6843 | 25.5211 | 24.9291 | 25.8881 |
| 458.05 | 9.09826 | 12.5871 | 14.065 | 15.2006 | 19.4346 | 19.3563 | 21.6512 | 23.0296 | 23.6663 | 23.4815 |
| 458.95 | 9.95316 | 11.4481 | 13.922 | 16.4134 | 16.9162 | 18.9401 | 20.1473 | 22.7273 | 22.6097 | 23.2775 |
| 460 | 8.28893 | 11.5338 | 13.3042 | 15.1184 | 16.9188 | 18.6698 | 20.0696 | 21.1337 | 22.8898 | 22.8549 |
| 461.06 | 7.72052 | 10.2594 | 12.4363 | 15.2747 | 16.3871 | 18.5697 | 19.8974 | 20.1462 | 20.9985 | 22.115 |
| 461.96 | 8.30973 | 9.28558 | 12.3033 | 15.1745 | 16.7218 | 17.5811 | 17.8612 | 19.925 | 18.2346 | 21.7002 |
| 463.03 | 7.4727 | 9.69368 | 11.8952 | 14.2536 | 16.8196 | 15.0362 | 18.2612 | 18.4864 | 19.8858 | 19.5509 |
| 463.93 | 7.02888 | 11.1178 | 12.2213 | 15.296 | 13.7891 | 15.1806 | 16.9294 | 18.7512 | 17.3387 | 19.7636 |
| 465 | 8.09717 | 9.16684 | 11.5245 | 12.8426 | 12.8106 | 13.8728 | 16.0842 | 18.6298 | 17.3791 | 18.1784 |
| 466.06 | 7.55572 | 9.28026 | 12.7617 | 12.2618 | 13.0208 | 14.9611 | 16.9116 | 16.119 | 16.4633 | 18.9107 |
| 466.96 | 7.81403 | 9.38584 | 11.7165 | 11.0034 | 13.8862 | 13.0575 | 15.4135 | 16.7523 | 15.9542 | 17.12 |
| 468.03 | 6.37323 | 10.4764 | 10.3053 | 12.1335 | 11.9322 | 15.2381 | 15.2234 | 14.6616 | 14.8297 | 16.7142 |
| 468.93 | 6.35965 | 7.46697 | 9.50349 | 11.9121 | 11.5035 | 12.8866 | 13.6813 | 16.4601 | 15.0754 | 15.6689 |
| 470 | 5.82563 | 8.30973 | 8.87346 | 10.596 | 12.663 | 12.2272 | 14.7632 | 12.6432 | 14.6969 | 14.9374 |
| 471.04 | 3.70693 | 7.39196 | 7.77732 | 10.1493 | 10.0873 | 11.5628 | 14.269 | 13.8099 | 15.7015 | 15.4793 |
| 471.94 | 6.38298 | 8.17221 | 8.07882 | 10.8485 | 10.6404 | 13.1023 | 14.255 | 14.4503 | 14.1142 | 14.8456 |
| 472.98 | 6.71141 | 7.38007 | 8.15328 | 9.88593 | 13.0381 | 11.169 | 10.7092 | 12.2473 | 13.0061 | 14.9495 |
| 474.02 | 5.57585 | 8.79037 | 8.56233 | 9.09619 | 10.6362 | 11.4203 | 11.0837 | 11.0388 | 11.9686 | 12.0301 |
| 475.07 | 5.92441 | 4.73373 | 8.32318 | 7.95003 | 9.99623 | 10.3576 | 10.7271 | 11.8928 | 11.1896 | 13.5934 |
| 475.97 | 4.58453 | 7.13267 | 6.06673 | 8.08519 | 9.69337 | 10.1588 | 11.3752 | 12.3534 | 12.9241 | 10.2273 |
| 477.01 | 4.32366 | 7.27917 | 8.35247 | 7.564 | 9.89346 | 9.10408 | 9.30877 | 11.0694 | 11.1896 | 11.0141 |
| 478.05 | 3.56436 | 4.41205 | 6.58157 | 4.94853 | 9.44914 | 9.70874 | 9.51272 | 10.6789 | 10.7254 | 11.7976 |
| 478.95 | 4.28182 | 7.63359 | 5.91716 | 8.23124 | 9.40376 | 8.2218 | 9.59379 | 10.3339 | 9.96016 | 12.1855 |
| 480 | 3.97906 | 4.75813 | 6.03935 | 7.99065 | 7.78816 | 7.70548 | 11.4512 | 10.6446 | 9.61539 | 10.3832 |
| 481.04 | 4.01861 | 6.39659 | 7.34583 | 6.48203 | 8.12536 | 9.06583 | 8.77535 | 9.42308 | 9.69368 | 10.4503 |
| 481.94 | 3.85083 | 5.81508 | 7.26246 | 5.87232 | 7.5275 | 7.67165 | 9.1482 | 9.5498 | 9.90868 | 9.93789 |
| 482.98 | 3.12561 | 4.71787 | 5.70797 | 5.99729 | 6.69819 | 8.6495 | 9.6 | 8.8218 | 9.83731 | 8.67761 |
| 484.02 | 4.12574 | 4.75813 | 5.90872 | 6.10105 | 7.68049 | 7.14838 | 7.90514 | 8.96767 | 8.47131 | 8.22023 |
| 485.07 | 3.43053 | 5.62023 | 5.32133 | 6.23903 | 5.63721 | 6.97404 | 8.58455 | 7.7821 | 7.17794 | 8.72034 |
| 485.97 | 2.37581 | 6.67059 | 5.50032 | 6.45529 | 6.77245 | 6.83555 | 5.94097 | 8.61888 | 6.9129 | 7.31804 |
| 487.01 | 2.6455 | 3.88199 | 6.18302 | 5.923 | 6.07165 | 6.20124 | 7.03812 | 7.24161 | 9.32874 | 6.84671 |
| 488.05 | 4.72376 | 2.95421 | 5.3742 | 5.35988 | 5.98456 | 6.58404 | 6.93793 | 7.98722 | 7.05077 | 7.46983 |
| 488.95 | 3.48147 | 5.00289 | 3.54122 | 4.82024 | 6.77245 | 6.71872 | 6.07839 | 5.63326 | 7.41211 | 6.63027 |
| 490 | 4.06669 | 3.87372 | 5.81758 | 6.22865 | 5.27383 | 4.61983 | 5.79189 | 6.54892 | 7.74886 | 6.33641 |
| 491.04 | 4.7553 | 4.01826 | 3.91389 | 5.13452 | 6.21616 | 5.90752 | 4.90196 | 6.54098 | 5.16055 | 6.28698 |
| 491.94 | 3.08107 | 2.4372 | 3.86266 | 5.55213 | 5.07713 | 5.00109 | 5.004 | 6.76104 | 5.88235 | 4.66291 |
| 492.98 | 3.25932 | 2.44399 | 2.72603 | 4.15282 | 4.9948 | 5.18962 | 6.24874 | 3.91349 | 6.07787 | 6.45161 |
| 494.02 | 3.21932 | 4.13956 | 3.17902 | 4.47518 | 4.85732 | 4.31711 | 5.01157 | 6.51273 | 4.80962 | 5.93362 |
| 495.07 | 2.52918 | 3.30275 | 2.58141 | 2.9985 | 3.09598 | 4.67472 | 5.28885 | 5.14953 | 6.14226 | 5.49853 |
| 495.97 | 2.76419 | 2.44315 | 3.33611 | 4.36162 | 4.22565 | 5.66458 | 4.74744 | 4.77008 | 4.99616 | 5.3246 |
| 497.01 | 2.27618 | 3.2381 | 2.36407 | 4.47845 | 5.21485 | 3.5447 | 4.8714 | 5.89431 | 5.31371 | 4.4496 |
| 498.05 | 1.91286 | 3.04067 | 4.23158 | 5.00867 | 4.41678 | 3.71602 | 3.79317 | 5.33281 | 2.99222 | 5.48101 |
| 498.95 | 2.88958 | 3.0303 | 3.82602 | 4.13223 | 3.20034 | 3.61664 | 5.28169 | 4.96752 | 4.92308 | 3.51591 |
| 500 | 1.01605 | 3.52595 | 3.1428 | 4.05104 | 4.64936 | 3.24346 | 3.65455 | 3.98804 | 5.62233 | 4.03226 |
| 501.04 | 3.19729 | 2.53214 | 3.69578 | 2.77668 | 3.0525 | 4.60918 | 4.15266 | 5.10949 | 4.34611 | 4.02194 |
| 501.94 | 2.8132 | 2.87466 | 2.42915 | 2.83401 | 3.16832 | 4.76092 | 3.4463 | 4.95933 | 4.36335 | 5.38358 |
| 502.98 | 2.18558 | 1.98373 | 1.99402 | 4.47427 | 4.22184 | 4.63451 | 3.35289 | 3.83951 | 3.53945 | 3.4078 |
| 504.02 | 0.392234 | 1.76713 | 3.83065 | 2.27187 | 5.49659 | 3.44968 | 4.40529 | 2.99345 | 2.80843 | 3.58491 |
| 505.07 | 2.41206 | 2.51062 | 2.93071 | 2.30282 | 4.33412 | 2.25989 | 2.66362 | 3.89984 | 4.42567 | 1.56955 |
| 505.97 | 1.38696 | 2.46026 | 2.89615 | 2.48804 | 4.74777 | 3.65186 | 2.8238 | 4.38596 | 3.75869 | 4.36764 |
| 507.01 | 0.209336 | 2.56562 | 1.90621 | 3.18789 | 3.77145 | 3.30225 | 4.47036 | 4.39699 | 1.58165 | 3.85431 |
| 508.05 | 2.47808 | 2.20103 | 1.22649 | 2.97453 | 3.7422 | 3.24366 | 2.87301 | 4.66626 | 1.842 | 3.56139 |
| 508.95 | 1.20313 | 2.83661 | 3.003 | 2.27838 | 2.30326 | 4.26712 | 2.88684 | 3.37205 | 2.77118 | 2.60678 |
| 510 | 1.42945 | 0.387222 | 0.993641 | 2.52427 | 2.85275 | 2.73598 | 3.663 | 4.24825 | 2.1215 | 2.93902 |
| 511.04 | 1.92567 | 2.26291 | 1.75507 | 3.06122 | 2.71399 | 3.51975 | 2.429 | 3.15147 | 3.00075 | 2.80636 |
| 511.94 | 2.45793 | 1.15274 | 3.0448 | 2.0393 | 3.56918 | 1.65837 | 2.18081 | 3.31255 | 2.32693 | 2.3678 |
| 512.98 | 0.962279 | 1.59936 | 0 | 0.856348 | 1.41614 | 3.49176 | 2.41887 | 2.9274 | 2.33554 | 2.90135 |
| 514.02 | 1.19119 | 1.35344 | 3.26131 | 2.69542 | 2.645 | 2.22312 | 2.6716 | 1.85839 | 3.42792 | 1.89861 |
| 515.07 | 2.0202 | 1.39359 | 2.09996 | 0.801122 | 1.75507 | 2.26543 | 3.82922 | 2.63108 | 2.0496 | 2.38569 |
| 515.97 | 1.92567 | 1.83636 | 1.83561 | 2.65144 | 2.08725 | 2.75808 | 2.44821 | 3.25421 | 2.47855 | 2.57324 |
| 517.01 | 1.5212 | 1.79283 | 1.63199 | 1.12444 | 1.01112 | 1.0661 | 3.01205 | 1.90694 | 0.804343 | 2.57579 |
| 518.05 | 1.752 | 1.80687 | 0.605205 | 1.93424 | 1.29618 | 1.55129 | 2.62732 | 2.81162 | 1.67255 | 1.79964 |
| 518.95 | 1.8943 | 2.31839 | 1.13982 | 2.36305 | 1.62999 | 2.18862 | 1.80542 | 2.38445 | 2.01045 | 2.28528 |
| 520 | 1.57698 | 0.396275 | 2.51942 | 0.411523 | 1.6691 | 0.9998 | 2.93198 | 3.79107 | 1.58667 | 1.52265 |
| 521.02 | 1.62767 | 0 | 1.84313 | 1.97628 | 1.19928 | 1.46781 | 1.17119 | 2.13344 | 1.67973 | 1.73678 |
| 522.05 | 1.04624 | 1.51918 | 1.15207 | 1.37795 | 2.46026 | 2.08768 | 0.96043 | 0.406669 | 1.9861 | 1.32853 |
| 522.94 | 2.54254 | 2.644 | 1.60546 | 2.45449 | 0.645995 | 0.390016 | 2.29314 | 2.64013 | 2.54952 | 2.28877 |
| 523.97 | 0.772648 | 1.19 | 0.398883 | 1.07921 | 2.27618 | 0 | 3.32019 | 0.578704 | 2.84306 | 1.74081 |
| 525 | 0.582637 | 1.91534 | 0.813339 | 1.63299 | 2.84495 | 1.31678 | 2.01694 | 1.17028 | 1.56311 | 2.06263 |
| 526.02 | 1.92456 | 1.17233 | 2.23577 | 2.37483 | 1.56372 | 1.60772 | 2.04613 | 1.7398 | 1.75473 | 1.48285 |
| 527.05 | 2.32513 | 1.51114 | 1.59299 | 1.06428 | 0.584112 | 1.96309 | 1.82076 | 1.51803 | 2.5082 | 1.51114 |
| 527.94 | 0.80016 | 1.78642 | 0.994431 | 1.70685 | 1.1818 | 1.455 | 2.25056 | 1.54859 | 1.74149 | 1.86811 |
| 528.97 | -0.210172 | 1.9044 | 0.784006 | 2.22627 | 1.73544 | 2.15813 | 0.952562 | 1.12317 | 1.90331 | 1.92308 |
| 530 | 1.00321 | 0.384246 | 2.63158 | 2.67105 | 0.769675 | 2.35571 | 2.29095 | 2.58956 | 1.56311 | 1.93199 |
| 531.04 | 1.60385 | 0.193349 | 1.4626 | 1.99005 | 2.03998 | 1.58636 | 0.781708 | 1.57233 | 3.19849 | 1.9425 |
| 531.94 | 0.783085 | 1.51688 | 0 | 1.89789 | 0.9994 | 0.995025 | 1.38861 | 2.16877 | 1.76574 | 0.770416 |
| 532.98 | 1.52934 | 1.35554 | 0.777001 | 1.56433 | 1.02838 | -0.404694 | 1.13186 | 1.63666 | 2.30725 | 0.591133 |
| 534.02 | 1.81928 | 1.36665 | 0.19972 | 1.63132 | 1.21139 | 0.781403 | 2.0496 | 1.91681 | 0 | 1.62075 |
| 535.07 | 1.002 | 2.33146 | 1.37255 | 0.38835 | 0.5923 | 1.88715 | 1.42045 | 1.9359 | 1.17925 | 0.390472 |
| 535.97 | -0.195848 | 1.20289 | 1.72745 | 1.01031 | 0.987557 | 2.91715 | 1.04058 | 2.19124 | 1.81378 | 0.611247 |
| 537.01 | 1.14416 | 0.189358 | 2.46399 | 1.01792 | 0.767902 | 1.04471 | 1.57604 | 1.47493 | 0.995619 | -0.194363 |
| 538.05 | 0.749625 | 0.575043 | 1.98491 | 0 | 0.405844 | 1.74723 | 1.90259 | 0.9802 | 0.756716 | 0.376577 |
| 538.95 | 0.976372 | 1.2109 | 2.0861 | 1.34693 | 1.38999 | 1.67007 | 0.403388 | 1.56771 | -0.377358 | 2.06806 |
| 540 | -0.9996 | 0.81666 | 0 | 1.39832 | 2.24765 | -0.383362 | 1.72377 | 1.76574 | 1.50574 | 2.93141 |
| 541.02 | 1.15585 | 1.92843 | -0.785238 | 1.02082 | 0.386548 | 2.39044 | 1.15741 | 1.22674 | 1.54919 | 0.779879 |
| 542.05 | 1.61453 | 1.50291 | 1.34641 | 0.21363 | 1.16777 | 1.00402 | 1.17716 | 1.58197 | 0.778816 | 0.959693 |
| 542.94 | 0.605938 | 1.36852 | 1.14591 | 1.85271 | 0.796813 | 0.596659 | -1.4917 | 1.27959 | 0.367512 | -0.196889 |
| 543.97 | 1.0006 | 0.200803 | 0.391313 | -0.979624 | -0.20674 | 0.596777 | 1.04406 | -0.383583 | 0.187758 | 0 |
| 545 | 1.33486 | 1.37174 | 0.189394 | 0.814498 | 0.200763 | 0.192308 | 1.38779 | 0.786782 | 0.400962 | 1.92308 |
| 546.02 | 0 | 0.583885 | 1.0018 | 0.959877 | 0.604351 | 0.593824 | 0.379867 | 0.390168 | 1.16369 | -0.197707 |
| 547.05 | 1.16868 | 0.945537 | 0.3803 | 1.22699 | 1.89072 | 0 | 0.413479 | 0 | -0.990099 | 0.564334 |
| 547.94 | 0 | 0.986777 | 1.17302 | 1.60643 | 1.20506 | 0.382336 | 1.01153 | 0.811853 | 0.595593 | 0.94038 |

**Fluorescence spectra of *N*^2^,*N*^6^-bis((6-(4-methylpiperazin-1-yl)-1*H*,3'*H*-[2,5'-dibenzo[*d*]imidazol]-2'-yl)methyl)pyridine-2,6-dicarboxamide (XI – DB_2_Py) binding to dsDNA**

DB2Py

Intensity, rel. units

| Wavelength, nm | 0 | 9 | 22 | 45 | 75 | 110 | 152 | 214 | 294 |
| --- | --- | --- | --- | --- | --- | --- | --- | --- | --- |
| 350 | 0.441696 | 0.402091 | 0.202429 | 0.812183 | 3.22646 | 1.98098 | 1.89681 | -0.390549 | 1.37174 |
| 351.07 | 0.402495 | 0.745295 | 0.61488 | 1.5447 | -0.199322 | 0.433088 | -0.558971 | 0.187266 | 0.949487 |
| 352 | 0.395648 | 0.19459 | 1.38313 | 0.612495 | 0.61792 | 0.945716 | 1.21212 | 0.777001 | 0.569044 |
| 353.07 | -0.820176 | -0.200723 | 1.74791 | 1.95656 | -0.654022 | 0.396511 | 1.10152 | 0.777303 | 1.14701 |
| 354 | 1.32601 | 1.15207 | 0.592768 | 2.24765 | 0.930406 | 0.992457 | 1.71331 | -0.182548 | 1.11132 |
| 355.07 | 0.841397 | 0.20475 | 0.385505 | 1.3245 | 1.42799 | 1.33894 | 0.967305 | -0.20008 | 1.53169 |
| 356 | -0.197902 | 1.39888 | 1.71821 | 0 | 0.959693 | 0.545355 | 1.13809 | 0.405104 | 0.926784 |
| 357.07 | 0.558659 | -0.401606 | 0.201207 | 1.48011 | 1.51286 | 0.750188 | 0.957488 | 2.35433 | 0.757432 |
| 358 | -0.591133 | 0.944287 | 0.205255 | 0.761325 | 1.61063 | 1.1396 | 0 | 1.98926 | 2.25394 |
| 359.07 | 0.418673 | -0.421852 | 0 | 1.23026 | 0.579262 | 1.90549 | 1.54589 | 1.88715 | 2.29183 |
| 360 | 1.3739 | 1.01256 | 0.4329 | 1.74452 | 0.960246 | 0.572628 | 2.29841 | 0.386399 | 0.957854 |
| 361.06 | 1.82964 | 0.786164 | -0.193498 | 1.24818 | 0.3996 | 1.33308 | 1.95503 | 2.01167 | 0.742115 |
| 361.96 | 0.622536 | 1.0227 | 2.14133 | 1.44152 | 0.962279 | 0.795545 | 1.95695 | 1.55794 | 0.757576 |
| 363.03 | 0.21106 | 1.34901 | 2.27747 | 1.48285 | 1.13593 | 1.11982 | 1.61193 | 2.00562 | 0.396354 |
| 363.93 | 0.391619 | 0 | 1.38806 | 1.33055 | 0.193573 | 1.79605 | 2.3037 | 2.32648 | 2.69334 |
| 365 | 0.767607 | 0.758869 | 2.62025 | 2.15602 | 0.396589 | 2.76571 | 1.13593 | 0 | 2.01687 |
| 366.06 | 1.53728 | 1.76091 | 2.32648 | 1.97161 | 0.977135 | 2.05719 | 1.57853 | 0.792864 | 1.79247 |
| 366.96 | 0.91954 | 2.18124 | 2.936 | 1.32375 | 1.01276 | 1.34357 | 1.7769 | 1.66976 | 1.89072 |
| 368.03 | 1.06022 | 0.594648 | 1.75747 | 1.37633 | 1.94666 | 1.55099 | 2.76352 | 2.85714 | 1.90621 |
| 368.93 | 1.24662 | 0.194856 | 1.57202 | 2.52036 | 2.93485 | 1.90913 | 3.27427 | 2.38949 | 3.05285 |
| 370 | 1.43003 | 0.63012 | 1.62602 | 2.65604 | 2.06108 | 2.39569 | 2.47627 | 1.74927 | 3.62538 |
| 371.07 | 0.429092 | 1.22299 | 1.75713 | 1.15274 | 0.594648 | 2.30194 | 3.23995 | 3.23071 | 2.58924 |
| 372 | 0.657318 | 2.73705 | 2.22672 | 2.83 | 5.26111 | 2.42229 | 3.71239 | 4.08481 | 2.41367 |
| 373.07 | 1.08672 | 1.67961 | 2.98211 | 2.53325 | 3.44828 | 2.65135 | 3.06455 | 2.29533 | 4.1849 |
| 374 | 1.06496 | 1.65187 | 2.92626 | 1.95122 | 3.35173 | 3.76014 | 4.04002 | 4.34783 | 4.51423 |
| 375.07 | -0.440626 | 1.01605 | 2.32313 | 2.87062 | 3.68928 | 3.4116 | 3.09534 | 3.84615 | 4.5977 |
| 376 | 0.59976 | -0.837696 | 3.45599 | 2.72798 | 2.68097 | 5.3903 | 4.49714 | 3.61079 | 5.75484 |
| 377.07 | 1.67154 | 2.20751 | 1.74419 | 2.66612 | 3.15048 | 5.2184 | 5.02804 | 5.59105 | 4.85437 |
| 378 | 1.9938 | 0.407747 | 4.22705 | 3.94867 | 4.35956 | 5.52704 | 6.34921 | 4.90659 | 6.06298 |
| 379.07 | 4.17674 | 4.16575 | 4 | 4.18577 | 5.56328 | 5.61363 | 4.02916 | 7.19697 | 6.63674 |
| 380 | 1.83038 | 3.22165 | 4.36681 | 4.40529 | 4.67017 | 6.01564 | 6.56537 | 7.36822 | 5.78381 |
| 381.06 | 0.811853 | 3.06547 | 4.52396 | 5.26427 | 6.47723 | 7.49211 | 9.00728 | 7.04955 | 7.10992 |
| 381.96 | 1.76922 | 3.57782 | 4.77802 | 6.71961 | 7.2549 | 6.98742 | 9.38158 | 6.89655 | 9.54563 |
| 383.03 | 2.45449 | 3.40368 | 5.26316 | 6.48692 | 9.20737 | 9.90099 | 9.78857 | 10.9105 | 9.21572 |
| 383.93 | 2.34842 | 3.57143 | 5.85774 | 7.53411 | 9.02346 | 9.56284 | 9.88069 | 12.5847 | 10.097 |
| 385 | 2.44101 | 5.00435 | 5.90476 | 8.87634 | 9.19188 | 10.7025 | 11.4723 | 12.3984 | 12.9113 |
| 386.06 | 2.15936 | 3.95175 | 5.53646 | 10.4551 | 10.1585 | 13.392 | 13.2915 | 12.4402 | 14.9282 |
| 386.96 | 1.64508 | 3.71098 | 7.39675 | 9.17957 | 12.7255 | 14.4789 | 14.8865 | 14.723 | 15.3729 |
| 388.03 | 3.57356 | 5.74332 | 8.12536 | 9.94346 | 12.188 | 14.1762 | 16.7986 | 16.3175 | 16.8199 |
| 388.93 | 2.31036 | 6.08382 | 8.0048 | 9.73654 | 13.8462 | 16.5735 | 17.5073 | 18.5261 | 19.7292 |
| 390 | 4.83189 | 5.9629 | 9.27419 | 13.5545 | 16.1449 | 17.7241 | 18.9584 | 20.8211 | 21.6061 |
| 391.06 | 3.50154 | 7.64169 | 10.6623 | 13.1706 | 17.5041 | 19.6721 | 21.1308 | 22.1205 | 22.6828 |
| 391.96 | 4.83969 | 6.77105 | 11.0906 | 14.4544 | 18.3164 | 21.3415 | 23.3733 | 23.1463 | 23.7014 |
| 393.03 | 5.00939 | 7.16393 | 11.0954 | 15.7709 | 19.6641 | 23.0681 | 25.8746 | 26.452 | 27.3416 |
| 393.93 | 4.29799 | 8.02026 | 12.6453 | 18.1499 | 23.4392 | 26.6586 | 29.2208 | 27.9193 | 30.4276 |
| 395 | 5.97076 | 9.64912 | 14.1343 | 21.1586 | 23.9892 | 27.2374 | 29.3166 | 30.6542 | 31.4453 |
| 396.06 | 5.3509 | 8.65801 | 16.798 | 21.8056 | 26.0786 | 29.7313 | 31.9984 | 35.822 | 36.1232 |
| 396.96 | 6.51659 | 11.795 | 15.7698 | 24.2631 | 30.4966 | 31.1292 | 35.4017 | 36.5036 | 36.2592 |
| 398.03 | 6.78558 | 12.2229 | 18.2728 | 25.6972 | 31.2437 | 33.9682 | 37.5387 | 39.1937 | 39.2118 |
| 398.93 | 7.00309 | 14.1549 | 17.9806 | 26.1758 | 33.6254 | 37.3039 | 40.8951 | 41.907 | 41.5624 |
| 400 | 8.86382 | 11.5013 | 20.5243 | 29.5527 | 33.7797 | 39.8104 | 41.6091 | 44.7297 | 45.9969 |
| 401.07 | 10.2605 | 15.9628 | 23.0074 | 31.0198 | 39.3652 | 42.7367 | 45.2552 | 46.9575 | 50.2977 |
| 402 | 9.46697 | 14.2268 | 22.8113 | 30.5135 | 39.2078 | 44.3128 | 47.1284 | 49.8887 | 49.094 |
| 403.07 | 11.5418 | 15.6552 | 23.9215 | 33.7641 | 43.4615 | 48.1714 | 50.9943 | 54.842 | 55.9694 |
| 404 | 11.6149 | 16.5149 | 27.5932 | 36.3565 | 44.0327 | 49.443 | 53.5609 | 54.0441 | 57.6963 |
| 405.07 | 11.4781 | 18.9416 | 27.0434 | 37.0937 | 47.0699 | 52.538 | 56.7333 | 59.7484 | 61.5647 |
| 406 | 14.2794 | 20.8459 | 27.9041 | 38.1584 | 47.5803 | 54.4335 | 59.7015 | 61.2011 | 63.9218 |
| 407.07 | 14.824 | 21.3232 | 30.8793 | 42.5816 | 52.0668 | 59.5736 | 60.8531 | 65.6163 | 68.0808 |
| 408 | 15.252 | 22.9909 | 32.2721 | 42.144 | 53.8205 | 59.4315 | 62.0843 | 66.211 | 69.2972 |
| 409.07 | 16.938 | 24.53 | 35.5006 | 45.5117 | 55.7191 | 61.3042 | 66.417 | 69.4523 | 72.1454 |
| 410 | 19.6123 | 26.0813 | 36.0771 | 49.3448 | 56.9647 | 64.8944 | 70.5006 | 72.171 | 75.4053 |
| 411.06 | 19.3522 | 25.9519 | 37.961 | 49.6012 | 61.7619 | 66.949 | 72.2527 | 77.1267 | 79.8169 |
| 411.96 | 18.9623 | 30.1706 | 38.8309 | 53.2779 | 64.3394 | 69.777 | 75.1822 | 77.1903 | 82.7874 |
| 413.03 | 22.9136 | 31.083 | 43.0192 | 54.4654 | 63.6854 | 73.9224 | 79.6443 | 81.8669 | 85.7308 |
| 413.93 | 23.2358 | 32.5238 | 42.5661 | 57.5397 | 68.1314 | 73.4844 | 77.7864 | 84.4037 | 86.0432 |
| 415 | 24.8998 | 33.0198 | 44.1506 | 58.9555 | 70.5657 | 77.2162 | 83.1899 | 85.3682 | 87.9765 |
| 416.06 | 27.1484 | 36.3002 | 46.0775 | 60.9831 | 73.0584 | 79.17 | 84.4991 | 88.0269 | 88.0569 |
| 416.96 | 30.5898 | 36.9526 | 49.6089 | 62.1709 | 73.4548 | 81.9703 | 88.7433 | 92.0574 | 93.1071 |
| 418.03 | 31.1771 | 39.5334 | 51.2821 | 65.2914 | 78.074 | 84.5872 | 89.1795 | 91.3859 | 97.3309 |
| 418.93 | 31.2775 | 40.2174 | 50.5366 | 64.2887 | 78.2889 | 85.1626 | 92.8489 | 96.1943 | 98.3501 |
| 420 | 33.7767 | 42.6312 | 52.4787 | 66.9184 | 81.1823 | 91.0754 | 91.6517 | 95.8771 | 100 |
| 421.06 | 35.9195 | 44.1044 | 54.8805 | 72.4903 | 81.8458 | 89.0752 | 94.9981 | 100.02 | 103.586 |
| 421.96 | 37.105 | 45.2536 | 59.1679 | 72.8605 | 83.3654 | 91.7281 | 95.7752 | 99.6427 | 101.287 |
| 423.03 | 39.5432 | 50.1062 | 58.657 | 73.8531 | 85.209 | 92.8003 | 100.5 | 100.775 | 106.428 |
| 423.93 | 40.9483 | 48.8273 | 60.7987 | 74.9331 | 87.2778 | 95.5292 | 100.941 | 105.814 | 105.909 |
| 425 | 43.3532 | 50.4076 | 63.5885 | 76.1711 | 86.8818 | 95.3815 | 101.963 | 106.079 | 106.595 |
| 426.06 | 44.0113 | 53.1526 | 64.6154 | 77.9409 | 90.5697 | 97.9576 | 103.278 | 105.892 | 112.896 |
| 426.96 | 46.2428 | 55.7392 | 64.7957 | 79.9087 | 90.6213 | 98.8896 | 104.37 | 105.081 | 109.277 |
| 428.03 | 50.4062 | 57.2189 | 65.2783 | 78.6968 | 93.3914 | 98.3362 | 103.529 | 107.274 | 109.447 |
| 428.93 | 50.1833 | 58.2667 | 66.9243 | 82.1624 | 93.4543 | 101.751 | 105.369 | 106.455 | 110.726 |
| 430 | 54.0367 | 60.9351 | 69.5251 | 81.108 | 95.3144 | 101.182 | 106.009 | 108.797 | 113.171 |
| 431.04 | 53.9872 | 59.466 | 69.9259 | 86.0899 | 93.7264 | 101.699 | 103.811 | 107.805 | 111.95 |
| 431.94 | 56.8134 | 64.1278 | 71.756 | 82.2528 | 95.4373 | 104.167 | 105.916 | 109.81 | 111.295 |
| 432.98 | 58.4857 | 63.9509 | 74.375 | 84.2233 | 94.4913 | 100.104 | 106.618 | 108.508 | 109.647 |
| 434.02 | 60.0588 | 67.3964 | 75.4912 | 87.5986 | 96.5275 | 100.461 | 107.843 | 108.983 | 111.668 |
| 435.07 | 60.0517 | 67.0426 | 78.534 | 83.8671 | 96.9866 | 103.455 | 107.317 | 109.749 | 111.241 |
| 435.97 | 63.9143 | 69.5351 | 75.7912 | 86.6239 | 98.4552 | 102.996 | 105.365 | 109.73 | 109.444 |
| 437.01 | 64.6412 | 69.4997 | 75.3147 | 88.4778 | 97.8044 | 102.486 | 106.16 | 110.328 | 111.68 |
| 438.05 | 67.8668 | 68.1916 | 79.9379 | 87.6299 | 96.0673 | 103.483 | 105.67 | 108.954 | 109.922 |
| 438.95 | 69.2308 | 74.8663 | 80.5829 | 91.4647 | 97.4886 | 101.744 | 106.531 | 106.758 | 107.901 |
| 440 | 70.5332 | 73.7115 | 82.7867 | 91.9893 | 98.4353 | 101.927 | 105.438 | 105.528 | 109.813 |
| 441.06 | 75.058 | 78.627 | 80.1394 | 88.7446 | 98.2881 | 99.7432 | 104.869 | 107.941 | 109.342 |
| 441.96 | 73.8015 | 77.4984 | 84.4371 | 91.7778 | 99.3095 | 103.742 | 104.208 | 108.241 | 109.951 |
| 443.03 | 76.1561 | 77.1355 | 84.9174 | 92.4722 | 98.899 | 104.659 | 105.893 | 110.913 | 110.918 |
| 443.93 | 78.9998 | 80.4361 | 86.9835 | 91.3533 | 99.7666 | 102.166 | 107.023 | 105.956 | 106.203 |
| 445 | 81.803 | 81.9996 | 87.8989 | 93.9346 | 100.802 | 102.644 | 105.866 | 106.329 | 105.792 |
| 446.06 | 82.4834 | 84.2311 | 86.6248 | 92.5926 | 99.1311 | 102.575 | 105.619 | 104.942 | 106.656 |
| 446.96 | 84.8383 | 84.6517 | 90.5615 | 91.8693 | 99.2963 | 105.24 | 104.909 | 105.578 | 106.901 |
| 448.03 | 84.9016 | 85.1648 | 88.4921 | 96.8188 | 97.3134 | 101.558 | 104.315 | 103.556 | 106.444 |
| 448.93 | 87.8364 | 90.9472 | 92.7379 | 96.3575 | 101.056 | 102.897 | 103.23 | 105.489 | 106.961 |
| 450 | 87.5645 | 89.795 | 93.7881 | 97.5851 | 100.777 | 102.142 | 102.062 | 103.038 | 106.396 |
| 451.04 | 89.1833 | 91.7118 | 95.835 | 96.6805 | 100.328 | 102.861 | 103.186 | 104.218 | 103.095 |
| 451.94 | 91.4597 | 92.5708 | 94.8086 | 97.2194 | 98.6307 | 103.144 | 104.348 | 104.843 | 103.567 |
| 452.98 | 95.9285 | 93.2643 | 95.4057 | 97.7205 | 101.559 | 101.355 | 102.744 | 104.694 | 105.23 |
| 454.02 | 94.6422 | 95.825 | 95.974 | 99.1427 | 99.108 | 101.073 | 104.261 | 104.514 | 103.442 |
| 455.07 | 96.7289 | 98.3948 | 97.8662 | 100.671 | 100.951 | 104.016 | 102.142 | 100.84 | 100.628 |
| 455.97 | 100.235 | 99.65 | 100.201 | 100.46 | 103.185 | 99.9176 | 99.8562 | 101.645 | 100.593 |
| 457.01 | 100.347 | 102.285 | 97.9489 | 99.7265 | 102.02 | 101.182 | 100.242 | 102.908 | 101.371 |
| 458.05 | 102.925 | 102.095 | 101.88 | 98.5045 | 99.6063 | 99.4085 | 99.7685 | 99.7061 | 99.7299 |
| 458.95 | 102.935 | 99.012 | 103.089 | 100.552 | 102.266 | 101.366 | 100.066 | 100.54 | 98.9289 |
| 460 | 104.837 | 102.071 | 100.84 | 101.801 | 102.504 | 100.039 | 100.593 | 99.3417 | 96.4544 |
| 461.06 | 107.071 | 102.388 | 101.82 | 105.509 | 97.7254 | 99.4387 | 97.1392 | 97.0429 | 97.5844 |
| 461.96 | 106.705 | 104.478 | 102.806 | 100.849 | 99.2481 | 100.136 | 98.4864 | 97.3522 | 96.6144 |
| 463.03 | 106.759 | 105.575 | 102.554 | 103.54 | 99.9566 | 98.1438 | 95.9533 | 95.5272 | 94.6825 |
| 463.93 | 109.043 | 102.729 | 104.095 | 100.594 | 98.1886 | 97.7076 | 96.6074 | 97.2139 | 95.3774 |
| 465 | 110.626 | 107.12 | 102.361 | 100.912 | 99.8623 | 95.8473 | 94.6795 | 93.968 | 92.4233 |
| 466.06 | 111.832 | 106.324 | 104.863 | 99.7338 | 98.768 | 94.943 | 93.3175 | 91.621 | 89.9408 |
| 466.96 | 111.281 | 106.807 | 103.595 | 98.9952 | 97.4843 | 94.7149 | 94.5153 | 92.0784 | 89.4823 |
| 468.03 | 110.493 | 109.398 | 104.964 | 98.586 | 96.6845 | 95.1309 | 96.5153 | 89.1655 | 87.7364 |
| 468.93 | 114.077 | 109.01 | 105.232 | 99.5466 | 95.0059 | 91.2498 | 91.0691 | 86.2841 | 87.3261 |
| 470 | 113.854 | 105.88 | 103.456 | 98.4221 | 95.1973 | 91.0509 | 88.978 | 87.9211 | 85.9328 |
| 471.04 | 113.177 | 108.989 | 101.645 | 97.456 | 94.2597 | 90.5818 | 87.9724 | 85.0347 | 84.2803 |
| 471.94 | 114.396 | 107.855 | 103.523 | 96.395 | 93.8444 | 91.2617 | 86.8083 | 85.8119 | 82.147 |
| 472.98 | 115.215 | 110.138 | 101.188 | 96.9574 | 93.7312 | 88.4782 | 85.9085 | 84.1463 | 81.6327 |
| 474.02 | 113.46 | 109.871 | 102.293 | 95.6337 | 89.2136 | 88.7779 | 83.8926 | 82.302 | 77.3939 |
| 475.07 | 113.742 | 108.176 | 100.591 | 96.3991 | 89.8808 | 84.1103 | 83.9363 | 79.5686 | 78.9523 |
| 475.97 | 112.869 | 105.989 | 101.606 | 95.7811 | 87.9666 | 84.9282 | 81.5271 | 77.0038 | 76.8623 |
| 477.01 | 112.128 | 105.576 | 97.4138 | 93.6448 | 86.5228 | 83.4187 | 78.1563 | 78.0661 | 76.7997 |
| 478.05 | 112.769 | 107.18 | 100.846 | 91.3471 | 84.7835 | 79.6355 | 78.1341 | 74.9806 | 72.9393 |
| 478.95 | 113.092 | 108.126 | 98.8474 | 92.0691 | 84.255 | 79.4626 | 76.5125 | 76.2887 | 72.5977 |
| 480 | 112.395 | 105.647 | 99.5084 | 92.8074 | 83.623 | 78.2813 | 75.4753 | 72.5318 | 73.2818 |
| 481.04 | 110.881 | 105.716 | 97.2107 | 89.741 | 81.9087 | 78.125 | 74.6969 | 69.6496 | 68.9167 |
| 481.94 | 110.902 | 101.997 | 96.4416 | 89.3695 | 84.629 | 77.8656 | 73.6158 | 70.4075 | 68.7453 |
| 482.98 | 111.608 | 104.019 | 96.9913 | 86.5424 | 80.2196 | 75.431 | 72.571 | 66.7992 | 68.0537 |
| 484.02 | 111.513 | 103.323 | 95.378 | 86.4877 | 77.8234 | 74.7556 | 70.9557 | 65.641 | 63.3789 |
| 485.07 | 109.742 | 102.304 | 95.6448 | 87.5206 | 78.6325 | 72.4019 | 68.5526 | 64.1227 | 63.6219 |
| 485.97 | 106.207 | 101.931 | 95.1802 | 84.5933 | 77.504 | 70.1896 | 68.109 | 64.3518 | 62.4242 |
| 487.01 | 109.154 | 100.193 | 91.3197 | 84.5499 | 77.0119 | 71.5388 | 65.91 | 62.4741 | 58.6792 |
| 488.05 | 108.471 | 99.2848 | 91.9176 | 82.3224 | 75.8608 | 67.2844 | 66.3006 | 60.7747 | 58.2109 |
| 488.95 | 106.772 | 101.931 | 89.372 | 79.9008 | 73.0541 | 67.28 | 63.3467 | 62.4474 | 58.9893 |
| 490 | 107.64 | 97.7606 | 89.6101 | 80.9496 | 69.2641 | 66.1014 | 62.2783 | 60.7646 | 55.0612 |
| 491.04 | 104.217 | 96.7101 | 90.0791 | 80.6758 | 72.4435 | 67.8939 | 61.5021 | 58.3063 | 54.4351 |
| 491.94 | 102.444 | 97.3712 | 87.1541 | 75.8099 | 69.7387 | 64.2165 | 60.3664 | 56.7436 | 54.6082 |
| 492.98 | 101.546 | 93.6615 | 87.3345 | 76.9861 | 69.6141 | 63.0169 | 59.5593 | 55.9739 | 53.3386 |
| 494.02 | 102.923 | 92.3845 | 87.2375 | 75.4502 | 66.9284 | 62.2187 | 55.6921 | 51.503 | 50.9194 |
| 495.07 | 101.314 | 94.3779 | 82.4387 | 75.4989 | 64.1359 | 60.6472 | 56.6268 | 52.2158 | 52.2057 |
| 495.97 | 99.7346 | 92.2122 | 83.2043 | 74.4724 | 66.1221 | 58.4708 | 54.5073 | 52.2876 | 48.2759 |
| 497.01 | 97.1625 | 90.8525 | 83.7876 | 71.8118 | 62.7865 | 56.3143 | 55.8346 | 50.7103 | 48.3678 |
| 498.05 | 100.96 | 90.224 | 81.9705 | 72.7381 | 61.4621 | 56.4021 | 52.5288 | 50.7412 | 46.7824 |
| 498.95 | 96.8917 | 88.8078 | 80.8364 | 69.1901 | 61.256 | 54.3267 | 52.6626 | 48.6994 | 45.1026 |
| 500 | 95.7698 | 84.6056 | 80.0582 | 69.3224 | 61.1855 | 54.6875 | 50.2434 | 46.4588 | 45.1211 |
| 501.04 | 94.2475 | 86.6734 | 76.7736 | 67.3956 | 59.905 | 52.35 | 49.2889 | 45.954 | 42.6602 |
| 501.94 | 91.5825 | 86.1362 | 74.8234 | 67.7999 | 59.0006 | 52.1621 | 46.3745 | 44.0341 | 41.3659 |
| 502.98 | 89.8901 | 85.5638 | 73.9747 | 70.8613 | 58.1197 | 50.7701 | 47.1718 | 44.4843 | 41.0586 |
| 504.02 | 90.9091 | 82.9431 | 75.9469 | 65.0343 | 55.1088 | 48.5456 | 44.6321 | 42.1343 | 39.0791 |
| 505.07 | 88.6273 | 80.2964 | 73.6365 | 62.6094 | 55.11 | 47.957 | 44.2395 | 39.4432 | 40.4398 |
| 505.97 | 89.254 | 81.1786 | 71.112 | 63.6308 | 53.7873 | 47.4187 | 43.9901 | 39.62 | 37.6303 |
| 507.01 | 84.9118 | 79.085 | 71.4286 | 60.1018 | 50.8376 | 47.5538 | 44.8328 | 39.397 | 37.4008 |
| 508.05 | 82.9398 | 80.7799 | 67.4543 | 59.5794 | 50.4803 | 45.0874 | 42.2694 | 37.8264 | 35.3014 |
| 508.95 | 84.122 | 75.2117 | 68.453 | 57.0647 | 49.7121 | 42.9185 | 41.562 | 36.7126 | 36.1809 |
| 510 | 83.4097 | 76.2748 | 68.2048 | 58.5075 | 48.3578 | 43.5377 | 41.6418 | 38.3505 | 34.3985 |
| 511.04 | 82.9268 | 73.494 | 63.745 | 57.087 | 46.7405 | 42.3511 | 39.3379 | 34.8793 | 33.276 |
| 511.94 | 76.7828 | 70.7351 | 65.9764 | 55.1213 | 45.3641 | 41.1038 | 37.4674 | 33.6215 | 32.4461 |
| 512.98 | 77.821 | 70.3141 | 64.9269 | 53.6595 | 46.9112 | 40.437 | 35.8753 | 34.1255 | 30.7785 |
| 514.02 | 75.9331 | 70.4168 | 61.7046 | 53.294 | 45.005 | 37.322 | 35.6374 | 32.2178 | 30.2907 |
| 515.07 | 72.6269 | 70.2079 | 61.0465 | 52.755 | 44.0385 | 37.6949 | 36.3191 | 33.6969 | 30.9779 |
| 515.97 | 74.7886 | 66.8943 | 58.4126 | 50.6353 | 43.1994 | 37.234 | 33.7447 | 30.9902 | 28.1632 |
| 517.01 | 72.6954 | 67.0482 | 58.0383 | 50.6849 | 43.2087 | 35.03 | 32.0061 | 28.8328 | 27.7886 |
| 518.05 | 72.5859 | 66.0377 | 57.6303 | 48.7568 | 40.7684 | 35.5981 | 33.0243 | 28.6811 | 27.6469 |
| 518.95 | 70.3593 | 63.9951 | 56.9718 | 46.8009 | 39.7656 | 34.7724 | 31.6027 | 29.877 | 25.1981 |
| 520 | 70.1496 | 62.2797 | 56.4071 | 46.1718 | 40.1318 | 34.3573 | 30.4666 | 28.904 | 25.2665 |
| 521.02 | 67.4595 | 62.0063 | 56.146 | 45.4079 | 37.9474 | 32.8215 | 31.7236 | 25.641 | 23.9521 |
| 522.05 | 69.0299 | 60.8215 | 54.5244 | 44.9102 | 36.6545 | 30.796 | 29.0995 | 26.6972 | 22.6225 |
| 522.94 | 65.1445 | 59.2021 | 52.9801 | 44.686 | 36.2164 | 32.3034 | 26.8168 | 25.2133 | 23.928 |
| 523.97 | 64.3501 | 58.3386 | 51.7139 | 42.86 | 35.8558 | 30.3916 | 28.091 | 23.6931 | 23.8935 |
| 525 | 60.3617 | 56.5846 | 49.8778 | 42.6017 | 35.8046 | 30.1662 | 26.4108 | 24.6479 | 23.3846 |
| 526.02 | 60.6801 | 55.4287 | 48.7616 | 39.4555 | 34.6192 | 28.3433 | 26.4165 | 23.8377 | 19.4298 |
| 527.05 | 59.9455 | 54.5994 | 49.0033 | 40.4292 | 35.7427 | 28.0833 | 25.3696 | 23.7262 | 18.997 |
| 527.94 | 60.5936 | 54.5789 | 48.1615 | 39.7705 | 30.7692 | 29.5662 | 24.23 | 21.7918 | 20.0741 |
| 528.97 | 57.677 | 52.4703 | 46.6128 | 39.0747 | 30.8325 | 26.3878 | 25.1743 | 21.4315 | 17.9984 |
| 530 | 55.2268 | 52.2876 | 44.7278 | 37.8831 | 31.2871 | 26.058 | 23.0219 | 21.6055 | 18.7977 |
| 531.04 | 53.1892 | 50.9091 | 45.6956 | 36.142 | 30.8239 | 26.6437 | 22.8248 | 19.3332 | 17.1045 |
| 531.94 | 54.2669 | 50.2453 | 42.9509 | 34.4188 | 29.7553 | 24.7819 | 22.9724 | 21.0399 | 18.7593 |
| 532.98 | 52.7542 | 47.3592 | 41.461 | 34.3936 | 26.4215 | 24.1584 | 21.1327 | 17.8747 | 18.4472 |
| 534.02 | 50.9418 | 47.3577 | 41.0133 | 33.7101 | 27.969 | 23.5903 | 20.4604 | 18.2119 | 15.8172 |
| 535.07 | 52.3344 | 46.0321 | 38.9715 | 33.623 | 26.7279 | 21.2976 | 20.6693 | 17.348 | 17.3421 |
| 535.97 | 50.6055 | 44.1865 | 38.5686 | 32.5618 | 27.7111 | 23.13 | 21.0547 | 16.6904 | 16.5657 |
| 537.01 | 48.5146 | 43.1739 | 38.0038 | 32.2327 | 26.8959 | 22.3966 | 18.4762 | 17.8504 | 15.1175 |
| 538.05 | 45.4076 | 44.0711 | 37.3419 | 29.9688 | 24.7983 | 22.4444 | 18.7609 | 15.1576 | 15.8666 |
| 538.95 | 45.9866 | 41.2586 | 37.7395 | 30.6445 | 23.753 | 19.2644 | 17.8218 | 15.3816 | 14.4155 |
| 540 | 43.7788 | 40.2288 | 34.5844 | 29.4845 | 24.5656 | 20.8665 | 17.2448 | 16.4389 | 15.3398 |
| 541.02 | 44.4399 | 39.0995 | 34.5405 | 28.1244 | 23.8046 | 19.5365 | 16.744 | 15.6504 | 14.0601 |
| 542.05 | 42.6855 | 38.5069 | 31.9719 | 28.169 | 22.8467 | 19.342 | 15.3754 | 14.4259 | 13.1655 |
| 542.94 | 43.8563 | 37.9548 | 31.9952 | 27.8377 | 20.8943 | 18.96 | 15.2491 | 14.1371 | 12.4951 |
| 543.97 | 40.1338 | 37.6163 | 29.6701 | 25.7732 | 19.8985 | 18.9739 | 16.2162 | 13.1982 | 12.833 |
| 545 | 41.6089 | 35.5589 | 29.9043 | 26.9805 | 20.9598 | 18.0894 | 14.364 | 12.7668 | 11.4808 |
| 546.02 | 38.5396 | 35.2645 | 30.5284 | 24.3949 | 20.3141 | 18.1467 | 14.7537 | 12.6607 | 11.4192 |
| 547.05 | 37.9831 | 33.7988 | 31.2438 | 26.1617 | 20.7809 | 16.6732 | 14.3521 | 12.9285 | 11.4353 |
| 547.94 | 37.2051 | 33.2925 | 29.6524 | 24.5221 | 19.7275 | 15.873 | 15.2626 | 12.9006 | 11.7158 |
| 548.97 | 38.6694 | 32.5 | 28.0106 | 23.5629 | 18.4995 | 15.3249 | 14.0387 | 11.9391 | 10.3637 |

**Fluorescence spectra of 2',6-dichloro-1*H*,3'*H*-2,5'-bibenzo[*d*]imidazole (XV – MB_2_Cl) binding to dsDNA**

MB2Cl

Intensity, rel. units

| Wavelength, nm | 0 | 9 | 53 | 140 | 341 | 525 |
| --- | --- | --- | --- | --- | --- | --- |
| 304.92 | 194.953 | 161.325 | 151.151 | 116.257 | 79.163 | 81.6168 |
| 306.92 | 151.625 | 156.376 | 133.764 | 113.115 | 84.1699 | 76.2493 |
| 308.92 | 138.514 | 123.783 | 146.48 | 112.035 | 69.949 | 67.3795 |
| 310.92 | 138.817 | 127.906 | 128.573 | 104.638 | 68.0179 | 68.2666 |
| 312.92 | 121.324 | 94.0694 | 118.167 | 102.003 | 74.002 | 59.0894 |
| 314.92 | 127.187 | 111.39 | 116.292 | 82.3286 | 68.4838 | 63.3074 |
| 316.92 | 105.047 | 109.46 | 107.503 | 84.7899 | 59.8662 | 52.941 |
| 318.92 | 112.809 | 110.776 | 103.669 | 79.0773 | 71.3429 | 59.8333 |
| 320.92 | 128.595 | 107.284 | 117.087 | 85.7873 | 73.0712 | 71.6094 |
| 322.92 | 145.495 | 156.988 | 129.083 | 124.873 | 99.8262 | 97.8539 |
| 324.92 | 170.629 | 172.072 | 181.702 | 165.946 | 126.518 | 118.361 |
| 326.92 | 213.013 | 178.448 | 209.802 | 183.771 | 154.28 | 131.983 |
| 328.92 | 193.586 | 219.383 | 197.913 | 183.005 | 162.395 | 150.708 |
| 330.92 | 194.3 | 189.702 | 170.638 | 158.331 | 151.764 | 126.091 |
| 332.92 | 168.217 | 164.56 | 163.537 | 135.484 | 143.189 | 104.293 |
| 334.92 | 159.246 | 153.908 | 141.522 | 147.813 | 123.033 | 96.7855 |
| 336.92 | 168.068 | 156.73 | 162.589 | 148.691 | 125.177 | 97.3007 |
| 338.92 | 178.152 | 167.584 | 155.221 | 136.113 | 136.173 | 109.63 |
| 340.9 | 169.346 | 159.878 | 174.406 | 145.856 | 139.428 | 120.533 |
| 342.87 | 181.345 | 166.12 | 178.277 | 157.699 | 128.265 | 101.909 |
| 344.84 | 177.994 | 188.258 | 186.108 | 168.959 | 131.052 | 118.632 |
| 346.96 | 192.615 | 180.938 | 183.639 | 165.995 | 145.77 | 121.135 |
| 348.93 | 195.51 | 183.647 | 165.765 | 151.626 | 135.435 | 123.897 |
| 350.92 | 176.811 | 174.225 | 178.779 | 169.081 | 161.559 | 130.686 |
| 352.92 | 197.099 | 172.259 | 178.19 | 182.029 | 146.527 | 132.424 |
| 354.92 | 197.861 | 191.97 | 188.135 | 170.056 | 162.298 | 116.506 |
| 356.92 | 179.7 | 193.126 | 192.66 | 164.554 | 157.037 | 129.473 |
| 358.92 | 190.929 | 196.173 | 198.348 | 165.626 | 132.544 | 136.605 |
| 360.9 | 213.563 | 210.531 | 188.057 | 167.424 | 142.806 | 125.333 |
| 362.87 | 191.024 | 185.258 | 192.12 | 162.293 | 139.339 | 139.671 |
| 364.84 | 178.13 | 188.802 | 171.871 | 168.021 | 153.245 | 121.178 |
| 366.96 | 176.586 | 178.172 | 163.408 | 165.611 | 140.644 | 113.953 |
| 368.93 | 183.7 | 169.991 | 177.334 | 162.768 | 129.171 | 118.905 |
| 370.92 | 182.119 | 161.528 | 154.352 | 155.637 | 127.959 | 108.346 |
| 372.92 | 165.397 | 154.643 | 165.693 | 144.604 | 117.756 | 111.751 |
| 374.92 | 151.554 | 146.09 | 144.64 | 135.857 | 120.968 | 101.869 |
| 376.92 | 148.734 | 147.215 | 144.204 | 132.794 | 132.404 | 107.906 |
| 378.92 | 135.119 | 136.279 | 145.81 | 143.911 | 119.654 | 110.383 |
| 380.9 | 137.218 | 133.598 | 131.319 | 131.87 | 101.773 | 96.6309 |
| 382.87 | 136.136 | 120.982 | 132.971 | 110.51 | 113.763 | 94.3631 |
| 384.84 | 104.245 | 121.747 | 132.772 | 110.864 | 95.4928 | 98.251 |
| 386.96 | 107.265 | 104.433 | 111.869 | 101.746 | 105.672 | 87.8517 |
| 388.93 | 98.88 | 112.781 | 103.987 | 111.462 | 96.6029 | 82.1127 |
| 390.9 | 89.107 | 112.515 | 105.224 | 96.6554 | 86.8205 | 76.9889 |
| 392.87 | 97.2851 | 113.456 | 107.548 | 90.9477 | 86.779 | 80.2139 |
| 394.84 | 98.521 | 100.366 | 88.2611 | 92.8544 | 85.2518 | 78.25 |
| 396.96 | 86.484 | 92.458 | 94.9608 | 94.765 | 72.6488 | 75.0263 |
| 398.93 | 89.7777 | 86.4517 | 84.3119 | 84.9028 | 78.7059 | 70.0931 |
| 400.92 | 75.2292 | 76.8671 | 80.2362 | 82.974 | 77.5047 | 74.8607 |
| 402.92 | 79.9282 | 74.9337 | 78.7802 | 78.8602 | 75.0429 | 64.9254 |
| 404.92 | 70.1748 | 69.6357 | 71.7383 | 64.3692 | 65.0584 | 69.3499 |
| 406.92 | 63.6787 | 74.1217 | 62.112 | 63.7214 | 60.5667 | 50.6924 |
| 408.92 | 58.5974 | 62.7187 | 62.7635 | 65.9567 | 63.8149 | 52.6218 |
| 410.9 | 64.6436 | 67.4602 | 68.8619 | 65.5993 | 68.0245 | 62.7236 |
| 412.87 | 49.6335 | 67.3384 | 67.474 | 74.3251 | 57.2871 | 59.6329 |
| 414.84 | 60.6965 | 53.4952 | 60.8943 | 62.6292 | 60.5565 | 56.4685 |
| 416.96 | 61.1954 | 64.8359 | 48.7081 | 52.9014 | 57.311 | 56.2696 |
| 418.93 | 45.0157 | 52.8999 | 53.1521 | 53.5567 | 60.241 | 53.8762 |
| 420.9 | 49.8401 | 44.0843 | 58.9516 | 55.0455 | 56.2471 | 49.544 |
| 422.87 | 50.8898 | 52.2228 | 39.9683 | 50.5031 | 53.8792 | 56.3612 |
| 424.84 | 41.1477 | 41.3589 | 55.096 | 43.7822 | 50.4466 | 43.1345 |
| 426.96 | 36.1685 | 43.0855 | 47.3324 | 40.506 | 43.1787 | 45.388 |
| 428.93 | 41.5635 | 36.6248 | 43.0071 | 44.6236 | 39.4559 | 33.55 |
| 430.89 | 38.0452 | 34.3734 | 38.0375 | 39.3287 | 36.2668 | 39.8209 |
| 432.98 | 33.9979 | 42.0454 | 46.953 | 34.8689 | 39.393 | 35.3702 |
| 434.92 | 31.4134 | 35.5435 | 38.8177 | 33.9389 | 35.2222 | 33.6634 |
| 436.86 | 20.0322 | 31.8647 | 28.492 | 36.7863 | 32.5641 | 41.445 |
| 438.95 | 30.839 | 34.0361 | 32.2174 | 36.1872 | 33.8788 | 28.2663 |
| 440.9 | 21.9789 | 26.7925 | 34.164 | 32.0076 | 26.4932 | 36.5607 |
| 442.87 | 28.184 | 29.1866 | 29.1971 | 27.6401 | 33.0285 | 32.9202 |
| 444.84 | 24.5318 | 24.095 | 24.2936 | 28.1627 | 28.4942 | 28.3442 |
| 446.96 | 22.2301 | 29.0056 | 26.5913 | 24.3662 | 28.6385 | 28.8848 |
| 448.93 | 22.7816 | 23.8635 | 24.1023 | 20.5068 | 32.6492 | 28.1151 |
| 450.89 | 24.646 | 20.5141 | 30.9138 | 23.087 | 27.3092 | 21.3113 |
| 452.98 | 17.2648 | 24.0398 | 19.9873 | 22.4393 | 25.4578 | 25.2184 |
| 454.92 | 18.4715 | 17.7001 | 20.4609 | 25.2488 | 30.746 | 25.2208 |
| 456.86 | 23.392 | 22.3932 | 23.8717 | 21.8543 | 19.7474 | 24.5121 |
| 458.95 | 27.2565 | 23.6854 | 19.2744 | 20.693 | 26.7082 | 24.998 |
| 460.9 | 16.3124 | 19.2895 | 27.2446 | 22.7465 | 26.5275 | 22.8301 |
| 462.87 | 18.4304 | 24.8071 | 21.2854 | 17.1319 | 22.4862 | 24.6437 |
| 464.84 | 15.5988 | 19.8794 | 18.9659 | 22.9307 | 25.7545 | 25.1559 |
| 466.96 | 16.8575 | 14.3651 | 20.7017 | 14.2069 | 18.5556 | 22.557 |
| 468.93 | 18.1298 | 19.5644 | 12.4966 | 17.2106 | 21.4197 | 19.1208 |
| 470.89 | 12.9129 | 17.2927 | 19.039 | 18.8679 | 20.4851 | 15.2887 |
| 472.98 | 18.5716 | 18.4756 | 18.1182 | 16.611 | 27.1144 | 20.2421 |
| 474.92 | 19.2551 | 17.718 | 25.9295 | 17.4342 | 16.6839 | 19.4422 |
| 476.86 | 17.1694 | 12.5807 | 12.8926 | 16.5858 | 19.1078 | 21.7049 |
| 478.95 | 14.315 | 15.0376 | 14.3701 | 11.7713 | 17.5332 | 19.9744 |
| 480.89 | 19.1842 | 15.2177 | 15.4698 | 15.7105 | 14.7489 | 19.0546 |
| 482.98 | 13.5552 | 17.5626 | 18.7127 | 13.0282 | 15.6638 | 23.0504 |
| 484.92 | 11.1493 | 15.5452 | 14.5852 | 15.8673 | 19.2454 | 15.1905 |
| 486.86 | 15.4157 | 8.62845 | 12.2324 | 14.19 | 17.1544 | 18.7827 |
| 488.95 | 12.5571 | 15.6369 | 18.7973 | 16.8077 | 16.9323 | 13.6729 |
| 490.89 | 9.6225 | 13.8452 | 16.6949 | 14.9683 | 13.9552 | 16.8762 |
| 492.98 | 9.51258 | 11.6331 | 14.194 | 13.2109 | 14.3951 | 19.399 |
| 494.92 | 20.7371 | 13.4846 | 11.6879 | 13.3994 | 14.2955 | 18.2356 |
| 496.86 | 14.9573 | 15.9137 | 16.1461 | 13.1218 | 17.267 | 17.2377 |
| 498.95 | 12.4727 | 11.8334 | 18.253 | 11.5165 | 15.2497 | 15.3506 |
| 500.89 | 11.8415 | 12.142 | 11.5082 | 15.5958 | 12.6062 | 14.0422 |
